# Protein lysine methylation is involved in modulating the response of sensitive and tolerant Arabidopsis species to cadmium stress

**DOI:** 10.1101/652651

**Authors:** Nelson B.C. Serre, Manon Sarthou, Océane Gigarel, Sylvie Figuet, Massimiliano Corso, Justine Choulet, Valérie Rofidal, Claude Alban, Véronique Santoni, Jacques Bourguignon, Nathalie Verbruggen, Stéphane Ravanel

## Abstract

The mechanisms underlying the response and adaptation of plants to excess of trace elements are not fully described. Here, we analyzed the importance of protein lysine methylation for plants to cope with cadmium. We analyzed the effect of cadmium on lysine-methylated proteins and protein lysine methyltransferases (KMTs) in two cadmium-sensitive species, *Arabidopsis thaliana* and *A. lyrata*, and in three populations of *A. halleri* with contrasting cadmium accumulation and tolerance traits. We showed that some proteins are differentially methylated at lysine residues in response to Cd and that a few genes coding KMTs is regulated by cadmium. Also, we showed that nine out of 23 *A. thaliana* mutants interrupted in *KMT* genes have a tolerance to cadmium that is significantly different from that of wild-type seedlings. We further characterized two of these mutants, one was knocked-out in the calmodulin lysine methyltransferase gene and displayed increased tolerance to cadmium, the other was interrupted in a *KMT* gene of unknown function and showed a decreased capacity to cope with cadmium. Together, our results showed that lysine methylation of non-histone proteins is impacted by cadmium and that several methylation events are important for modulating the response of Arabidopsis plants to cadmium stress.

## INTRODUCTION

As sessile organisms, land plants must deal with fluctuating levels of essential and non-essential trace elements in soils. Some plant species have the ability to colonize soils contaminated by toxic levels of metals and display remarkable leaf metal accumulation without visible toxicity symptoms. Understanding tolerance and accumulation of metals in these species, referred to as hyperaccumulators, offers the unique opportunity to uncover key mechanisms governing metal homeostasis and adaptation to challenging environments (for reviews, see Verbruggen *et al*., 2009; Kramer, 2010). *Arabidopsis halleri*, a close relative of *A. thaliana* and *A. lyrata*, is a model species for studying tolerance and accumulation of cadmium (Cd), one of the most toxic metal for living organisms (for reviews, see Kramer, 2010; DalCorso *et al*., 2013; Verbruggen *et al*., 2013; Moulis *et al*., 2014). While Cd and zinc tolerance seems to be constitutive in *A. halleri*, populations originating from different genetic units and from metallicolous or non-metallicolous soils display important variability in terms of Cd accumulation (Meyer *et al*., 2015; Stein *et al*., 2017; Corso *et al*., 2018; Schvartzman *et al*., 2018). This intraspecific variability suggests adaptation at the local scale and, possibly, the involvement of different molecular mechanisms to account for metal accumulation and tolerance traits.

In the last 15 years, the combination of genetic, ‘omics’ and functional approaches in both tolerant and non-tolerant species have contributed considerably to the understanding of Cd toxicity, tolerance and accumulation. Key mechanisms involved in metal uptake, translocation, chelation with ligands, vacuolar sequestration, and cell signaling have been characterized (for reviews, see Villiers *et al*., 2011; DalCorso *et al*., 2013; Clemens *et al*., 2013; Clemens & Ma, 2016). The coordination of these processes is accomplished through multilevel regulatory mechanisms, including the epigenetic, transcriptional, and post-translational levels (Gallego *et al*., 2012; Haak *et al*., 2017). Despite recent progress, the role of post-translational modifications (PTMs) in the response and adaptation of plants to Cd and other trace elements is still poorly documented. Given their pivotal role in the regulation of many cellular processes, it is anticipated that PTMs are important for plants to cope with biotic and abiotic stresses (Dahan *et al*., 2011; Kosava *et al*., 2011). To date, the most studied modification in the field of metal stress is protein phosphorylation. It has been shown that Cd poisoning in *A. thaliana* or *Medicago sativa* is accompanied by the activation of several mitogen-activated protein kinases (Jonak *et al*., 2004; Liu *et al*., 2010) and SNF1-related protein kinases (Kulik *et al*., 2012). Also, it has been reported that Cd stress induces the hyper-phosphorylation of the eukaryotic translation initiation factor eIF2a (Sormani *et al*., 2011) and the plasma membrane H^+^-ATPase (Janicka-Russak *et al*., 2012). Metal stress is also responsible for changes in the pattern of PTMs on tubulin (Gzyl *et al*., 2015). Last, it is worth noting that the production of reactive oxygen species triggered by Cd is responsible for the carbonylation of proteins, an irreversible oxidative modification leading to the degradation of damaged proteins by the proteasome. Several studies indicated that plants treated with Cd display an increased oxidation of proteins (e.g. Romero-Puertas *et al*., 2002; Pena *et al*., 2012) and an upregulation of the 20S proteasome proteolytic pathway (e.g. Pena *et al*., 2006; Sarry *et al*., 2006; Polge *et al*., 2009). Protein methylation is a very diverse PTM acting on different protein residues. It is widespread in eukaryotic proteomes, modifying both histones and non-histone proteins, and contributes to the fine-tuned regulation of protein function. The modification of the lysine (Lys) side chain of proteins is among the most frequent methylation event (Lanouette *et al*., 2014; Falnes *et al*., 2016). It is catalyzed by two structurally different classes of protein Lys methyltransferases (KMTs), the SET domain-containing group (SDG) and the seven-beta-strand (SBS) superfamily, which are able to add one to three methyl groups to specific Lys residues in proteins (Serre *et al*., 2018). In plants, these enzymes have been shown to methylate histones and non-histone proteins involved in all aspects of cell biology (e.g. transcription, protein synthesis, metabolism). Protein Lys methylation can be reversible by the action of demethylases. Demethylases acting on non-histone Lys-methylated proteins have not been yet reported in plants (Serre *et al*., 2018).

The identification of the enzyme/substrate (KMT/protein) relationship is a critical step towards understanding the role of protein methylation (Lanouette *et al*., 2014; Falnes *et al*., 2016; Serre *et al*., 2018). As such, inactivating the genes coding KMTs is the best way to analyze the functional outcomes of methylation *in vivo*. Despite recent progress, the role of non-histone protein Lys methylation in regulating plant cellular functions is still limited (Serre *et al*., 2018). In particular, no information is available about the role of protein methylation in the response and adaptation of plants to metal stress. The present work is based on the assumption that this PTM could be important for plants to efficiently address stress situations induced by metals. This hypothesis is supported by the abundance and diversity of Lys-methylated proteins, possibly targeting components of metal transport, signaling pathways or detoxification machineries, and the recognized role of this modification in the regulation of protein function (Lanouette *et al*., 2014; Falnes *et al*., 2016). To test this hypothesis, we analyzed the expression of the two main players involved in protein Lys methylation, i.e. Lys-methylated proteins and genes coding KMTs, in three Arabidopsis species challenged with Cd. We used *A. thaliana* and *A. lyrata* non-tolerant plants and three populations of *A. halleri* from different genetic units and showing contrasting tolerance and accumulation of Cd (Meyer *et al*., 2015; Corso *et al*., 2018; Schvartzman *et al*., 2018). First, we showed that some non-histone proteins are differentially methylated at Lys residues in response to Cd and we identified one of these proteins by MS/MS. Second, we showed that Cd stress has limited impact on the transcriptional regulation of *KMT* genes. Third, using a root growth inhibition assay with *A. thaliana* mutants disrupted in genes coding KMTs, we showed that nine out of 23 mutants have a tolerance to Cd that is different from that of wild-type seedlings. Last, we characterized two of these mutants that are either more tolerant or more sensitive to Cd. Together, our results showed that Cd triggers changes in the expression of a few Lys-methylated proteins and *KMT* genes, and that several Lys-methylating enzymes are important for modulating the response of Arabidopsis plants to Cd stress.

## MATERIALS AND METHODS

### Plant material and growth conditions

*Arabidopsis thaliana* ecotype Columbia (Col-0), *Arabidopsis lyrata* ssp. *petraea* (Linnaeus) O’Kane & Al-Shehbaz, and the *Arabidopsis halleri* ssp. *halleri* (Linnaeus) O’Kane & Al-Shehbaz populations from the metallicolous soils located in Auby (North of France, AU population), Val del Riso (North of Italy, I16 population), and Bukowno (South of Poland, PL22 population) (Meyer *et al*., 2015) were grown in hydroponic conditions. The standard control medium (CM) was composed of 0.88 mM K_2_SO_4_, 2 mM Ca(NO_3_)_2_, 1 mM MgSO_4_, 0.25 mM KH_2_PO_4_, 10 µM H_3_BO_3_, 0.1 µM CuSO_4_, 0.6 µM MnSO_4_, 0.01 µM (NH_4_)_6_Mo_7_O_24_, 10 µM ZnSO_4_, 10 µM NaCl, 20 µM Fe-EDTA, and 0.25 mM MES, pH 5.8 (Meyer *et al*., 2015). Plants were grown at 21°C, 70% air humidity, under short day conditions (8 hours of light per day) with a light intensity of 80 µmol of photons m^-2^ s^-1^. After four weeks of growth plants were maintained in CM or challenged with 0.2 to 5 µM CdSO_4_ in CM for 7 to 10 days. At the end of the treatment, roots and leaves were harvested separately from each individual, washed twice in distilled water, dried with absorbent paper, and frozen in liquid nitrogen.

### Protein extraction and immunoblotting

Proteins from Arabidopsis tissues were extracted by grinding frozen-powdered tissues in 50 mM Tris-HCl, pH 8.0, 10% (v/v) glycerol, and protease inhibitors (Roche Applied Science). Samples were centrifuged at 16,000 x g for 20 min and the supernatant used as a source of soluble proteins. Pellets were suspended in the extraction buffer supplemented with 2% (w/v) SDS, incubated for 15 min at room temperature, and centrifuged as before to recover solubilized membrane proteins. Proteins were resolved by SDS-PAGE, transferred to nitrocellulose membrane, and probed with an antibody against trimethyl-Lys (abcam 76118). Membranes were also probed with antibodies against fructose bisphosphate aldolase (Mininno *et al*., 2012) and the beta subunit of ATPase (Agrisera, AS03 030) for the normalization of protein loading. Protein detection was achieved using the ECL Plus™ Western Blotting detection reagents.

### Identification of Lys-methylated proteins by mass spectrometry

Sample preparation - Soluble proteins from root and leaf samples were resolved by SDS-PAGE and bands were cut in the range 25-30 kDa. After washing with water and then 25 mM NH_4_HCO_3_ gel bands were destained twice with 1 mL of CH_3_CN and dried at room temperature. Disulphide bridges were reduced using 10 mM dithiothreitol at 56°C for 45 min and cysteine were alkylated using 55 mM iodoacetamide for 30 min in darkness. Gel bands were washed twice with 50% (v/v) CH_3_CN in 25 mM NH_4_HCO_3_, then dehydrated with CH_3_CN, and finally dried at room temperature. In-gel protein digestion was performed overnight at 37°C with trypsin (Sequencing Grade Modified Trypsin, Promega, Madison, USA) at a final concentration of 0.005 µg/µL. Peptides were extracted twice using 2% (v/v) formic acid in 80% (v/v) CH_3_CN, dried, and then suspended in 20 µL of 2% (v/v) formic acid before LC-MS/MS analysis.

Mass-spectrometry analysis - LC-MS/MS experiments were done using an UltiMate 3000 RSLCnano system interfaced online with a nano easy ion source and a Q Exactive Plus Orbitrap mass spectrometer (Thermo Fisher Scientific Inc, Waltham, MA, USA) operating in a Data Dependent Acquisition (DDA) mode. Peptides were separated by reverse-phase chromatography (PepMap C18, 2 µm particle size, 100 Å pore size, 75 µm i.d. x 50 cm length, Thermo Fisher Scientific) at a flow rate of 300 nL/min. Loading buffer (solvent A) was 0.1% (v/v) trifluoroacetic acid in water and elution buffer (solvent B) was 0.1% (v/v) trifluoroacetic acid in 80% (v/v) acetonitrile. The three-step gradient employed was 4 to 25% of solvent B in 103 min, then 25 to 40% of solvent B from 103 to 123 min, finally 40 to 90% of solvent B from 123 to 125 min. Peptides were transferred to the gaseous phase with positive ion electrospray ionization at 1.7 kV. In DDA, the top 10 precursors were acquired between 375 and 1500 m/z with a 2 Thomson selection window, dynamic exclusion of 40 s, normalized collision energy of 27 and resolutions of 70,000 for MS and 17,500 for MS2. Spectra were recorded with the Xcalibur software (4.0.27.19) (Thermo Fisher Scientific).

Identification of methylpeptides - Mass spectrometry data were processed using the Proteome Discoverer software (version 1.4.0.288, Thermo Fisher Scientific) and a local search engine (Mascot, version 2.4.1, Matrix Science). Data from A. thaliana samples were searched against the TAIR (2011) non-redundant database containing 35,387 sequences with the following parameters: trypsin as enzyme, 3 missed cleavages allowed, carbamidomethylation of cysteine as a fixed modification, and mono-, di-, tri-methylation of Lys, acetylation of Lys, N-terminal acetylation of the protein, deamidation of asparagine and glutamine, N-terminal pyromutamylation of glutamine and glutamate, and oxidation of methionine as variable modifications. Mass tolerance was set at 10 ppm on full scans and 0.02 Da for fragment ions. Proteins were validated once they contained at least two peptides with a p-value <0.05. Two additional filters were used to improve the identification of trimethylated Lys peptides: 1/ selection of peptides with Mascot score ≥30, 2/ discrimination of Lys trimethylation (mass shift of 42.04695) and Lys acetylation (mass shift of 42.01056) using a mass tolerance at 2 ppm. Ambiguous peptides were eliminated and spectra of interest were checked manually to confirm their sequence and the nature of modifications. Similar parameters were used to identify trimethylated Lys peptides in samples from *A. halleri* and *A. lyrata* but MS data were searched using a local database built using the *A. lyrata* genome resources (Alyrata_384_v2.1 from the Joint Genome Institute) (Hu *et al*., 2011; Rawat *et al*., 2015).

### Screening of *A. thaliana* mutants in protein Lys methyltransferase genes

Seeds of the T-DNA insertion lines in 23 *KMT* genes were obtained from the European Nottingham Arabidopsis Stock Centre. Mutants were genotyped by PCR using gene- and T-DNA-specific primers (Table S1). Amplicons were sequenced to map the insertion sites.

Seeds of Col-0 and homozygous *KMT* mutants were surface sterilized and sown onto Petri dishes containing half-strength Murashige and Skoog (MS/2) medium with 0.8% (w/v) agar. After two days of stratification at 4°C, plates were transferred to a growth chamber for four days (21°C, 70% air humidity, 18 hours of light per day, 80 µmol of photons m^-2^ s^-1^). Twenty seedlings per genotype were then transferred to square Petri dishes containing either MS/2 medium or MS/2 medium with 20 µM CdSO_4_. Plates were oriented vertically in the growth chamber and scanned (GS-800 scanner, BioRad) after 0, 3, 6, 8 and 10 days of treatment. The root length of each seedling was measured with the ImageJ software. Root length at day 8 and root elongation rate (in cm.day^-1^) between days 3 and 8 were used as primary criteria to monitor the inhibitory effect of Cd. Also, the tolerance index (TI) to Cd was calculated for each line by dividing the primary parameter (length or elongation rate) measured in the presence of Cd by the one measured in the control condition. One hundred TI were calculated for each line by random sampling of one value in the Cd and control conditions, respectively, with replacement at each draw.

### Gene expression data mining

The expression of genes coding KMTs in *A. thaliana* seedlings challenged with Cd was analyzed using data from the literature. First, we used our genome-wide CATMA microarray analysis of roots and shoots from 4-week-old *A. thaliana* plants exposed to 5 or 50 µM Cd for 2, 6 or 30 hours (GSE10675) (Herbette *et al.,* 2006). Next, we collected four datasets from the Gene Expression Omnibus database. They correspond to i/ 2-week-old seedlings grown in MS/2 agar plates and treated with 70 µM Cd for 2 hours (GSE90701) (Khare *et al.,* 2016); ii/ roots from 5-week-old plants grown in hydroponics and treated for 7 days with 1 µM Cd (GSE94314) (Fischer *et al.,* 2017); iii/ roots from 3-week-old seedlings grown in hydroponics and challenged with 200 µM Cd for 6 hours (GSE22114) (Li *et al.,* 2010); and iv/ 7-day-old seedlings grown in MS/2 agar plates and treated for 6 hours with 200 µM Cd (GSE35869) (Jobe *et al.,* 2012). Also, we analyzed curated data from the comparative transcriptomic analysis of *A. thaliana* and *A. halleri* plants challenged with 10 µM or 25 µM Cd, respectively, in hydroponics for 2 hours (Weber *et al.,* 2006). Relative expression levels of *KMT* genes were retrieved from the different experiments, the ratio between Cd and control conditions were calculated from pairwise comparisons, and a non-parametric Student t-test was performed on log2 ratio to determine differentially expressed genes (DEGs, with p-value<0.05) and the threshold was set at 2-fold (-1≤ log2 fold change ≥1). Finally, we used the RNA-seq data we obtained to analyze the tolerance strategies to Cd of two metallicolous populations of *A. halleri* (BioProject PRJNA388549) (Corso *et al.,* 2018). In this experiment, gene expression in the I16 and PL22 populations was analyzed after 10 days of treatment with 5 µM Cd in hydroponics. For the identification of DEGs, we selected genes with more than 10 read counts in any of the triplicate, applied a non-parametric t-test (p-value<0.05) in pairwise comparisons, and used a 1.4-fold change threshold value (-0.5≤ log2 fold change ≥0.5).

### Determination of Cd by inductively coupled plasma mass spectrometry (ICP-MS)

Plant samples were dehydrated at 90°C, weighed for data normalization, and digested at 90°C for 4 hours in 65% (w/w) ultrapure HNO_3_. Mineralized samples were diluted in 0.5% (v/v) HNO_3_ and analyzed using an iCAP RQ quadrupole mass instrument (Thermo Fisher Scientific GmbH, Germany). ^111^Cadmium concentration was determined using a standard curve and corrected using an internal standard solution of ^103^Rhodium added online.

### Statistical analysis

Non-parametric statistical analysis was performed on our datasets, which typically contain small sample sizes (n≤20) and do not meet the assumptions of parametric tests (normal distribution and homogeneity of variance, as determined using the Shapiro-Wilk and Fisher tests, respectively). Multiple non-parametric comparisons were performed with the Dunnett’s many-to-one test using the nparcomp package (Konietschke *et al*., 2015) and the R computing environment. The Fischer’s approximation method was used and the confidence level was set at 95%.

## RESULTS

### Analysis of the patterns of Lys-methylated proteins in sensitive and tolerant Arabidopsis species exposed to Cd

We analyzed the effect of Cd stress on the pattern of Lys-methylated proteins in roots and leaves of three Arabidopsis species showing contrasting Cd tolerance and accumulation. The analysis was done by immunoblotting and focused on Lys trimethylation on proteins other than histones because antibodies against mono- and dimethyl-Lys are less sensitive and specific than anti trimethyl-Lys antibodies (Alban *et al*., 2014). Moreover, the procedure used for protein isolation was not appropriate for the extraction of histones, which requires acidic or high salt conditions (Shechter *et al*., 2007). We used the Cd-sensitive species *A. thaliana* (ecotype Columbia, Col-0) and *A. lyrata* ssp. *petraea*, and the *A. halleri* species (Auby population) that displays Cd hypertolerance and hyperaccumulation traits (Meyer *et al*., 2015). Plants were grown hydroponically for five weeks in a standard culture medium and then challenged with 5 µM CdSO_4_ for nine days. In these conditions, the symptoms of Cd toxicity (growth inhibition, chlorosis, and inhibition of photosynthesis) were visible for *A. thaliana* and *A. lyrata* plants, but not for *A. halleri* (Figure S1). The patterns of Lys trimethylated proteins were complex with many polypeptides detected in root extracts and in leaf soluble extracts (Figure 1), illustrating the wide array of targets of Lys methylation. The analysis was less informative for leaf membrane proteins with only a few and diffuse bands detected. A careful examination of the trimethyl-Lys signals indicated several changes in the expression patterns of methylated proteins between species or between control and Cd-treated plants (Figure 1). For example, a Lys-trimethylated protein of 43-45 kDa was strongly labeled in *A. lyrata* leaf soluble extracts, regardless of growth conditions, but was not detectable in *A. thaliana* and *A. halleri* extracts. The most obvious example regarding the effect of Cd was a doublet of proteins at about 26-28 kDa in leaf soluble extracts. This doublet was constitutive in *A. halleri,* i.e. present in both culture conditions, detected in *A. lyrata* treated with Cd, but not observed in *A. lyrata* in control conditions nor in *A. thaliana* with or without Cd treatment (Figure 1a). Noteworthy, a doublet of proteins with a similar migration behavior was detected with a strong and constant immunostaining in root soluble extracts from the three Arabidopsis species in both culture conditions (Figure 1c). Similar western blot analyses were performed with two other populations of *A. halleri,* I16 and PL22, that are hypertolerant to Cd (Meyer *et al*., 2015). The patterns of Lys-trimethylated proteins in I16 and PL22 were similar to those observed for AU, and notably the doublet of proteins at 26-28 kDa in leaf soluble extracts was detected in control and stress conditions (Figure S2). Together, these results indicate that Cd triggers changes in the steady-state level of some Lys-methylated proteins, with contrasting patterns depending on the Arabidopsis species and possibly their ability to tolerate and accumulate Cd.

**Figure 1:**
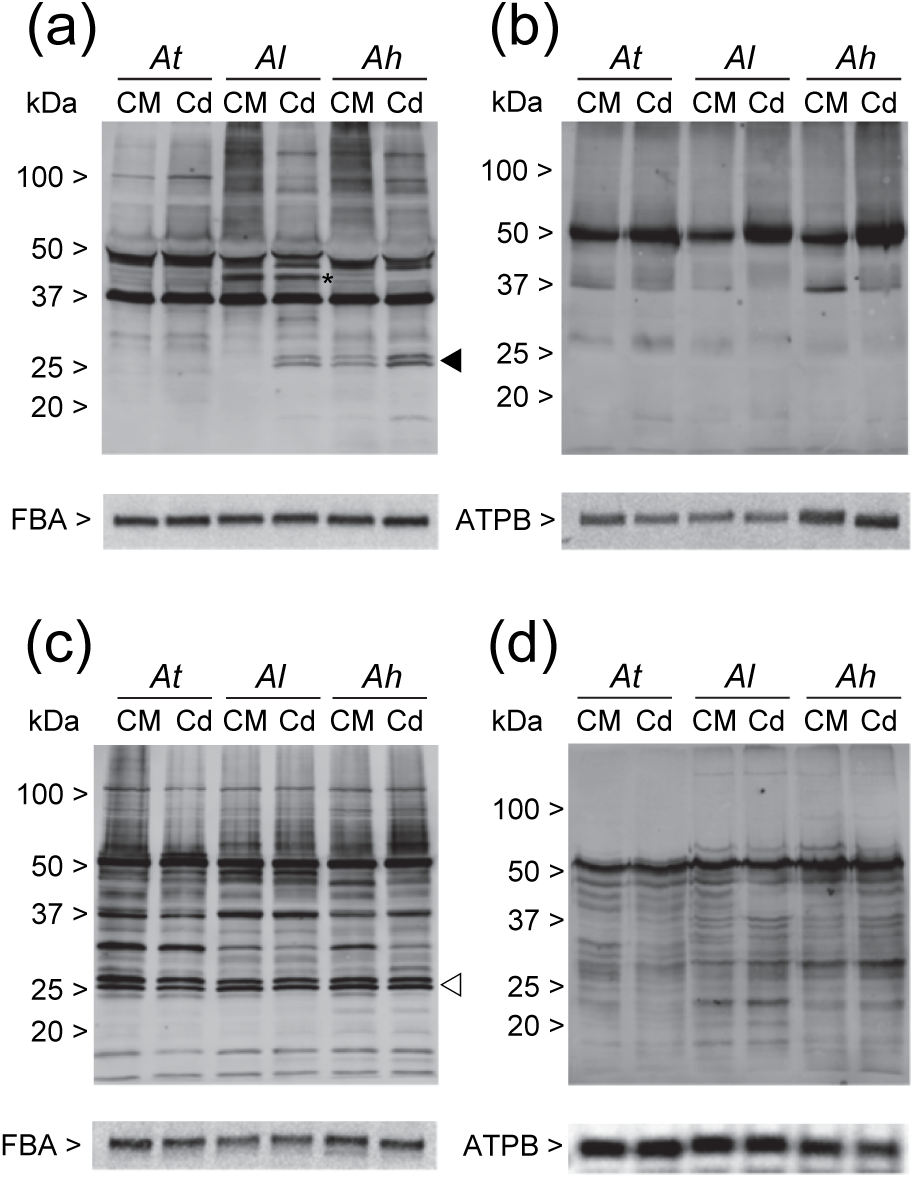
Immunodetection of Lys-trimethylated proteins in roots and leaves from Arabidopsis plants challenged with Cd. Plants grown in hydropony were maintained in control medium (CM) or challenged with 5 µM CdSO_4_ for 9 days. Soluble and membrane proteins (30 µg per lane) from root and leaf tissues were analyzed by western blot using antibodies specific to trimethyl-Lys (ab76118, abcam). (a) - Leaf soluble proteins. (b) - Leaf membrane proteins. (c) - Root soluble proteins. (d) - Root membrane proteins. The asterisk indicates a protein that is labeled specifically in leaf soluble extracts from *A. lyrata*. The triangles indicate protein doublets at 26-28 kDa that is methylated in all root samples (⊳) and follows a species- and/or condition-dependent immunolabeling in leaves (◄). *At*, *A. thaliana*; *Al*, *A. lyrata*; *Ah*, *A. halleri* (AU population). Hybridizations with antibodies against fructose 1,6-bisphosphate aldolase (FBA) and the beta subunit of ATPase (ATPB) have been used as loading controls for soluble and membrane fractions, respectively.

### Identification of Lys-methylated proteins related to Cd stress in Arabidopsis

We used protein tandem mass spectrometry (MS/MS) to identify Lys-methylated proteins whose expression is modulated by Cd. The identification of Lys-methylated peptides by MS/MS is still challenging for many reasons (Wang *et al*., 2017), including the low abundance and/or low methylation level of targets and the high false discovery rates for methylpeptides identification due to amino acid substitutions that are isobaric with methylation events (Ong *et al*., 2004; Hart-Smith *et al*., 2016). To address this challenge we focused on the identification of the abundant doublet of trimethylated proteins at 26-28 kDa for which the expression pattern was potentially interesting regarding Cd stress (Figure 1). We used a filtering procedure adapted from Alban *et al*. (2014) to identify Lys-trimethylated peptides with high confidence. Also, MS/MS data from *A. lyrata* and *A. halleri* were searched against a database built from the *A. lyrata* genome, and not against the *A. thaliana* genome, to improve the identification of Lys-methylated peptides. Using this procedure, we were able to identify Lys-trimethylated peptides belonging to nine proteins in the gel bands of interest in the range 25-30 kDa (Table 1, Table S2, and Figure S3).

**Table 1:**
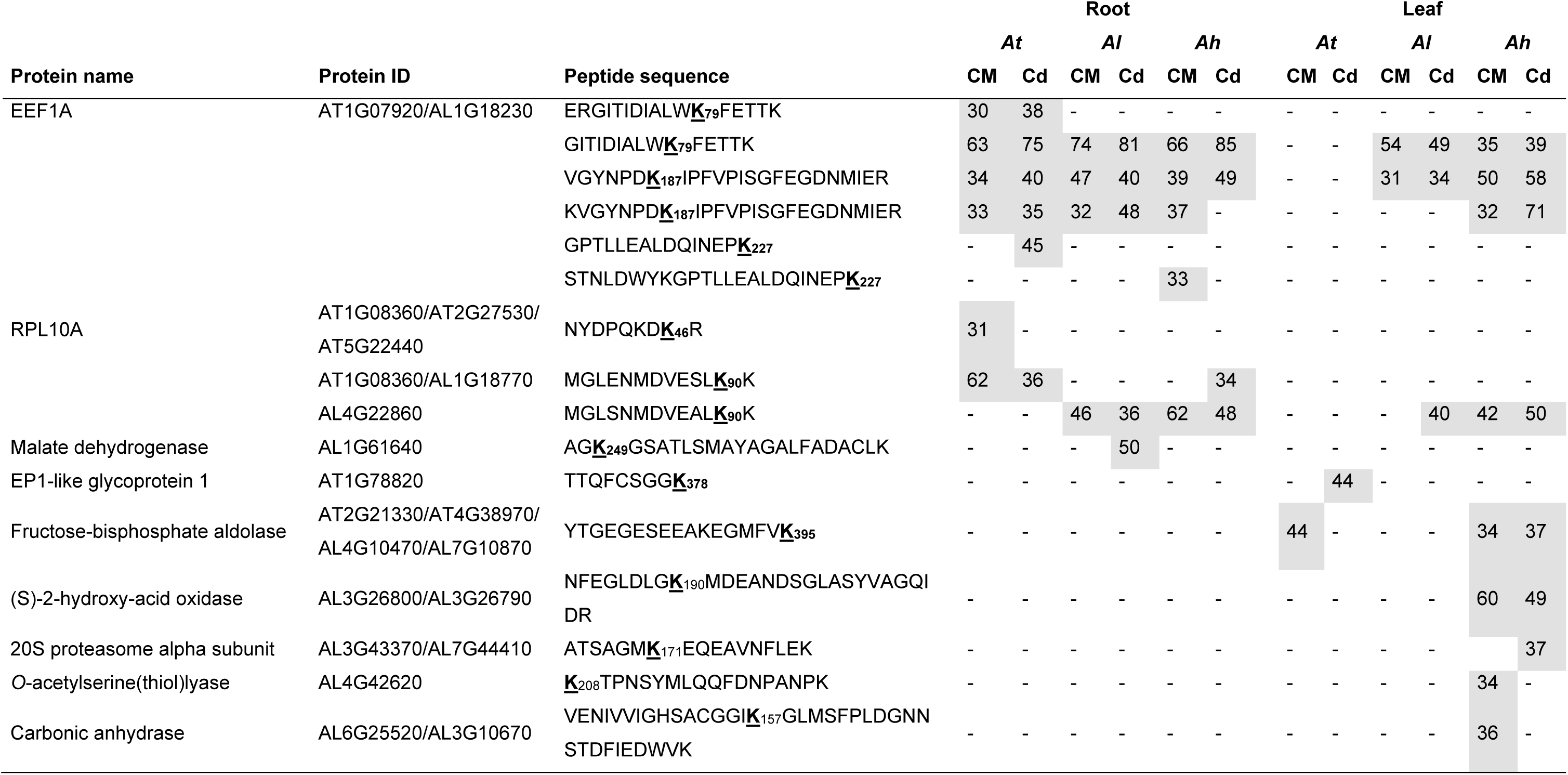
Lys-trimethylated proteins identified by MS/MS in root and leaf samples from Arabidopsis plants challenged with Cd. Soluble proteins were extracted from root and leaf tissues from Arabidopsis plants grown in control medium (CM) or challenged with 5 µM Cd for 9 days. Following SDS-PAGE, protein bands in the range 25-30 kDa were excised from the gel, digested with trypsin and analyzed by MS/MS using a Q Exactive Plus Orbitrap mass spectrometer. MS/MS data were searched for peptides bearing Lys trimethylated peptides as detailed in the Methods section. Sixteen Lys trimethylated peptides belonging to nine proteins have been identified with high confidence. Peptides detected in at least one of the 12 samples with Mascot scores ≥30 are shown in grey boxes. A dash indicates that the peptide was not detected in the corresponding sample (or with a Mascot score <30). *At, A. thaliana; Al, A. lyrata; Ah, A. halleri* (AU population). A comprehensive description of peptide properties and representative MS/MS spectra are available in Table S2 and Figure S3, respectively.

In root samples, where two protein bands at 26-28 kDa were strongly labeled with the trimethyl-Lys antibodies in all species and conditions (Figure 1c), we identified Lys-trimethylated peptides belonging to the Eukaryotic Elongation factor 1A (EEF1A), the ribosomal protein L10 (RPL10A), and a malate dehydrogenase. For malate dehydrogenase, the peptide bearing the previously unknown trimethylated Lys249 was detected only in the extract from *A. lyrata* plants treated with Cd (Table 1). For EEF1A, we identified three Lys trimethylation sites, two of them (Lys79 and Lys187) were detected in the three Arabidopsis species and were already known in several plant species (Lopez-Valenzuela *et al*., 2003; Ndamukong *et al*., 2011; Alban *et al*., 2014) while the third one (Lys227) was only detected in *A. thaliana* and was previously unknown. For RPL10A, two Lys trimethylation sites were identified, the first (Lys90) was formerly identified in *A. thaliana* (Carroll *et al*., 2008) while the second (Lys46) was not known. The identification of known methylation sites in EEF1A and RPL10A validated the overall pipeline for methylpeptide search and the use of the *A. lyrata* genome for MS/MS spectra assignation in both *A. lyrata* and *A. halleri*.

In leaf samples, where the immunodetection of the doublet of Lys-methylated proteins is species- and condition-dependent (Figure 1a), we identified trimethylated Lys residues in EEF1A, RPL10A, and six additional proteins (Table 1). Methylation of chloroplastic fructose 1,6-bisphosphate aldolases at a specific Lys residue (Lys395) was reported earlier (Mininno *et al*., 2012; Alban *et al*., 2014; Ma *et al*., 2016), while the other proteins were not previously known to be methylated.

We compared western blot and MS/MS analyses to try to assign the major trimethylated proteins at 26-28 kD. The detection pattern of peptides from RPL10A bearing a trimethylated Lys90 in root and leaf samples (Table 1) matched exactly the signals obtained with the antibodies against trimethyl-Lys (Figure 1a). For EEF1A, the overlap between methylpeptides and immunoblotting signals was also important. However, EEF1A is a very abundant cytosolic protein of about 50 kDa (Figure S4), suggesting that its identification in bands of 25-30 kDa was due to the high sensitivity of MS/MS detection and presumably protein smearing. Although the approach we used did not provide quantitative information about the identified methylpeptides, these results suggest that RPL10A (25 kDa) could contribute to one of the two intense signals observed by western blot. Despite the use of a high sensitive mass spectrometer and a robust identification pipeline, we have reached the limits of our approach and were not able to identify the second Lys-trimethylated protein, presumably abundant and interesting regarding Cd stress. Because of this technical bottleneck to investigate further the role of Lys-methylated proteins during metal stress, we moved on to the analysis of KMTs, the main drivers in the dynamics of protein Lys methylation.

### Expression of *A. thaliana* genes coding protein Lys methyltransferases in response to Cd

In *A. thaliana*, 48 genes coding KMTs from the SET domain-containing group (SDG) have been identified (Serre *et al*., 2018). Only two KMTs belonging to the seven-beta-strand (SBS) superfamily have been yet characterized in plants, namely the cytosolic enzyme CaMKMT that methylates calmodulin (CaM) (Banerjee *et al*., 2013) and the PrmA methyltransferase that modifies ribosomal protein L11 in plastids and mitochondria (Mazzoleni *et al*., 2015). Using BLAST searches, we identified 11 genes from *A. thaliana* that are orthologous to bacterial, yeast and human KMTs with a SBS structural fold (Figure S5). Thus, as a whole, the set of genes coding putative KMTs in *A. thaliana* comprised 59 members, with 48 *SDG* genes and 11 *SBS* genes.

In order to determine whether Cd could regulate the expression of *KMT* genes in *A. thaliana*, we analyzed transcriptomic datasets from published works (Herbette *et al*., 2006; Weber *et al*., 2006; Li *et al*., 2010; Jobe *et al*., 2012; Khare *et al*., 2016; Fischer *et al*., 2017). These datasets correspond to different conditions of stress with variations in Cd concentration (1 to 200 µM), treatment duration (2 hours to 7 days), growth medium (agar plates or hydroponics), and stage of development (7-day-old seedlings to 5-week-old mature plants). The coverage of *KMT* genes was important in each of the microarray experiment (51 to 59 genes identified out of 59). We found that the expression of some *KMT* genes was regulated by Cd (Table S3). Most of the differentially expressed genes (DEGs) were found in an experiment with drastic conditions of stress (200 µM Cd for 6 hours in hydroponics) (Li *et al*., 2010). In these conditions, Cd triggered the up-regulation of 12 genes and the down-regulation of three genes in roots (Table S3). Among these genes only *SBS7* was differentially regulated at a lower Cd concentration. Also, the expression of *SDG29* was upregulated following a short-term exposure to Cd. Together, these data indicate that the expression of a limited number of *KMT* genes is influenced by Cd in *A. thaliana*.

### Expression of genes coding protein Lys methyltransferases in *A. halleri* populations with different properties of Cd accumulation

To analyze whether Cd could modify the expression of genes coding KMTs in the Cd-tolerant species *A. halleri* we first used the comparative transcriptomic analysis from Weber *et al*. (2006). In this study, in which *A. halleri* plants from the population Langelsheim (Germany) were challenged with 25 or 125 µM Cd in hydroponic conditions for a short period (2 hours), none of the *KMT* genes was differentially expressed. Then, the expression of *KMT* genes was analyzed in the I16 and PL22 populations challenged with Cd. After four weeks of acclimatization in hydroponic growth medium, plants were treated with 5 µM CdSO_4_ for 10 days and transcriptomic analysis was performed in root and shoot samples using RNA sequencing (Corso *et al*., 2018). Genes coding KMTs were retrieved from the RNAseq data and their expression was analyzed. A principal component analysis (PCA) showed that the factor having the strongest impact on the expression profiles of *KMT* genes is the genetic unit (PL22 vs I16), accounting for 52 and 65% of the variance in roots and shoots, respectively (Figure 2a). The effect of the treatment (Cd vs control medium) was less important, accounting for 34 and 19% of the variance in roots and shoots, respectively (Figure 2a). DEGs were then identified in two pairwise comparisons to estimate the effect of the genetic unit and the treatment (p<0.05, threshold set at 1.4-fold change) (Corso *et al*., 2018). In agreement with the PCA, the PL22/I16 comparison identified 16 DEGs in roots and 10 DEGs in shoots (Figure 2b), whereas the Cd/control comparison yielded only four DEGs in PL22 and none in I16 (Figure 2c). In PL22, three genes were induced by Cd in roots (*SBS2* and *SBS9*) or in shoots (*SDG52*), and the *SBS5* gene was down-regulated by Cd in roots. The four genes regulated by Cd are predicted to code for KMTs modifying non-histone substrates (Serre *et al*., 2018), suggesting that these methylation events could be related to the tolerance and/or accumulation properties of the metallicolous PL22 population.

**Figure 2:**
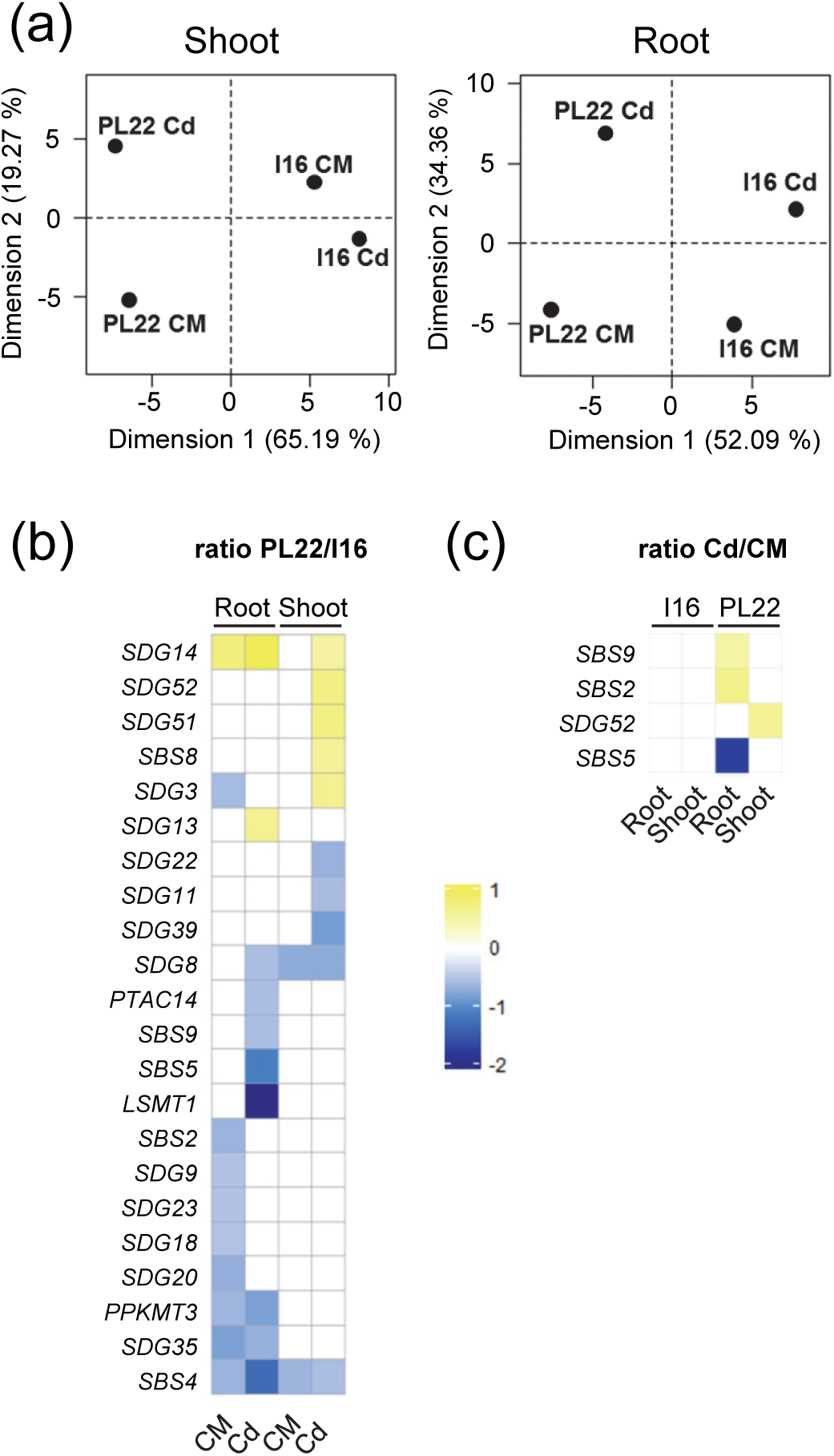
Expression of genes coding KMTs in the PL22 and I16 metallicolous populations of *A. halleri* challenged with Cd. RNAseq analysis was performed using root and shoot samples from plants exposed to 5 µM CdSO_4_ for 10 days (Corso *et al*., 2018). (a) - Principal component analysis of *KMT* genes expression in roots and shoots. (b) - Differentially expressed *KMT* genes according to the genetic unit (PL22 vs I16). The ratio between the steady-state expression level of each *KMT* gene in PL22 over I16 was calculated in all conditions (root ± Cd, shoot ± Cd) and DEGs were selected using a log2 fold-change ≥0.5 or ≤-0.5. (c) - Differentially expressed *KMT* genes according to the Cd treatment (Cd vs control). The ratio between the steady-state expression level of each *KMT* gene in Cd-treated over untreated plants was calculated in all conditions (root and shoot from I16 and PL22) and DEGs were selected using a log2 fold-change ≥0.5 or ≤-0.5. CM, control medium; Cd, medium containing 5 µM CdSO_4_.

### Identification of protein Lys methyltransferase mutants from *A. thaliana* with altered tolerance to Cd

We used a screening procedure with knock-out mutants to determine whether some *KMT* genes could play a role in the response of *A. thaliana* to Cd. We included only genes coding for KMTs modifying, or predicted to modify, non-histone substrates (Serre *et al*., 2018). Our selection comprised all genes (11) coding SBS enzymes and 15 genes coding SDG enzymes from classes VI and VII. SDG enzymes from classes I to V are known to methylate histones and some of them also accept non-histone substrates (Serre *et al*., 2018). Genes coding these enzymes were not included in our analysis since mutations in KMTs acting on histones, or on histones plus non-histone substrates, can lead to pleiotropic effects (e.g. Ndamukong *et al*., 2011), thus complicating the interpretation of the screening results. We obtained homozygous T-DNA insertion lines disrupting 23 of the selected genes (Table S1). Three genes could not be retained for the screening, of which *PAP7* for which the mutation is lethal in photoautotrophic conditions (Grübler *et al*., 2017).

We analyzed mutant seedlings for root growth inhibition by Cd, which is a simple and efficient method to assess tolerance to a toxic element (Remy & Duque, 2016). The procedure was set up using Col-0 seedlings and the *cad2.1* null mutant that is hypersensitive to Cd (Howden *et al*., 1995). In brief, 4-day-old seedlings were transferred to MS/2 medium supplemented or not with 20 µM CdSO_4_ and grown vertically for another 10 days in photoautotrophic conditions (no source of reduced carbon added to the medium) (Figure 3). Root length at day 8 and root elongation rate between days 3 and 8 were used as primary criteria to assess tolerance of the mutant lines to Cd (Figure 3). To address line-dependent differences in root growth that could interfere with the interpretation of the screening we also calculated the tolerance index (TI) for the two primary parameters, which corresponds to the ratio between the values in Cd-containing over control medium (Metwally *et al*., 2005) (Figure 3). The concentration of Cd in the medium (20 µM) was selected to produce a significant root growth inhibition (TI about 0.5) and to allow the identification of insertion lines that are either more tolerant or more sensitive to Cd than Col-0 in our experimental conditions.

**Figure 3:**
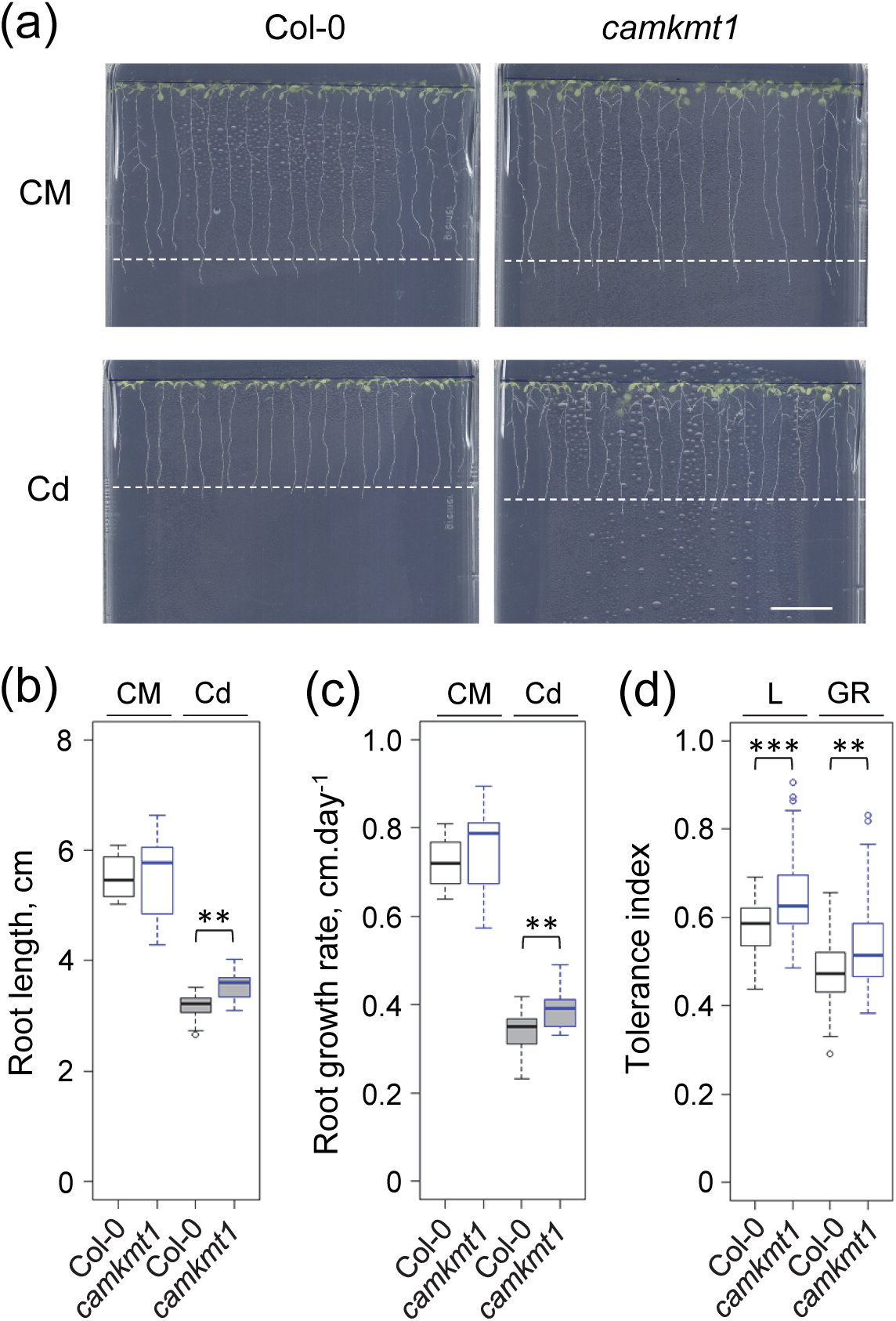
Root growth inhibition assays designed to analyze the tolerance to Cd of *KMT* mutants from *A. thaliana*. Results obtained for the *camkmt1* mutant are shown. Four-day-old seedlings (20 per genotype and condition) were transferred to square Petri dishes containing MS/2 medium (CM) or MS/2 with 20 µM CdSO_4_ (Cd) and grown in a vertical orientation. (a) Pictures were taken after eight days of treatment. Dotted lines show mean root length. Scale bar = 2 cm. (b) Effect of Cd on root length. Measurements have been done at day 8. (c) Effect of Cd on root growth rate. Measurements have been done between days 3 and 8. (d) Tolerance indices for Cd. TIs (ratio Cd/CM) have been calculated for root lengths (L) and root growth rates (GR). Data distribution is displayed in Tukey’s boxplots with the median as the solid line inside the box, the first and third quartiles as the bottom and top lines of the box, and whiskers with maximum 1.5 interquartile range of the lower and upper quartile, respectively. Outliers are plotted as individual dots. Each distribution represents n=20 seedlings (b, c) and n=100 calculations of TI (d). Statistical significance determined using a non-parametric Dunnett’s test is shown, with p <0.01 (**), and p <0.001 (***).

The results of the screening procedure have been summarized in a heat-map displayed in Figure 4. Mutants were clustered in three mains categories. First, the calculated TIs for 14 insertion lines were comparable with the Col-0 ecotype. Second, five mutants (*sdg51*, *sdg52*, *camkmt1*, *sbs7*, and *sbs9*) displayed a higher tolerance to Cd than the wild-type. Third, four mutants (*sdg50*, *sbs2*, *sbs6*, and *sbs8*) were found more sensitive to Cd than the wild-type. Together, the screening procedure allowed for the identification of nine out of 23 insertion mutants with a tolerance to Cd that is significantly different from that of the wild-type ecotype, suggesting that protein Lys methylation is part of the responses used by *A. thaliana* to cope with Cd stress.

**Figure 4:**
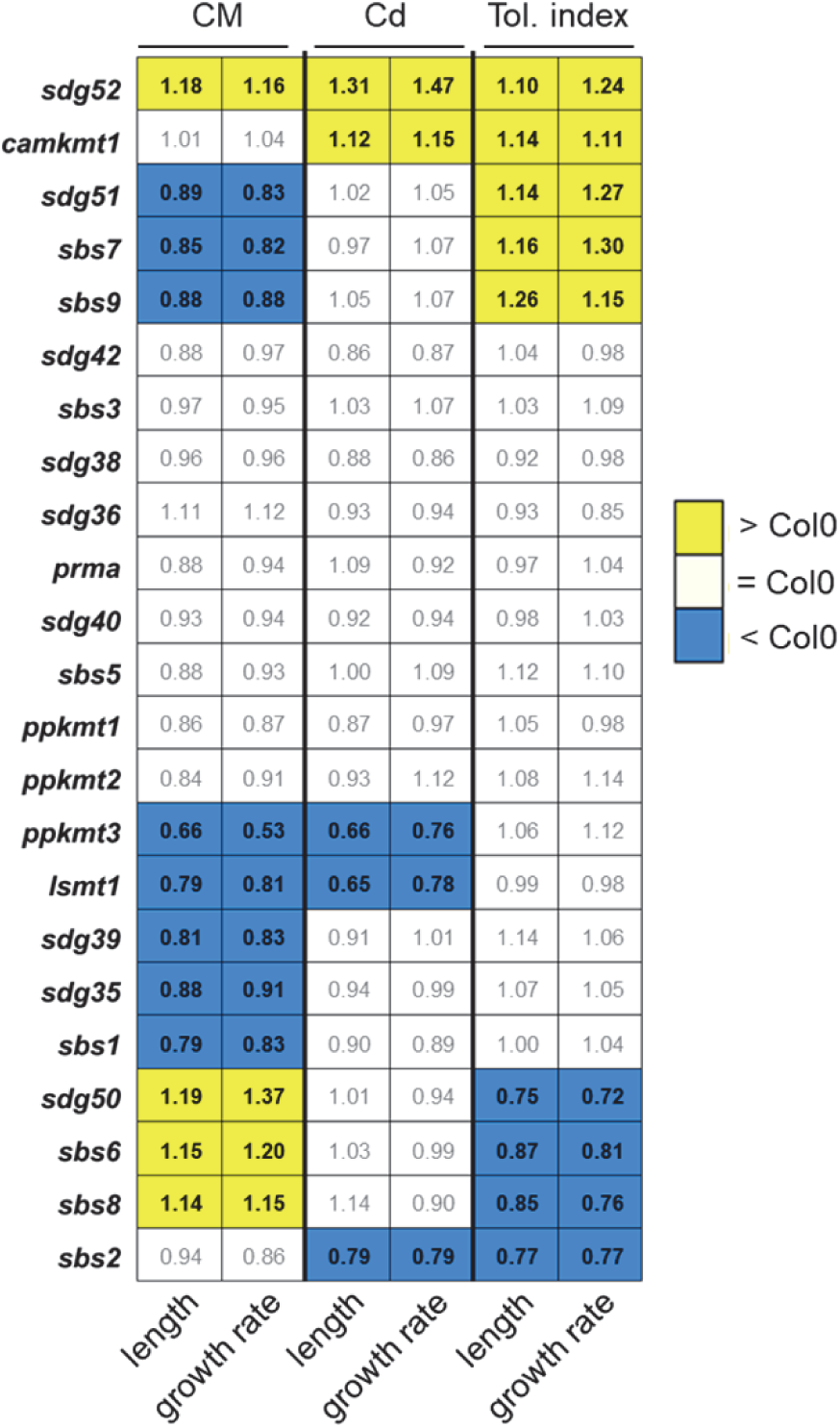
Heatmap summarizing the genetic screening of *KMT* mutants from *A. thaliana* for their tolerance to Cd. Each line identifies a *KMT* insertion line, each column defines the primary parameters of the screening procedure (root length at day 8 and root growth rate from day 3 to 8 in control and Cd-containing medium) and the calculated tolerance indices (ratio Cd/CM). For each parameters, statistical analysis indicated whether a *KMT* mutant was similar (white box), lower (blue box) or higher (yellow box) than the wild-type ecotype Col-0. Values indicate the ratio between the mutant and Col-0.

### Characterization of a Cd-tolerant mutant deficient in calmodulin Lys methyltransferase

Two mutants identified in the screening were selected for further investigations. The first insertion line, *camkmt1*, was found more tolerant to Cd than the wild-type (Figure 4) and is inactivated in the *CAMKMT* gene coding the CaM Lys methyltransferase (Banerjee *et al*., 2013). A previous analysis of the *camkmt1* null-mutant showed that disruption of the *CAMKMT* gene abolished CaM methylation at Lys315 and revealed a link between the methylation status of CaM and seedling tolerance to salt, heat, and cold stress (Banerjee *et al*., 2013).

The tolerance to Cd of the *camkmt1* knock-out line was verified using root growth assays and seedling biomass measurements using variable concentrations of the toxic metal (from 5 to 20 µM). For root elongation inhibition, the improved tolerance of *camkmt1* was significant only at the highest Cd concentration (Figure 5a). For seedling growth inhibition, the inhibitory effect of Cd on biomass was significantly less important for *camkmt1* than for the wild-type at 10 and 20 µM Cd (Figure 5b). CaMKMT is involved in the methylation of the major calcium (Ca) sensor CaM (Banerjee *et al*., 2013) and Ca is known to alleviate Cd toxicity (Suzuki, 2005; Baliardini *et al*., 2015). Consequently, the tolerance of *camkmt1* was analyzed using a fixed concentration of Cd (20 µM) and fluctuating concentrations of Ca (0.5, 1, and 1.5 mM). Changes in Ca availability did not modify the growth of seedlings in the absence of Cd (Figure 6). The inhibition of root elongation and seedling biomass by Cd was inversely correlated to Ca concentration in the medium. Also, the *camkmt1* line was found significantly more tolerant to Cd than the wild-type at each Ca concentration tested (Figure 6). Together, these data validated our screening approach and confirmed the identification of a Cd-tolerant *A. thaliana* mutant affected in the methylation of CaM.

**Figure 5:**
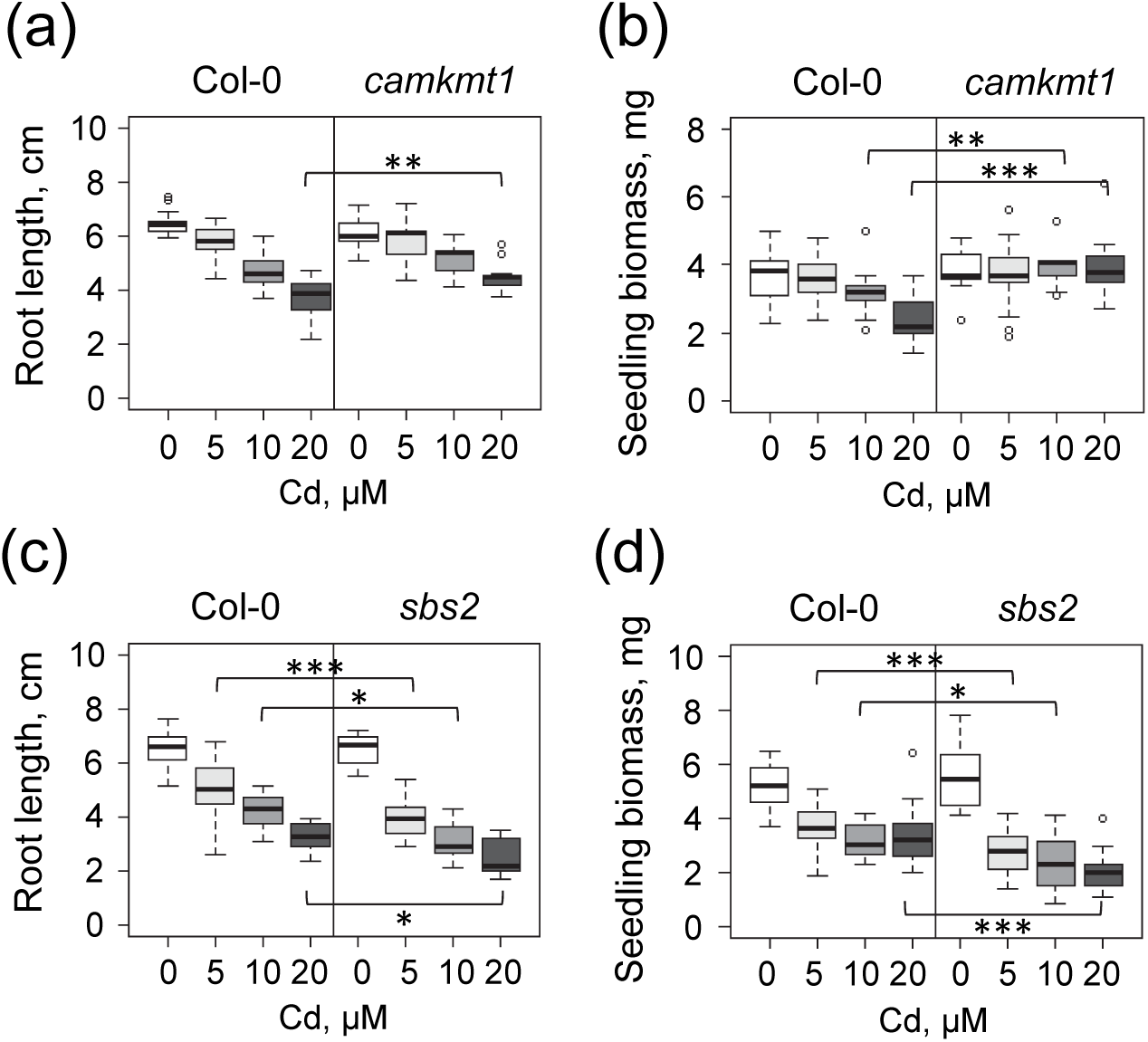
Tolerance to Cd of the protein Lys methyltransferases *camkmt1* and *sbs2* mutants. Four-day-old seedlings were transferred onto MS/2 medium containing various amount of CdSO_4_ and grown vertically for 10 days in photoautotrophic conditions. (a,c) – Dose-dependent inhibition of root growth by Cd. Root length was measured at day 8. (b,d) – Dose-dependent inhibition of seedling biomass by Cd. Seedling fresh weight was measured at day 10. Each distribution represents n=20 seedlings. Statistical significance determined using a non-parametric Dunnett’s test is shown, with p <0.05 (*), p <0.01 (**), and p <0.001 (***).

**Figure 6:**
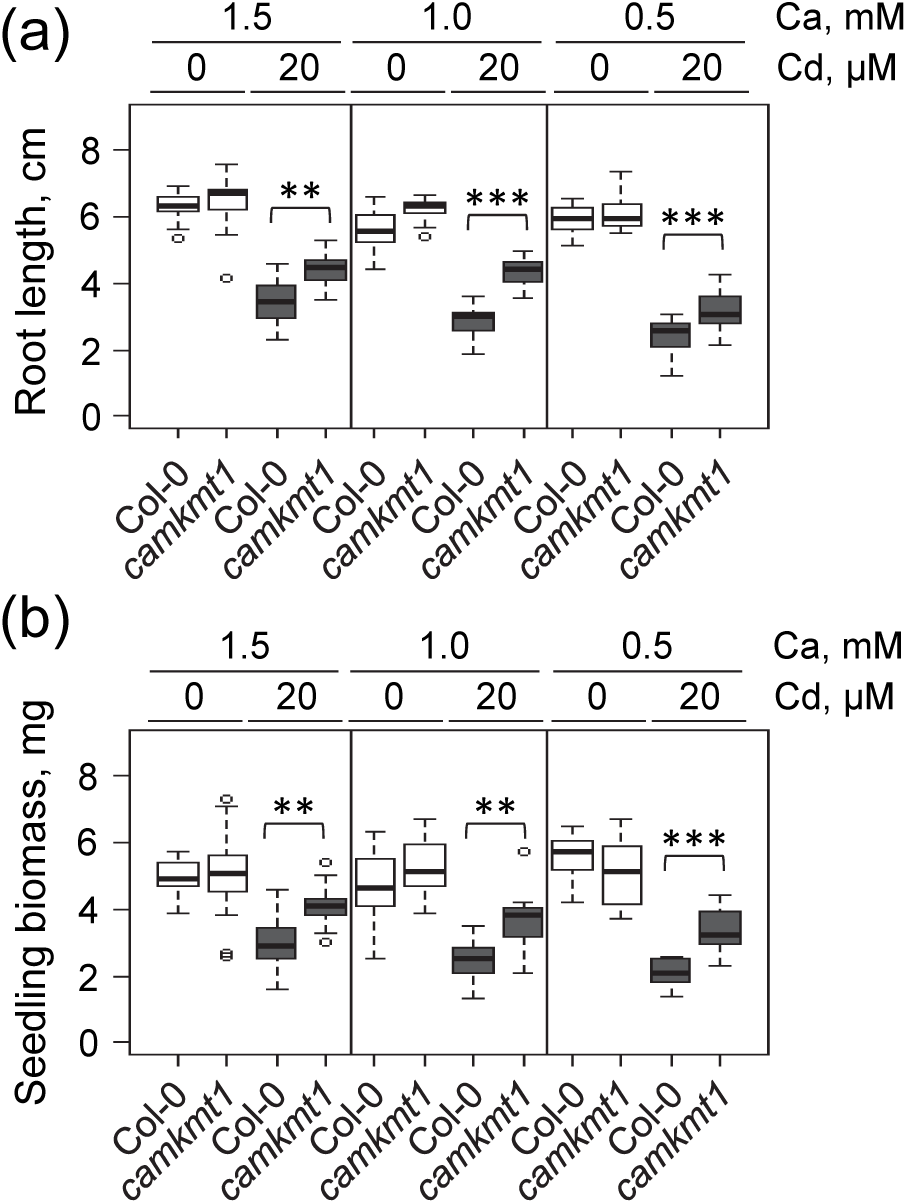
Tolerance to Cd of the calmodulin Lys methyltransferase *camkmt1* mutant. Four-day-old seedlings were transferred onto MS/2 medium containing various amount of CdSO_4_ and CaCl_2_ and grown vertically for 10 days in photoautotrophic conditions. (a) – Calcium-dependent inhibition of root growth by Cd. Root length was measured at day 8. (b) – Calcium-dependent inhibition of seedling biomass by Cd. Seedling fresh weight was measured at day 10. Each distribution represents n=20 seedlings. Statistical significance determined using a non-parametric Dunnett’s test is shown, with p-value <0.01 (**), and p-value <0.001 (***).

Then, we used ICP-MS to determine whether the difference in Cd-tolerance of *camkmt1* could be due to changes in its capacity to take up the element from the environment. Cadmium was measured in roots and shoots of plants grown in hydroponics and challenged with various Cd concentrations (0.2, 1 and 5 µM) for 7 days. There was no significant difference in the absorption and translocation of Cd in *camkmt1* as compared to Col-0 (Figure S6). Thus, the tolerance to Cd of *camkmt1* was not due to changes in Cd accumulation but rather to an improved capacity to cope with the toxic element.

### Characterization of a Cd-sensitive mutant affected in the protein Lys methyltransferase SBS2

The *sbs2* line was selected for further investigations because it is more sensitive to Cd (Figure 4) and the *SBS2* gene is upregulated in the roots of the *A. halleri* PL22 population challenged with Cd (Figure 2c). Yet, the function of the *SBS2* gene is unknown.

Similar to *camkmt1*, we first confirmed the phenotype of *sbs2* by measuring the inhibition of root elongation and seedling growth with different concentrations of Cd. Root growth of *sbs2* was significantly more inhibited by Cd than the wild-type at all concentrations tested (5 to 20 µM; Figure 5c). Also, the biomass of *sbs2* seedlings was lower than Col-0 seedlings for the three concentrations tested (Figure 5d), confirming the Cd-sensitive phenotype of *sbs2*.

To gain insight into the role of the *SBS2* gene in the response to Cd we selected a second independent insertion line, referred to as *sbs2b*. The T-DNA insertions were located in the fourth exon of *SBS2* for *sbs2b* and downstream the fourth exon for *sbs2*, in a region that is either an intron or the 3’ untranslated region of *SBS2* transcript variants (Figure 7). Reverse transcription-PCR analysis indicated that the two lines are loss-of-function alleles with no detectable *SBS2* transcripts. Also, root growth assays showed that *sbs2b* behaved as *sbs2* and was less tolerant to Cd than wild-type seedlings (Figure 7). Together, these data indicated that the invalidation of the *SBS2* gene is responsible for an increased sensitivity to Cd.

**Figure 7:**
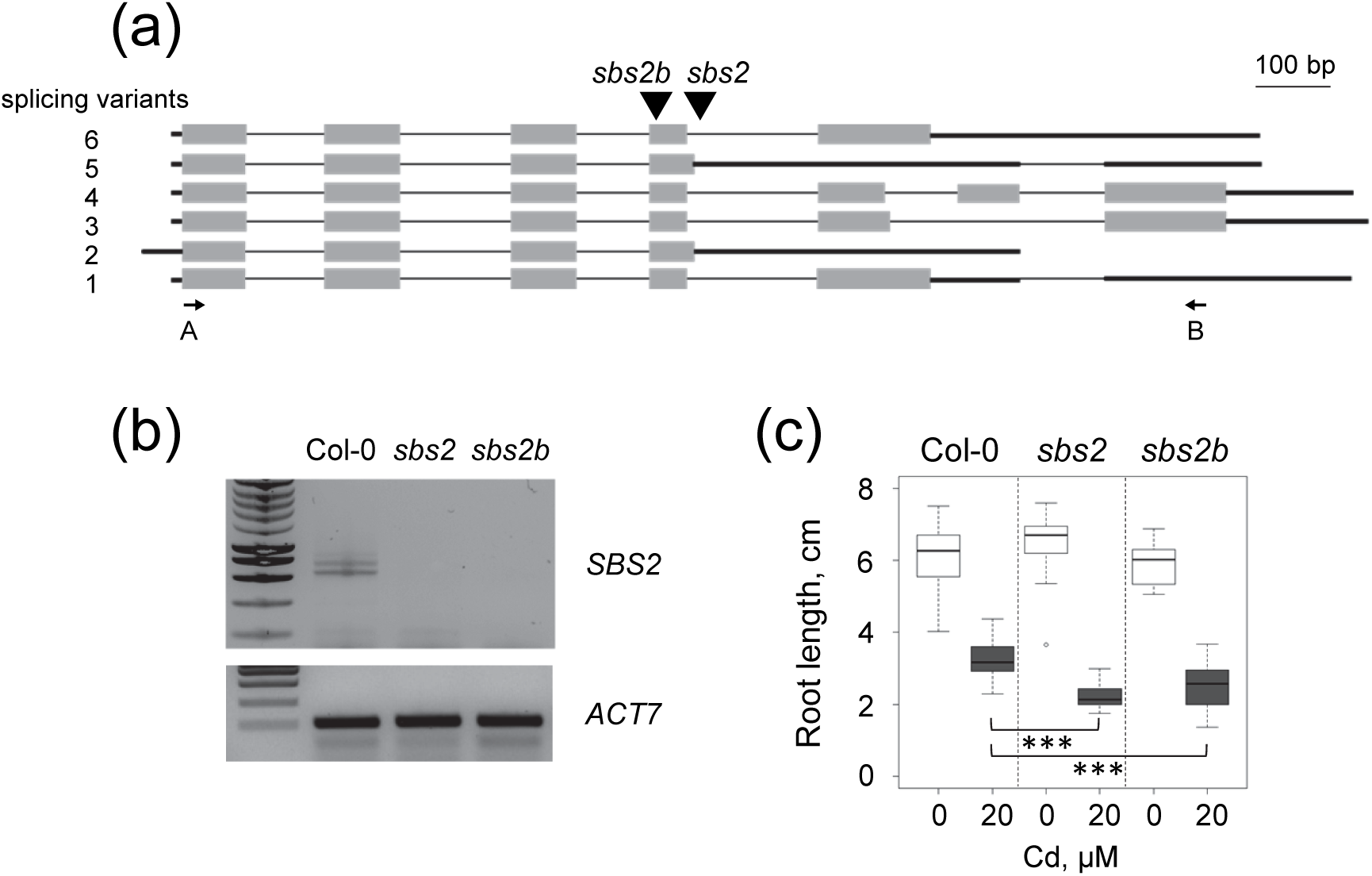
Molecular characterization of the *A. thaliana sbs2* and *sbs2b* mutants. (a) - Structure of the *SBS2* locus. Splicing variants predicted in the Araport database are shown (https://apps.araport.org/thalemine/portal.do?externalids=AT1G63855). Thin lines represent introns, grey boxes represent exons, thick dark lines represent untranslated regions, and triangles indicate T-DNA insertion sites in the *sbs2* (GK-911F08) and *sbs2b* (SALK_037552) mutants. Primers A and B are indicated by arrows. (b) - Amplification of *SBS2* transcripts by PCR. PCR was performed using reverse transcribed total RNA from 10-day-old seedlings with primers A (ATGATGACTACTACGACGACGAC) and B (CTCAATACGATCTCAACCAACTGA) for *SBS2*, and ACT7-F (ACATCGTTCTCAGTGGTGGTCC) and ACT7-R (ACCTGACTCATC-GTACTCACTC) for *ACTIN7*. PCR products were resolved by agarose gel electrophoresis. Two major amplicons of 700 to 800 bp were amplified in Col-0, cloned and sequenced. They correspond to splicing variants 1 and 4. (c) - Tolerance to Cd of the *sbs2* and *sbs2b* mutants. Four-day-old seedlings were transferred onto MS/2 medium supplemented or not with 20 µM CdSO_4_ and grown vertically for 10 days in photoautotrophic conditions. Root length was measured at day 8. Each distribution represents n=20 seedlings. Statistical significance determined using a non-parametric Dunnett’s test is shown, with p-value <0.001 (***).

We analyzed whether the uptake and distribution of Cd was affected in *sbs2*. The Cd content in roots and shoots of *sbs2* was similar to that of Col-0 at any Cd concentration tested (Figure S6). Thus, the increased sensitivity to Cd of *sbs2* was not associated with an increased absorption of the toxic element from the medium but rather to a reduced capacity to deal with its deleterious effects.

Last, as a preliminary approach to identify the substrate of the SBS2 methyltransferase, we used western blot analysis to compare the patterns of Lys-trimethylated proteins in *sbs2* and Col-0. We could not observe any significant decrease in band intensity (hypomethylation) in *sbs2* relative to Col-0 in soluble or membrane proteins from roots and shoots of seedlings grown in standard conditions (Figure S7). Thus, the substrate of SBS2 is probably a low abundant protein that was not detectable by the current immunolabeling approach.

## DISCUSSION

The methylation status of Lys residues in proteins is controlled by KMTs and contributes to the regulation of protein properties in diverse biological processes. To address whether Lys methylation of non-histone proteins is important for metal tolerance in Arabidopsis species, we analyzed the effect of Cd on the two partners participating in this PTM, i.e. methylated proteins on the one hand and KMTs on the other hand. Using an immunoblotting approach, we showed that the Lys-methylation status of some proteins is influenced by a Cd stress in the roots and leaves of Arabidopsis (Figure 1). Changes in methylation patterns were observed between Cd-tolerant and Cd-sensitive species and between treated and untreated plants. This analysis provided the first evidence that the steady-state level of some methylproteins, or the stoichiometry of Lys methylation of these proteins, could be linked with metal stress and with the genetic diversity of the Arabidopsis species. Then, we used MS/MS to identify Lys-trimethylated proteins of low molecular weight (25-30 kDa) that displayed different expression profiles in the leaves of *A. thaliana*, *A. lyrata* or *A. halleri* in response to Cd. Using a specific pipeline for the identification of Lys trimethylation events we identified 12 methylsites in nine proteins (Table 1). Six of these proteins and eight of the Lys-methylated sites were not previously known, illustrating the depth of the analysis. In addition, by using genomic resources of *A. lyrata* for the assignment of MS/MS spectra from *A. lyrata* and *A. halleri* samples, we were able to identify, for the first time, post-translationally modified proteins in these model species.

The methylation status of only one of the identified methylproteins, RPL10A, was correlated with the different responses of Arabidopsis species upon Cd stress in leaves. RPL10A is involved in translation as a subunit of the 60S large ribosomal subunit and has non-canonical functions linked with its translocation to the nucleus. RPL10A is an essential protein in plants since knockout mutants are lethal and *rpl10a/RPL10A* heterozygous plants are deficient in translation under UV-B stress conditions (Falcone Ferreyra *et al*., 2010). Also, RPL10A is a substrate of the receptor-like kinase NIK1 and its phosphorylation redirects the protein from the cytosol to the nucleus where it may act to modulate viral infection (Carvalho *et al*., 2008). We identified two Lys trimethylation sites in Arabidopsis RPL10A proteins. The first one (Lys46) has been previously identified as monomethylated by the RKM5 methyltransferase in the homolog of RPL10A from yeast (Webb *et al*., 2011). Trimethylation of Lys46 was detected only in the roots of *A. thaliana* grown in control conditions and, so, has probably no link with the response to metal stress. This assumption is supported by the observation that a mutation in the *SBS1* gene, the ortholog of RKM5 (Figure S5), did not change the tolerance to Cd of *A. thaliana* seedlings (Figure 4). The pattern of trimethylation of the second residue (Lys90) in RPL10A in leaves was influenced by Cd stress in a species-depend manner (Table 1). The functional outcome of Lys90 trimethylation in RPL10A is not known; the modification may contribute to the optimization of ribosomal function or may affect its non-canonical functions, as previously showed for RPL10A phosphorylation (Carvalho *et al*., 2008).

We also analyzed the expression of genes coding protein Lys methyltransferases in response to Cd in wild-type *A. thaliana* and in populations of *A. halleri* with different capacities to tolerate and accumulate the toxic metal. In *A. thaliana,* the steady-state level of only two *KMT* genes is regulated by moderate concentrations of Cd (Table S3). In *A. halleri*, we showed that Cd induces a significant change in the expression of four *KMT* genes in the PL22 population, but none in the I16 population (Figure 2c). The transcriptomic, ionomic and metabolomic analysis of these two metallicolous populations from different European genetic units indicated that distinct strategies driven by different sets of genes have evolved for the adaptation to high Cd (Corso *et al*., 2018) or high zinc in soils (Schvartzman *et al*., 2018). Since PL22 accumulates Cd in roots and shoots whereas I16 behaves as a Cd excluder, both *in situ* and in hydroponic conditions, these results suggest that the regulation of *KMT* genes expression in PL22 could be correlated with the level of Cd that is taken up from the environment and translocated to shoots. The substrates of the KMTs encoded by these four genes (*SDG52*, *SBS2*, *SBS5*, and *SBS9*) are likely not histones (Figure S5), suggesting that Lys methylation of non-histone proteins could contribute to the regulation of cellular mechanisms involved in Cd accumulation or detoxification in the PL22 population. The analysis of DEGs between I16 and PL22, regardless of the presence of Cd in the culture medium, identified 22 *KMT* genes (Figure 2b). This suggests that Lys methylation of histones and non-histone substrates could be part of the diverging adaptation strategies of metallicolous populations. The expression of *KMT* genes coding enzymes of the SDG family has been previously analyzed in cotton plants stressed with high temperature (Huang *et al*., 2016) and in foxtail millet under different abiotic stresses (Yadav *et al*., 2016). In these studies, the expression pattern of some *KMT* genes was significantly changed in stress conditions. These data, together with our results, suggest that protein Lys methylation could play a role in the responses of plants to a variety of abiotic stresses.

Last, we used a screening procedure based on root growth inhibition assays to determine whether some *KMT* genes could be important for *A. thaliana* to cope with Cd. We showed that nine out of 23 insertion mutants displayed a tolerance to Cd that was significantly different from that of wild-type seedlings (Figure 4). These KMTs belong to the SDG class VII (SDG50, SDG51, SDG52) and to the SBS family (SBS2, SBS6, SBS7, SBS8, SBS9, CaMKMT) and are known, or predicted, to modify non-histone targets (Serre *et al*., 2018), suggesting that Lys methylation of non-histone proteins is one of the regulatory mechanisms modulating the response of *A. thaliana* to Cd stress.

Two of the identified mutants were further investigated. The *camkmt1* line is unable to methylate CaM (Banerjee *et al*., 2013) and is more tolerant to Cd than the wild-type at each Ca concentration tested (Figure 5). Cadmium is known to interfere with Ca homeostasis and the Ca/CaM system has been hypothesized to participate in heavy metal signaling (Gallego *et al*., 2012; Baliardini *et al*., 2015). More generally, CaM has been implicated in the response and recovery to different stresses and CaM methylation has been proposed to play a regulatory role in these processes. Indeed, a *camkmt1* null mutant displayed increased tolerance to salt, heat and cold stress whereas lines overexpressing *CAMKMT* were hypersensitive to these stresses (Banerjee *et al*., 2013). Together, these data suggest that Lys methylation of CaM also plays a role in the signaling cascade triggered by Cd, probably at a level that is common between different abiotic stresses. The precise role of Lys methylation in the modulation of CaM activity is still unclear.

Our data also indicated that the invalidation of the *SBS2* gene in *A. thaliana* is associated with a decreased capacity to cope with Cd (Figure 7). Also, the expression of *SBS2* was increased in the roots of *A. halleri* PL22 plants challenged with Cd (Figure 2c), suggesting that the methylation reaction catalyzed by SBS2 is useful to limit the deleterious effects of Cd. The function of SBS2 is still not known in plants. Its ortholog in animal cells is METTL23 (Figure S5). METTL23 is located in the cytoplasm and the nucleus, interacts with a subunit of the GA-binding protein transcription factor, but its target(s) has not been yet identified (Bernkopf *et al*., 2014; Reiff *et al*., 2014). The profiling of methylproteins in *sbs2* and Col-0 by western blot was not sensitive enough to detect any change between the two lines (Figure S7), providing no clues to the nature of the substrate(s) of the SBS2 enzyme. The identification of the target(s) of SBS2 is the next step to gain insight into the role of this methylation event under favorable growth conditions and in the response to Cd stress.

Together, the data presented in this study provide the first evidence for a link between the methylation status of Lys in non-histone proteins and the response of plants to a stress induced by Cd. They pave the way for the identification of cellular mechanisms that are regulated by protein Lys methylation and are important for plants to cope with toxic elements. To reach this goal one has to identify the KMT/substrate relationships to be able to modulate the methylation status of protein targets *in vivo*. To summarize, this work suggests that the characterization of the KMT involved in the methylation of Lys90 in RPL10A and the identification of the substrates of the KMTs encoded by genes that are modulated by Cd and play a role in Cd tolerance will provide significant insights into the role of protein Lys methylation during metal stress.

## AUTHORS CONTRIBUTION

NBCS, VS, NV and SR conceived and designed the study; NBCS, MS, OG, SF, MC, JC, VR and SR performed the experiments; NBCS, MS, OG, MC, VR, CA, VS, JB, NV and SR analyzed the data; NBCS and SR wrote the paper, with the input from all co-authors.

## ACKNOWLEDGEMENTS

This work was supported by grants from the Région Auvergne Rhône-Alpes (PhD to NBCS), the Fondation de Coopération Scientifique Rovaltain (PlantStressMetalPTMs project), the Toxicology program of the Commissariat à l’Energie Atomique et aux Energies Alternatives, the Department of Plant Biology and Breeding of the Institut National de la Recherche Agronomique, the LabEx GRAL (ANR-10-LABX-49-01), and the Fonds de la Recherche Scientifique–FNRS (PDR T.0206.13 to NV). This paper is dedicated to the memory of Professor Roland Douce.

## SUPPORTING INFORMATION

**Table S1:** A. thaliana insertion lines used in this study.

**Table S2:** Properties of Lys-trimethylated peptides identified by MS/MS.

**Table S3:** Differentially expressed *KMT* genes in *A. thaliana* exposed to Cd.

**Figure S1:** Phenotype and photosynthesis of Arabidopsis plants challenged with Cd.

**Figure S2:** Immunodetection of Lys-trimethylated proteins in roots and leaves from I16 and PL22 *A. halleri* plants challenged with Cd.

**Figure S3:** LC-MS/MS fragmentation spectra of Lys-trimethylated peptides identified in *A. thaliana*, *A. lyrata*, and *A. halleri* protein samples.

**Figure S4:** Immunodetection of EEF1A in roots and leaves from Arabidopsis plants challenged with Cd.

**Figure S5:** Phylogenetic analysis of KMTs from the seven-beta strand (SBS) superfamily.

**Figure S6:** Absorption and translocation of Cd in the *camkmt1* and *sbs2* mutants.

**Figure S7:** Immunodetection of Lys-trimethylated proteins in roots and leaves from *sbs2* and Col-0 plants.

