## Supplemental data for "Protein lysine methylation is involved in modulating the response of sensitive and tolerant Arabidopsis species to cadmium stress"

**Table S1:** *A. thaliana* insertion lines used in this study.

**Table S2:** Properties of Lys-trimethylated peptides identified by MS/MS.

**Table S3:** Differentially expressed *KMT* genes in *A. thaliana* exposed to Cd.

**Figure S1:** Phenotype and photosynthesis of Arabidopsis plants challenged with Cd.

**Figure S2:** Immunodetection of Lys-trimethylated proteins in roots and leaves from I16 and PL22 *A. halleri* plants challenged with Cd.

**Figure S3:** LC-MS/MS fragmentation spectra of Lys-trimethylated peptides identified in *A. thaliana*, *A. lyrata*, and *A. halleri* protein samples.

**Figure S4:** Immunodetection of EEF1A in roots and leaves from Arabidopsis plants challenged with Cd.

**Figure S5:** Phylogenetic analysis of KMTs from the seven-beta strand (SBS) superfamily.

**Figure S6:** Absorption and translocation of Cd in the *camkmt1* and *sbs2* mutants.

**Figure S7:** Immunodetection of Lys-trimethylated proteins in roots and leaves from *sbs2* and Col-0 plants.

**Supplemental Table S1: *A. thaliana* insertion lines used in this study.**

| Gene ID | Common name | | Mutant line | Mutant name | Primers used for genotyping | | Screening for Cd tolerance |
|---|---|---|---|---|---|---|---|
| | | | | | Right primer | Left primer | |
| **Seven-beta-strand (SBS) putative KMTs** | | | | | | | |
| AT1G08125 | SBS1 | | SALK_124644C | *sbs1* | AAAGACGAAACCTCAAAAGGC | CTTCCTGCTATTTTTCCCCTG | Yes |
| AT1G63855 | SBS2 | | GK-911F08 | *sbs2* | TTTGGAGAAGAAATGAGACAGTCG | GTGATTCTTCGATACTCGAGGTTA | Yes |
| AT1G66680 | SBS3 | | GK-032H01 | *sbs3* | AGGTAACAGAAAGAAAGTGCGGTA | TGTTTCTGAAAACGATGTTACGAC | Yes |
| AT1G73320 | SBS4 | | | | | | No (mutant line not available) |
| AT1G79915 | SBS5 | | SALKseq_42690 | *sbs5* | CTGATTTCCATGGCATTGTG | TTCGCTATTATCCCCCAAAC | Yes |
| AT3G50850 | SBS6 | | GK-125E08 | *sbs6* | CCAAAGTATAAGGGTGTTCTGGAT | ATAACAACCCTCCACGTGTAACC | Yes |
| AT4G35987 | CaMKMT | | SALK_138607C | *camkmt1* | GCTCTGCTCTGTGTATTTGGC | TCAACTTTCAAGTCCACCCAC | Yes |
| AT5G27400 | SBS7 | | SALK_019618C | *sbs7* | AAATCCCGACCATACCAACTC | CTCCGTCATCACTCTCGCTAG | Yes |
| AT5G44170 | SBS8 | | SALK_128491C | *sbs8* | AAAACATCTCCGATTTGACCC | TCGCATAACTCCCAAAACAAC | Yes |
| AT5G49560 | SBS9 | | SALK_046849C | *sbs9* | CGGTAATGTACTGAAGCTCCG | GCTGACGTCACAAGGTCTCTC | Yes |
| AT5G53920 | PrmA | | SALK_070621 | *prma* | GAAGCTGCAAAAAGAGCATTG | GTTATGCTCAAAACTCAGCGC | Yes |
| **SET-domain group (SDG) KMTs** | | | | | | | |
| | | sub-class | | | | | |
| AT1G01920 | SDG42 | VII | SAIL_321_C10 | *sdg42* | TCCACATTCTTTCGTGTTTCC | TACTCGGAATTGCCTCAACAG | Yes |
| AT1G14030 | LSMT | VII | SAIL_1156_C01 | *lsmt1* | CCAGAGACAGTGACTGCTTCC | CACAAGCCGAAGGTACTGAAG | Yes |
| AT1G24610 | PPKMT2 | VII | GK-833G12 | *ppkmt2* | AAATTTCCAGTAATCGCATTGAAC | CCGTTTATGAATTTTGGTTCG | Yes |
| AT1G26760 | SDG35 (ATXR1) | VI | SALK_117606 | *sdg35* | ATCGTAGTACCGCTGCTGTTG | CCTAACGCAAGACGTCTTCAC | Yes |
| AT2G17900 | SDG37 (ASHR1) | VI | SALK_127952C | *sdg37* | AAGTTCCGCCTAACACTAGCC | TGCTTGCTTCTTAGGAGCAAG | No (insertion site not confirmed) |
| AT2G18850 | SDG50 | VII | SAIL_849_G09 | *sdg50* | GAACTTCCCAAAGATGATCCC | TGGTTTGAATTTCGAGTCCAG | Yes |
| AT2G19640 | SDG39 (ASHR2) | VI | SALK_024470C | *sdg39* | GGATAAATCCCATAAGCTCGC | TATGGACGGCTAAGTGTGGAG | Yes |
| AT3G07670 | PPKMT3 | VII | SALK_150387 (CA) | *ppkmt3* | AACCGAACCAAACCGTAAATC | ACTGCATCAATGATTTCCAGC | Yes |
| AT3G21820 | SDG36 (ATXR2) | VI | SALK_026154C | *sdg36* | TTGTGAAGCAGAAGTCTTCCTC | TCTCGATTGATCCAATGAACC | Yes |
| AT3G55080 | SDG51 | VII | SALK_057077C | *sdg51* | TTGAATTTTTGTTTTGGCCAC | TTGTCTTCATCCCTCAAAACG | Yes |
| AT3G56570 | SDG52 | VII | SALK_131900C | *sdg52* | CCATGGTAATCATCGATTTCG | AACCAAAATTCGGATTGAACAC | Yes |
| AT4G20130 | PAP7 (PTAC14) | VII | SAIL_566_F06 | *pap7* | TGCAGAGAATGATCAATCGTG | AGAAGGTCCAGATGGTTTTGG | No (lethal mutation in photoautotrophic conditions) |
| AT5G06620 | SDG38 (ATXR4) | VI | SAIL_1267_H02 | *sdg38* | TTGTCATGTATGAGATCAGCCC | TTAAGACATTGACGACGCCTC | Yes |
| AT5G14260 | PPKMT1 | VII | SALK_123180C | *ppkmt1* | TTTCCATACGAAATGCGTCTC | ATGGGTATTGCTGCAAAACAC | Yes |
| AT5G17240 | SDG40 | VII | SALK_097673C | *sdg40* | GATATCAGCGCAAAAGACAGC | CACTTTCTTCGACTCAGGTGC | Yes |

#### Supplemental Table S2: Properties of Lys-trimethylated peptides identified by MS/MS

##### *A.thaliana* / root samples

| | | | | | | | | | | CM | | | | | | Cd | | | | | |
|---|---|---|---|---|---|---|---|---|---|---|---|---|---|---|---|---|---|---|---|---|---|
| Sequence | # PSMs | # Proteins | # Protein Groups | Protein Group Accessions | Modifications | MH+ [Da] | q-Value | PEP | # Missed Cleavages | C2 | IonScore C2 | Exp Value C2 | Charge C2 | m/z [Da] C2 | RT C2 | F2 | IonScore F2 | Exp Value F2 | Charge F2 | m/z [Da] F2 | RT F2 |
| GITIDIALWKFETTK | 8 | 1 | 1 | AT1G07920.1 | K10(Trimethyl) | 1778.01091 | 0 | 1.122E-10 | 1 | High | **63** | 0.000167776 | 2 | 889.50818 | 118.13 | High | **75** | 1.16959E-05 | 2 | 889.50909 | 118.20 |
| KVGYNPDKIPFVPISGFEGDNMIER | 5 | 1 | 1 | AT1G07920.1 | K1(Trimethyl) or K8(Trimethyl) | 2864.46279 | 0 | 0.0002318 | 2 | High | **33** | 0.51608221 | 4 | 716.87384 | 112.49 | High | **35** | 0.274843058 | 4 | 716.87115 | 112.49 |
| ERGITIDIALWKFETTK | 4 | 1 | 1 | AT1G07920.1 | K12(Trimethyl) | 2063.15677 | 0 | 0.000002115 | 2 | High | **30** | 0.417713757 | 4 | 516.54468 | 111.46 | High | **38** | 0.065137207 | 3 | 688.39044 | 111.50 |
| GPTLLEALDQINEPK | 2 | 1 | 1 | AT1G07920.1 | K15(Trimethyl) | 1679.92380 | 0 | 0.0001456 | 0 | High | 27 | 0.685659191 | 3 | 560.64899 | 118.45 | High | **45** | 0.012110451 | 3 | 560.64612 | 118.47 |
| VGYNPDKIPFVPISGFEGDNMIER | 14 | 1 | 1 | AT1G07920.1 | K7(Trimethyl); M21(Oxidation) | 2752.37143 | 0 | 4.009E-08 | 1 | High | **34** | 0.357892601 | 3 | 918.12720 | 115.78 | High | **40** | 0.078825896 | 3 | 918.12665 | 115.80 |
| VGYNPDKIPFVPISGFEGDNMIER | 4 | 1 | 1 | AT1G07920.1 | K7(Trimethyl); N20(Deamidated); M21(Oxidation) | 2753.35819 | 0 | 1.015E-09 | 1 | High | 18 | 13.6220347 | 3 | 918.46259 | 116.46 | High | **49** | 0.009381666 | 3 | 918.45758 | 117.16 |
| VGYNPDKIPFVPISGFEGDNMIER | 1 | 1 | 1 | AT1G07920.1 | K7(Trimethyl) | 2736.37003 | 0 | 9.259E-12 | 1 | | | | | | | High | **59** | 0.00116692 | 3 | 912.79486 | 118.31 |
| MGLENMDVESLKK | 3 | 1 | 1 | AT1G08360.1 | M1(Oxidation); M6(Oxidation); K12(Trimethyl) | 1567.77019 | 0 | 0.0003545 | 1 | High | **62** | 0.000172064 | 2 | 784.38873 | 46.63 | High | 17 | 5.400472289 | 3 | 523.26239 | 46.65 |
| MGLENMDVESLKK | 11 | 1 | 1 | AT1G08360.1 | M1(Oxidation); K12(Trimethyl) | 1551.77165 | 0 | 0.0007926 | 1 | High | **51** | 0.002579271 | 2 | 776.38947 | 65.14 | High | **36** | 0.07650783 | 3 | 517.92993 | 65.25 |
| NYDPQKDKR | 1 | 3 | 3 | AT1G08360.1;AT2G27530.1;AT5G22440.1 | N1(Deamidated); K8(Trimethyl) | 1206.61100 | 0.006 | 0.08072 | 2 | High | **31** | 0.203673772 | 3 | 402.87518 | 18.37 | | | | | | |

##### *A.lyrata* / root samples

| | | | | | | | | | | CM | | | | | | Cd | | | | | |
|---|---|---|---|---|---|---|---|---|---|---|---|---|---|---|---|---|---|---|---|---|---|
| Sequence | # PSMs | # Proteins | # Protein Groups | Protein Group Accessions | Modifications | MH+ [Da] | q-Value | PEP | # Missed Cleavages | C2 | IonScore C2 | Exp Value C2 | Charge C2 | m/z [Da] C2 | RT C2 | F2 | IonScore F2 | Exp Value F2 | Charge F2 | m/z [Da] F2 | RT F2 |
| GITIDIALWKFETTK | 8 | 2 | 1 | AL1G18230.t1 | K10(Trimethyl) | 1778.00859 | 0 | 7.946E-09 | 1 | High | **74** | 1.17942E-05 | 2 | 889.50885 | 115.92 | High | **81** | 2.37502E-06 | 2 | 889.50885 | 115.84 |
| VGYNPDKIPFVPISGFEGDNMIER | 12 | 2 | 1 | AL1G18230.t1 | K7(Trimethyl); M21(Oxidation) | 2752.36740 | 0 | 2.048E-12 | 1 | High | **47** | 0.014710998 | 3 | 918.12732 | 113.35 | High | **40** | 0.071408189 | 3 | 918.12610 | 113.94 |
| KVGYNPDKIPFVPISGFEGDNMIER | 18 | 2 | 1 | AL1G18230.t1 | K1(Trimethyl) or K8(Trimethyl); M22(Oxidation) | 2880.46401 | 0 | 0.00002406 | 2 | High | **32** | 0.501013925 | 4 | 720.87097 | 98.90 | High | **48** | 0.011636072 | 4 | 720.87146 | 98.68 |
| AGKGSATLSMAYAGALFADACLK | 1 | 1 | 1 | AL1G61640.t1 | K3(Trimethyl); C21(Carbamidomethyl) | 2316.17722 | 0 | 0.00004089 | 1 | | | | | | | High | **50** | 0.005985893 | 3 | 772.73059 | 112.98 |
| MGLSNMDVEALKK | 20 | 1 | 1 | AL4G22860.t1 | M1(Oxidation); M6(Oxidation); K12(Trimethyl) | 1509.76545 | 0 | 0.000507 | 1 | High | **46** | 0.006761111 | 3 | 503.92667 | 45.27 | High | **36** | 0.073218605 | 3 | 503.92654 | 43.25 |

##### *A.halleri* / root samples

| | | | | | | | | | | CM | | | | | | Cd | | | | | |
|---|---|---|---|---|---|---|---|---|---|---|---|---|---|---|---|---|---|---|---|---|---|
| Sequence | # PSMs | # Proteins | # Protein Groups | Protein Group Accessions | Modifications | MH+ [Da] | q-Value | PEP | # Missed Cleavages | C2 | IonScore C2 | Exp Value C2 | Charge C2 | m/z [Da] C2 | RT C2 | F2 | IonScore F2 | Exp Value F2 | Charge F2 | m/z [Da] F2 | RT F2 |
| GITIDIALWKFETTK | 6 | 2 | 1 | AL1G18230.t1 | K10(Trimethyl) | 1778.01238 | 0 | 4.414E-18 | 1 | High | **66** | 6.95384E-05 | 2 | 889.51019 | 115.92 | High | **85** | 9.35094E-07 | 2 | 889.50983 | 115.94 |
| VGYNPDKIPFVPISGFEGDNMIER | 9 | 2 | 1 | AL1G18230.t1 | K7(Trimethyl); M21(Oxidation) | 2752.36448 | 0 | 4.937E-10 | 1 | High | **39** | 0.098815213 | 3 | 918.12695 | 114.09 | High | **49** | 0.008823305 | 3 | 918.12634 | 114.24 |
| KVGYNPDKIPFVPISGFEGDNMIER | 2 | 2 | 1 | AL1G18230.t1 | K1(Trimethyl) or K8(Trimethyl) | 2864.47207 | 0 | 0.0002889 | 2 | High | **37** | 0.155356965 | 4 | 716.87347 | 106.22 | High | 17 | 17.89656509 | 4 | 716.87280 | 106.54 |
| STNLDWYKGPTLLEALDQINEPK | 1 | 2 | 1 | AL1G18230.t1 | K23(Trimethyl) | 2687.38785 | 0 | 0.0001027 | 1 | High | **33** | 0.371887311 | 3 | 896.46747 | 121.05 | | | | | | |
| MGLENMDVESLKK | 11 | 1 | 1 | AL1G18770.t1 | M1(Oxidation); K12(Trimethyl) | 1551.77744 | 0 | 0.0001228 | 1 | High | 29 | 0.356202705 | 3 | 517.92969 | 61.39 | High | **34** | 0.123136688 | 3 | 517.93066 | 60.98 |
| MGLSNMDVEALKK | 11 | 1 | 1 | AL4G22860.t1 | K12(Trimethyl) | 1477.77527 | 0 | 1.004E-07 | 1 | High | **62** | 0.000197207 | 2 | 739.39050 | 69.56 | High | **48** | 0.006511349 | 2 | 739.39160 | 69.67 |
| MGLSNMDVEALKK | 94 | 1 | 1 | AL4G22860.t1 | M1(Oxidation); K12(Trimethyl) | 1493.76908 | 0 | 0.0000863 | 1 | High | **47** | 0.006272166 | 3 | 498.59479 | 59.93 | High | **49** | 0.003909218 | 3 | 498.59454 | 60.85 |
| MGLSNMDVEALKK | 39 | 1 | 1 | AL4G22860.t1 | M1(Oxidation); M6(Oxidation); K12(Trimethyl) | 1509.76535 | 0 | 0.0002944 | 1 | High | **45** | 0.008947476 | 3 | 503.92636 | 43.30 | High | **46** | 0.007327231 | 3 | 503.92664 | 44.09 |

##### *A.thaliana* / leaf samples

| | | | | | | | | | | CM | | | | | | Cd | | | | | |
|---|---|---|---|---|---|---|---|---|---|---|---|---|---|---|---|---|---|---|---|---|---|
| Sequence | # PSMs | # Proteins | # Protein Groups | Protein Group Accessions | Modifications | MH+ [Da] | q-Value | PEP | # Missed Cleavages | I2 | IonScore I2 | Exp Value I2 | Charge I2 | m/z [Da] I2 | RT I2 | L2 | IonScore L2 | Exp Value L2 | Charge L2 | m/z [Da] L2 | RT L2 |
| TTQFCSGGK | 1 | 1 | 1 | AT1G78820.1 | C5(Carbamidomethyl); K9(Trimethyl) | 1027.48747 | 0 | 0.01253 | 0 | | | | | | | High | **44** | 0.002123707 | 2 | 514.24738 | 34.64 |
| YTGEGESEEAKEGMFVK | 4 | 3 | 2 | AT2G21330.1;AT4G38970.1 | M14(Oxidation); K17(Trimethyl) | 1948.88449 | 0 | 0.00001006 | 1 | High | **44** | 0.005158432 | 3 | 650.29968 | 48.46 | | | | | | |

##### *A.lyrata* / leaf samples

| | | | | | | | | | | CM | | | | | | Cd | | | | | |
|---|---|---|---|---|---|---|---|---|---|---|---|---|---|---|---|---|---|---|---|---|---|
| Sequence | # PSMs | # Proteins | # Protein Groups | Protein Group Accessions | Modifications | MH+ [Da] | q-Value | PEP | # Missed Cleavages | C2 | IonScore C2 | Exp Value C2 | Charge C2 | m/z [Da] C2 | RT C2 | F2 | IonScore F2 | Exp Value F2 | Charge F2 | m/z [Da] F2 | RT F2 |
| VGYNPDKIPFVPISGFEGDNMIER | 5 | 2 | 1 | AL1G18230.t1 | K7(Trimethyl); M21(Oxidation) | 2752.37766 | 0 | 0.00115 | 1 | High | **31** | 0.584552891 | 3 | 918.12695 | 113.45 | High | **34** | 0.310275882 | 3 | 918.12653 | 113.56 |
| GITIDIALWKFETTK | 2 | 2 | 1 | AL1G18230.t1 | K10(Trimethyl) | 1778.01084 | 0 | 0.0001573 | 1 | High | **54** | 0.001232724 | 3 | 593.34180 | 115.95 | High | **49** | 0.004175102 | 3 | 593.34161 | 115.93 |
| MGLSNMDVEALKK | 5 | 1 | 1 | AL4G22860.t1 | M1(Oxidation); K12(Trimethyl) | 1493.76899 | 0 | 0.001164 | 1 | | | | | | | High | **40** | 0.027045181 | 3 | 498.59451 | 59.60 |

##### *A.halleri* / leaf samples

| | | | | | | | | | | CM | | | | | | Cd | | | | | |
|---|---|---|---|---|---|---|---|---|---|---|---|---|---|---|---|---|---|---|---|---|---|
| Sequence | # PSMs | # Proteins | # Protein Groups | Protein Group Accessions | Modifications | MH+ [Da] | q-Value | PEP | # Missed Cleavages | E2 | IonScore E2 | Exp Value E2 | Charge E2 | m/z [Da] E2 | RT E2 | K2 | IonScore K2 | Exp Value K2 | Charge K2 | m/z [Da] K2 | RT K2 |
| GITIDIALWKFETTK | 3 | 2 | 1 | AL1G18230.t1 | K10(Trimethyl) | 1778.01128 | 0 | 5.095E-14 | 1 | High | **35** | 0.091346455 | 3 | 593.34302 | 116.19 | High | **39** | 0.038962155 | 3 | 593.34241 | 116.20 |
| KVGYNPDKIPFVPISGFEGDNMIER | 4 | 2 | 1 | AL1G18230.t1 | K1(Trimethyl) or K8(Trimethyl) | 2864.46816 | 0 | 6.153E-08 | 2 | High | **32** | 0.559959442 | 3 | 955.48901 | 108.11 | High | **71** | 7.21281E-05 | 4 | 716.8725 | 107.84 |
| VGYNPDKIPFVPISGFEGDNMIER | 6 | 2 | 1 | AL1G18230.t1 | K7(Trimethyl) | 2736.37296 | 0 | 8.612E-10 | 1 | High | **50** | 0.008665759 | 3 | 912.79517 | 116.58 | High | **58** | 0.001207585 | 3 | 912.79584 | 116.53 |
| VGYNPDKIPFVPISGFEGDNMIER | 7 | 2 | 1 | AL1G18230.t1 | K7(Trimethyl); M21(Oxidation) | 2752.36448 | 0 | 2.875E-08 | 1 | High | **46** | 0.020450023 | 3 | 918.13025 | 113.34 | High | **51** | 0.006048272 | 3 | 918.12634 | 112.69 |
| NFEGLDLGKMDEANDSGLASYVAGQIDR | 2 | 2 | 2 | AL3G26800.t1 ;AL3G26790.t1 | K9(Trimethyl); M10(Oxidation); N14(Deamidated) | 3044.41861 | 0 | 1.489E-07 | 1 | High | **60** | 0.00044293 | 3 | 1015.47772 | 106.17 | High | **49** | 0.00793912 | 3 | 1015.48615 | 106.22 |
| ATSAGMKEQEAVNFLEK | 1 | 2 | 2 | AL3G43370.t1 ;AL7G44410.t1 | M6(Oxidation); K7(Trimethyl); Q9(Deamidated) | 1911.93564 | 0 | 0.0007097 | 1 | | | | | | | High | **37** | 0.063578037 | 3 | 637.9834 | 51.53 |
| YTGEGESEEAKEGMFVK | 2 | 2 | 2 | AL4G10470.t1 ;AL7G10870.t5 | K17(Trimethyl) | 1932.89005 | 0 | 0.000335 | 1 | High | **34** | 0.051446694 | 3 | 644.96558 | 55.24 | High | **37** | 0.036604011 | 3 | 644.96820 | 55.44 |
| MGLSNMDVEALKK | 17 | 1 | 1 | AL4G22860.t1 | K12(Trimethyl) | 1477.77861 | 0 | 0.0004045 | 1 | High | **42** | 0.021103396 | 2 | 739.39148 | 70.44 | High | **50** | 0.004023052 | 2 | 739.39185 | 70.32 |
| MGLSNMDVEALKK | 20 | 1 | 1 | AL4G22860.t1 | M1(Oxidation); K12(Trimethyl) | 1493.76750 | 0 | 0.00004872 | 1 | | | | | | | High | **62** | 0.000184478 | 2 | 747.38739 | 59.62 |
| KTPNSYMLQQFDNPANPK | 2 | 1 | 1 | AL4G42620.t1 | K1(Trimethyl); M7(Oxidation); Q9(Deamidated) | 2152.04203 | 0 | 0.008292 | 1 | High | **34** | 0.162569545 | 3 | 718.01886 | 50.66 | | | | | | |
| VENIVVIGHSACGGIKGLMSFPLDGNNSTDFIEDWVK | 1 | 3 | 3 | AL6G25520.t1 ;AL3G10670.t1 ;AL3G10670.t3 | C12(Carbamidomethyl); K16(Trimethyl); M19(Oxidation) | 4077.01235 | 0 | 0.00000783 | 1 | High | **36** | 0.232231028 | 4 | 1020.0085 | 122.40 | | | | | | |

### PSMs = number of peptide spectrum matches
### Proteins = number of proteins containing the peptide
### Protein Groups = number of different proteins containing the peptide
### Protein Group Accessions = accessions in TAIR or Alyrata_384_v2.1
Modifications = as identified by Mascot

MH+ = protonated peptide mass
q-Value = false discovery rate (Mascot/Percolator)
PEP = posterior error probability (Mascot/Percolator)
### Missed Cleavages = number of trypsin missed cleavages
X2 = confidence in peptide identification (Proteome discoverer)

IonScore X2 = peptide score (Mascot)
Exp value X2 = e-value (Mascot)
Charge X2 = number of charges
m/z [Da] X2 = peptide mass-to-charge ratio
RT X2 = retention time

**Supplemental Table S3:** Differentially expressed *KMT* genes in *A. thaliana* exposed to Cd.

| **Biological material** | **Cd concentration** | **Treatment duration** | **Coverage (# *KMT* genes)** | **Differentially expressed *KMT* genes** | **$Log_2$ fold change** | **Reference** |
|---|---|---|---|---|---|---|
| 4-week-old plants (hydroponics) | 5 µM | 2, 6, 30 hours | 59 | none | - | Herbette *et al*. 2006 |
| | 50 µM | 2, 6, 30 hours | | *SBS7* (30 hours in roots) | -1.11 | |
| 3-week-old plants (hydroponics) | 10 µM | 2 hours | 51 | none | - | Weber *et al*. 2006 |
| 3-week-old roots (hydroponics) | 200 µM | 6 hours | 51 | *SDG6*<br>*SDG34*<br>*PPKMT2*<br>*PTAC14*<br>*PPKMT3*<br>*SDG50*<br>*SDG18*<br>*SDG9*<br>*SBS9*<br>*SDG40*<br>*SBS7*<br>*SDG32*<br>*SDG5*<br>*SDG15*<br>*SDG13* | 1.72<br>1.67<br>1.40<br>1.36<br>1.20<br>1.19<br>1.17<br>1.16<br>1.09<br>1.07<br>1.04<br>1.02<br>-1.11<br>-1.23<br>-1.58 | Li *et al*. 2010 |
| 7-day-old seedlings (agar medium) | 200 µM | 6 hours | 51 | none | - | Jobe *et al*. 2012 |
| 2-week-old seedlings (agar medium) | 70 µM | 2 hours | 59 | *SDG29* | 1.01 | Khare *et al*. 2016 |
| 5-week-old roots (hydroponics) | 1 µM | 7 days | 51 | none | - | Fischer *et al*. 2017 |

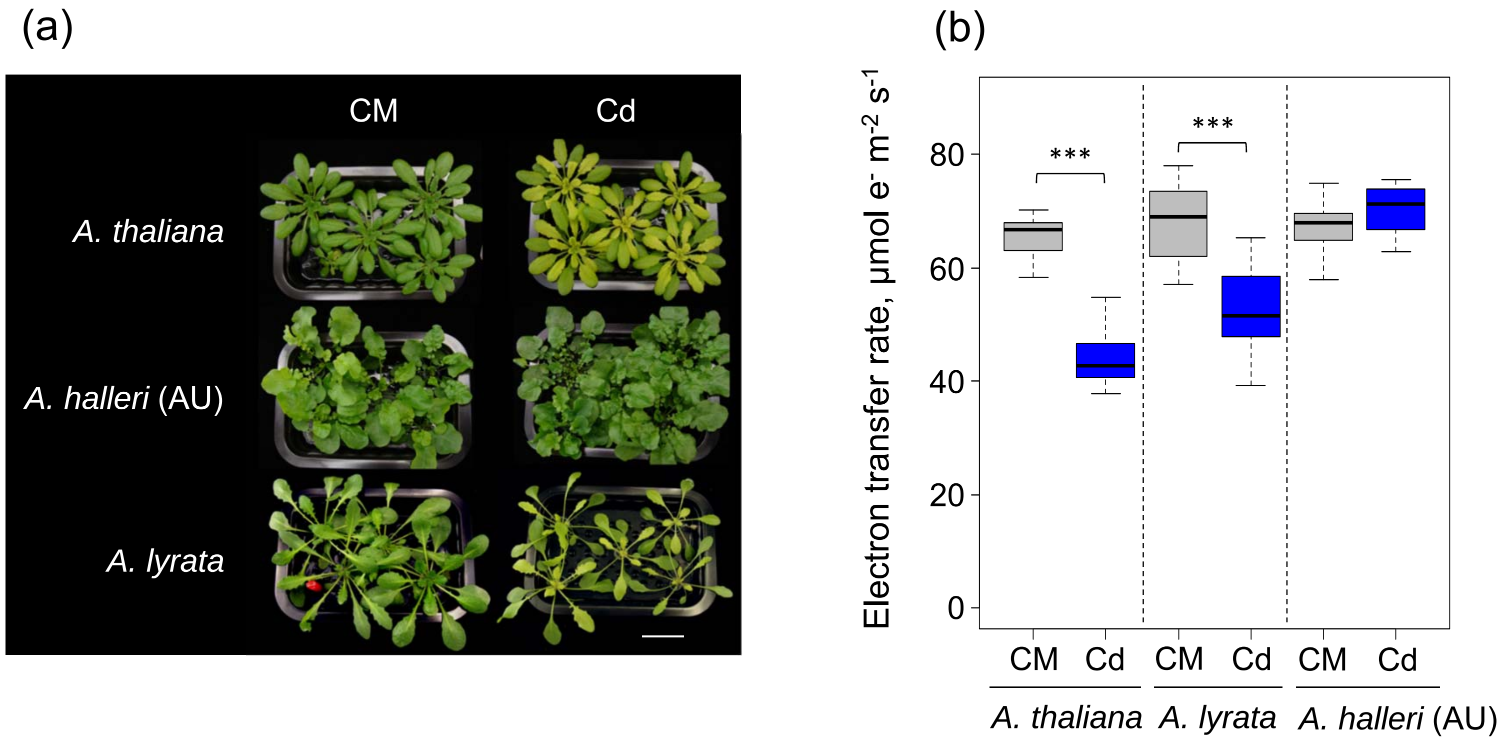


**Supplemental Figure S1**: Phenotype and photosynthesis of Arabidopsis plants challenged with Cd. *A. thaliana*, *A. lyrata*, and *A. halleri* (population Auby, AU) were grown hydroponically for five weeks in a standard culture medium and then challenged with 5 µM $CdSO_4$. Pictures (A) and photosynthesis measurements (B) were done after 9 days of treatment. Electron transfer rate was measured in young leaves at 300 µmol photons $m^{-2}$ $s^{-1}$. Each distribution in boxplots represents n = 20 measurements (5 plants, 4 leaves per plant). Statistical significance determined with Dunnett's test is shown, with $p < 0.001$ (***). CM, control medium; Cd, medium with 5 µM $CdSO_4$. Scale bar = 3 cm.

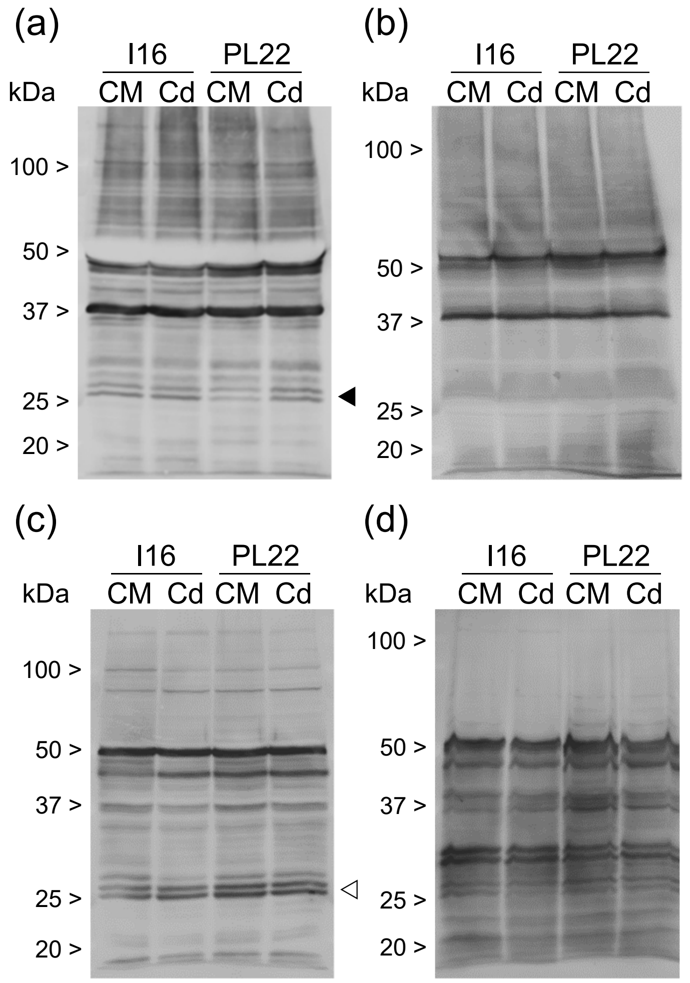


**Supplemental Figure S2:** Immunodetection of Lys-trimethylated proteins in roots and leaves from I16 and PL22 *A. halleri* plants challenged with Cd. Plants grown in hydropony were maintained in control medium (CM) or challenged with 5 µM $CdSO_4$ for 9 days. Soluble and membrane proteins were extracted from root and leaf tissues and analyzed by Western blot using antibodies specific to trimethyl-Lys. A - Leaf soluble proteins. B - Leaf membrane proteins. C - Root soluble proteins. D - Root membrane proteins. The triangles indicate the same protein doublets as in Fig. 1.

**Supplemental Figure S3** : LC-MS/MS fragmentation spectra of Lys trimethylated peptides identified in *A. thaliana*, *A. lyrata*, and *A. halleri* protein samples.

MS/MS fragmentation of ERGITIDIALWKFETTK found in AT1G07920 (GTP binding Elongation factor Tu family protein)

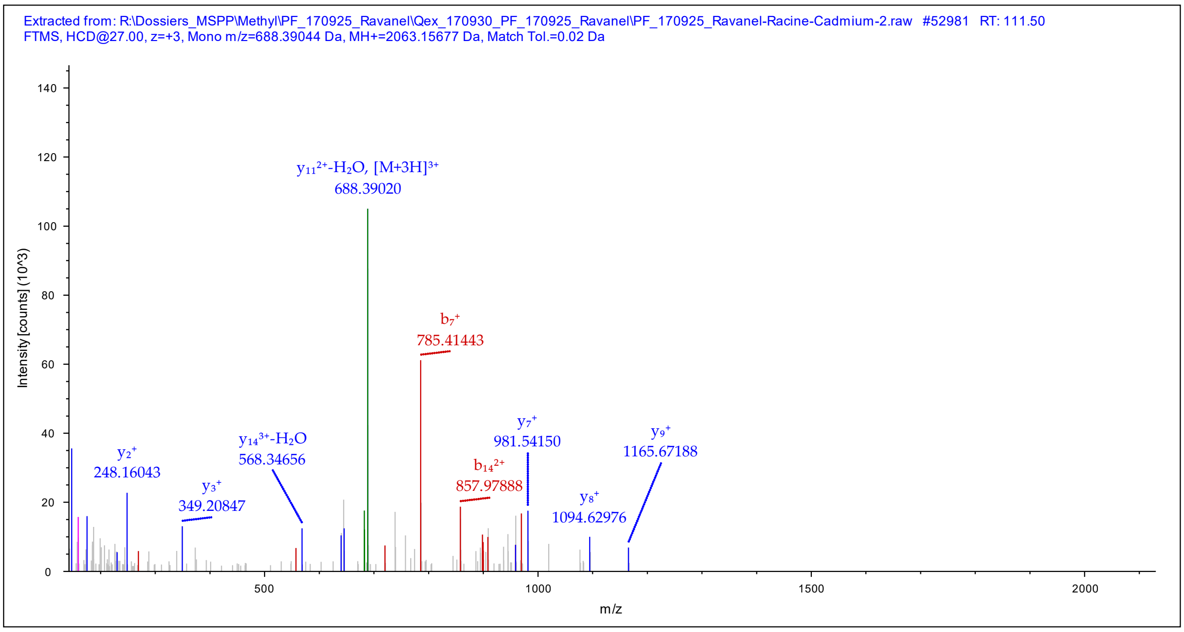


Peptide Summary

Sequence: ERGITIDIALWKFETTK, K12-Trimethyl (42.04695 Da)
Charge: +3, Monoisotopic m/z: 688.39044 Da (+1.04 mmu/+1.51 ppm), MH+: 2063.15677 Da, RT: 111.50 min,
Identified with: Mascot (v1.30); IonScore:38, Exp Value:1.5E-002, Ions matched by search engine: 6/188
Fragment match tolerance used for search: 20 mmu

Fragment Matches

Value Type: Theo. Mass [Da]

Ion Series | Neutral Losses | Precursor Ions

| #1 | Immonium | b+ | b2+ | b3+ | Seq. | y+ | y2+ | y3+ | #2 |
|---|---|---|---|---|---|---|---|---|---|
| 1 | 102.05496 | 130.04988 | 65.52858 | 44.02148 | E | | | | 17 |
| 2 | 129.11348 | 286.15100 | 143.57914 | 96.05518 | R | 1934.11105 | 967.55916 | 645.37520 | 16 |
| 3 | 30.03383 | 343.17247 | 172.08987 | 115.06234 | G | 1778.00993 | 889.50860 | 593.34149 | 15 |
| 4 | 86.09643 | 456.25654 | 228.63191 | 152.75703 | I | 1720.98846 | 860.99787 | 574.33434 | 14 |
| 5 | 74.06004 | 557.30422 | 279.15575 | 186.43959 | T | 1607.90439 | 804.45583 | 536.63965 | 13 |
| 6 | 86.09643 | 670.38829 | 335.69778 | 224.13428 | I | 1506.85671 | 753.93199 | 502.95709 | 12 |
| 7 | 88.03931 | 785.41524 | 393.21126 | 262.47660 | D | 1393.77264 | 697.38996 | 465.26240 | 11 |
| 8 | 86.09643 | 898.49931 | 449.75329 | 300.17129 | I | 1278.74569 | 639.87648 | 426.92008 | 10 |
| 9 | 44.04948 | 969.53643 | 485.27185 | 323.85033 | A | 1165.66162 | 583.33445 | 389.22539 | 9 |
| 10 | 86.09643 | 1082.62050 | 541.81389 | 361.54502 | L | 1094.62450 | 547.81589 | 365.54635 | 8 |
| 11 | 159.09168 | 1268.69982 | 634.85355 | 423.57146 | W | 981.54043 | 491.27385 | 327.85166 | 7 |
| 12 | 101.10733 | 1438.84174 | 719.92451 | 480.28543 | K-Trimethyl | 795.46111 | 398.23419 | 265.82522 | 6 |
| 13 | 120.08078 | 1585.91016 | 793.45872 | 529.30824 | F | 625.31919 | 313.16323 | 209.11125 | 5 |
| 14 | 102.05496 | 1714.95276 | 857.98002 | 572.32244 | E | 478.25077 | 239.62902 | 160.08844 | 4 |
| 15 | 74.06004 | 1816.00044 | 908.50386 | 606.00500 | T | 349.20817 | 175.10772 | 117.07424 | 3 |
| 16 | 74.06004 | 1917.04812 | 959.02770 | 639.68756 | T | 248.16049 | 124.58388 | 83.39168 | 2 |
| 17 | | | | | K | 147.11281 | 74.06004 | 49.70912 | 1 |

MS/MS fragmentation of GITIDIALWKFETTK found in AL1G18230 (GTP binding Elongation factor Tu family protein)

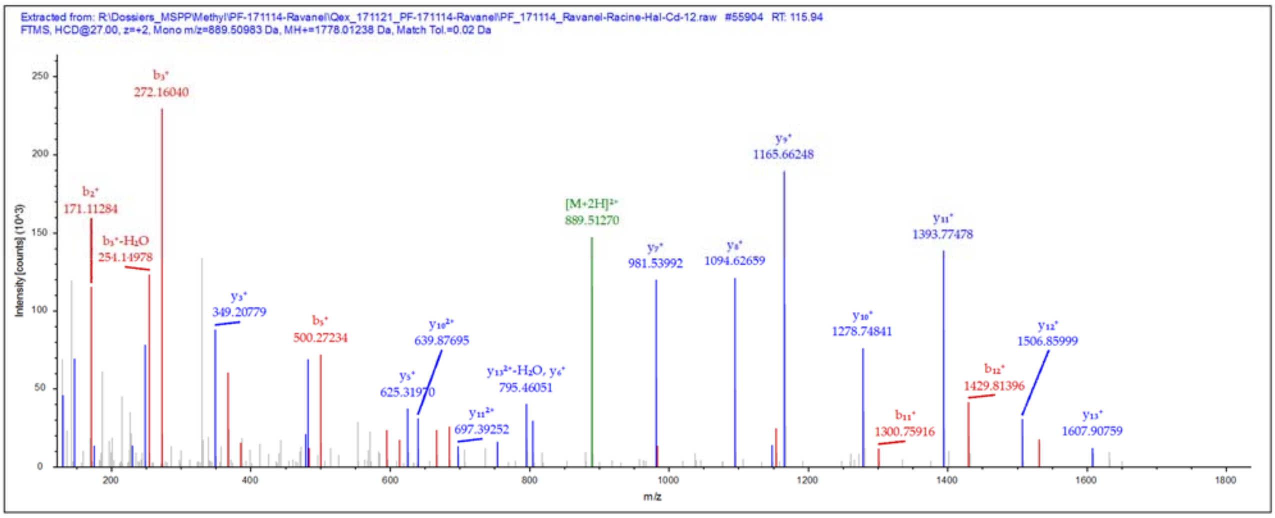


**Peptide Summary**

Sequence: GITIDIALWKFETTK, K10-Trimethyl (42.04695 Da)
Charge: +2, Monoisotopic m/z: 889.50983 Da (+1.22 mmu/+1.38 ppm), MH+: 1778.01238 Da, RT: 115.94 min,
Identified with: Mascot (v1.30); IonScore:85, Exp Value:1.7E-007, Ions matched by search engine: 13/132
Fragment match tolerance used for search: 20 mmu

**Fragment Matches**

Value Type: Theo. Mass [Da]

Ion Series | Neutral Losses | Precursor Ions

| #1 | Immonium | b⁺ | b²⁺ | Seq. | y⁺ | y²⁺ | #2 |
|---|---|---|---|---|---|---|---|
| 1 | 30.03383 | 58.02875 | 29.51801 | G | | | 15 |
| 2 | 86.09643 | 171.11282 | 86.06005 | I | 1720.98846 | 860.99787 | 14 |
| 3 | 74.06004 | 272.16050 | 136.58389 | T | 1607.90439 | 804.45583 | 13 |
| 4 | 86.09643 | 385.24457 | 193.12592 | I | 1506.85671 | 753.93199 | 12 |
| 5 | 88.03931 | 500.27152 | 250.63940 | D | 1393.77264 | 697.38996 | 11 |
| 6 | 86.09643 | 613.35559 | 307.18143 | I | 1278.74569 | 639.87648 | 10 |
| 7 | 44.04948 | 684.39271 | 342.69999 | A | 1165.66162 | 583.33445 | 9 |
| 8 | 86.09643 | 797.47678 | 399.24203 | L | 1094.62450 | 547.81589 | 8 |
| 9 | 159.09168 | 983.55610 | 492.28169 | W | 981.54043 | 491.27385 | 7 |
| 10 | 101.10733 | 1153.69802 | 577.35265 | K-Trimethyl | 795.46111 | 398.23419 | 6 |
| 11 | 120.08078 | 1300.76644 | 650.88686 | F | 625.31919 | 313.16323 | 5 |
| 12 | 102.05496 | 1429.80904 | 715.40816 | E | 478.25077 | 239.62902 | 4 |
| 13 | 74.06004 | 1530.85672 | 765.93200 | T | 349.20817 | 175.10772 | 3 |
| 14 | 74.06004 | 1631.90440 | 816.45584 | T | 248.16049 | 124.58388 | 2 |
| 15 | | | | K | 147.11281 | 74.06004 | 1 |

MS/MS fragmentation of VGYNPDKIPFVPISGFEGDNMIER found in AL1G18230 (GTP binding Elongation factor Tu family protein)

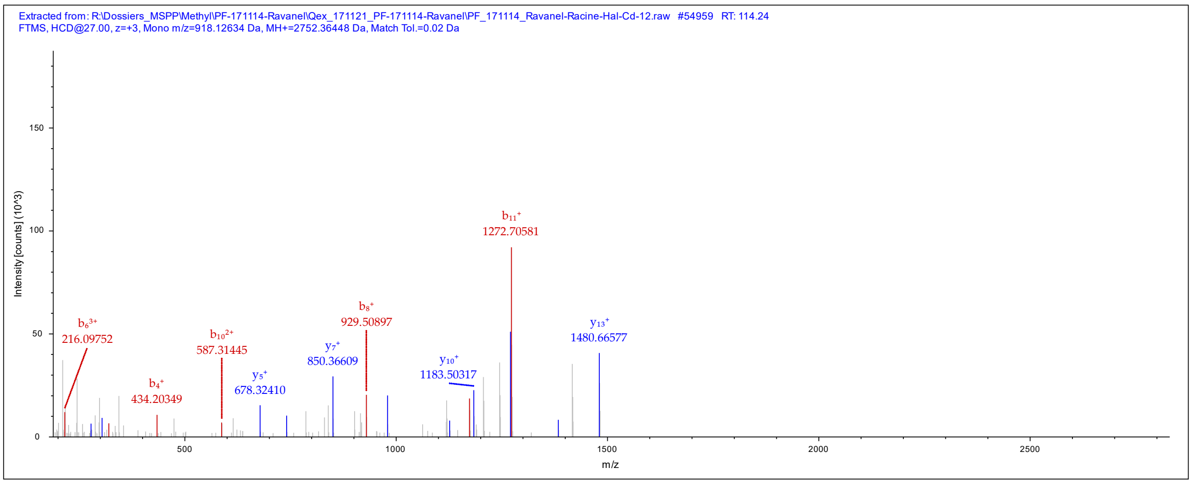


**Peptide Summary**

Sequence: VGYNPDKIPFVPISGFEGDNMIER, K7-Trimethyl (42.04695 Da), M21-Oxidation (15.99492 Da)
Charge: +3, Monoisotopic m/z: 918.12634 Da (-0.25 mmu/-0.27 ppm), MH+: 2752.36448 Da, RT: 114.24 min,
Identified with: Mascot (v1.30); IonScore:49, Exp Value:1.8E-003, Ions matched by search engine: 18/264
Fragment match tolerance used for search: 20 mmu

**Fragment Matches**

Value Type: Theo. Mass [Da]

Ion Series | Neutral Losses | Precursor Ions

| #1 | Immonium | $b^+$ | $b^{2+}$ | $b^{3+}$ | Seq. | $y^+$ | $y^{2+}$ | $y^{3+}$ | #2 |
|---|---|---|---|---|---|---|---|---|---|
| 1 | 72.08078 | 100.07570 | 50.54149 | 34.03008 | V | | | | 24 |
| 2 | 30.03383 | 157.09717 | 79.05222 | 53.03724 | G | 2653.29680 | 1327.15204 | 885.10378 | 23 |
| 3 | 136.07568 | 320.16049 | 160.58388 | 107.39168 | Y | 2596.27533 | 1298.64130 | 866.09663 | 22 |
| 4 | 87.05529 | 434.20342 | 217.60535 | 145.40599 | N | 2433.21201 | 1217.10964 | 811.74219 | 21 |
| 5 | 70.06513 | 531.25619 | 266.13173 | 177.75691 | P | 2319.16908 | 1160.08818 | 773.72788 | 20 |
| 6 | 88.03931 | 646.28314 | 323.64521 | 216.09923 | D | 2222.11631 | 1111.56179 | 741.37695 | 19 |
| 7 | 101.10733 | 816.42506 | 408.71617 | 272.81320 | K-Trimethyl | 2107.08936 | 1054.04832 | 703.03464 | 18 |
| 8 | 86.09643 | 929.50913 | 465.25820 | 310.50789 | I | 1936.94744 | 968.97736 | 646.32066 | 17 |
| 9 | 70.06513 | 1026.56190 | 513.78459 | 342.85882 | P | 1823.86337 | 912.43532 | 608.62597 | 16 |
| 10 | 120.08078 | 1173.63032 | 587.31880 | 391.88162 | F | 1726.81060 | 863.90894 | 576.27505 | 15 |
| 11 | 72.08078 | 1272.69874 | 636.85301 | 424.90443 | V | 1579.74218 | 790.37473 | 527.25224 | 14 |
| 12 | 70.06513 | 1369.75151 | 685.37939 | 457.25535 | P | 1480.67376 | 740.84052 | 494.22944 | 13 |
| 13 | 86.09643 | 1482.83558 | 741.92143 | 494.95004 | I | 1383.62099 | 692.31413 | 461.87851 | 12 |
| 14 | 60.04439 | 1569.86761 | 785.43744 | 523.96072 | S | 1270.53692 | 635.77210 | 424.18382 | 11 |
| 15 | 30.03383 | 1626.88908 | 813.94818 | 542.96788 | G | 1183.50489 | 592.25608 | 395.17315 | 10 |
| 16 | 120.08078 | 1773.95750 | 887.48239 | 591.99068 | F | 1126.48342 | 563.74535 | 376.16599 | 9 |
| 17 | 102.05496 | 1903.00010 | 952.00369 | 635.00488 | E | 979.41500 | 490.21114 | 327.14318 | 8 |
| 18 | 30.03383 | 1960.02157 | 980.51442 | 654.01204 | G | 850.37240 | 425.68984 | 284.12898 | 7 |
| 19 | 88.03931 | 2075.04852 | 1038.02790 | 692.35436 | D | 793.35093 | 397.17910 | 265.12183 | 6 |
| 20 | 87.05529 | 2189.09145 | 1095.04936 | 730.36867 | N | 678.32398 | 339.66563 | 226.77951 | 5 |
| 21 | 104.05286 | 2336.12686 | 1168.56707 | 779.38047 | M-Oxidation | 564.28105 | 282.64416 | 188.76520 | 4 |
| 22 | 86.09643 | 2449.21093 | 1225.10910 | 817.07516 | I | 417.24563 | 209.12645 | 139.75339 | 3 |
| 23 | 102.05496 | 2578.25353 | 1289.63040 | 860.08936 | E | 304.16156 | 152.58442 | 102.05870 | 2 |
| 24 | | | | | R | 175.11896 | 88.06312 | 59.04450 | 1 |

MS/MS fragmentation of KVGYNPDKIPFVPISGFEGDNMIER found in AT1G07920 (GTP binding Elongation factor Tu family protein)

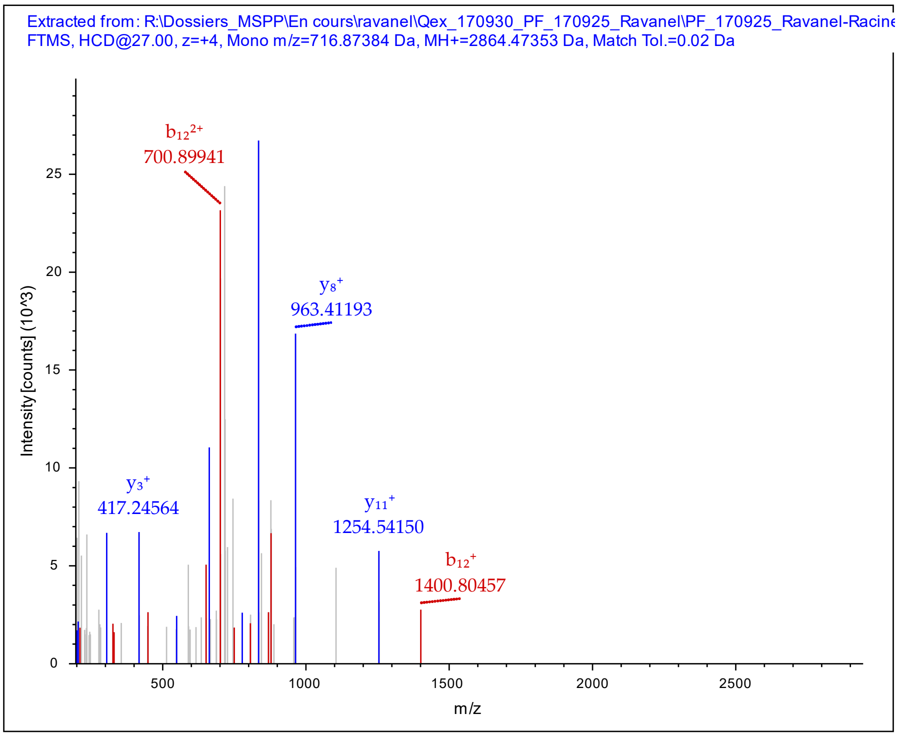


Peptide Summary

Sequence: KVGYNPDKIPFVPISGFEGDNMIER, K8-Trimethyl (42.04695 Da)
Charge: +4, Monoisotopic m/z: 716.87384 Da (+2.07 mmu/+2.88 ppm), MH+: 2864.47353 Da, RT: 112.49 min,
Identified with: Mascot (v1.30); IonScore:33, Exp Value:5.2E-001, Ions matched by search engine: 7/288
Fragment match tolerance used for search: 20 mmu

Fragment Matches

Value Type: Theo. Mass [Da]

Ion Series | Neutral Losses | Precursor Ions

| #1 | Immonium | $b^+$ | $b^{2+}$ | $b^{3+}$ | $b^{4+}$ | Seq. | $y^+$ | $y^{2+}$ | $y^{3+}$ | $y^{4+}$ | #2 |
|---|---|---|---|---|---|---|---|---|---|---|---|
| 1 | 101.10733 | 129.10225 | 65.05476 | 43.70560 | 33.03102 | K | | | | | 25 |
| 2 | 72.08078 | 228.17067 | 114.58897 | 76.72841 | 57.79812 | V | 2736.37030 | 1368.68879 | 912.79495 | 684.84803 | 24 |
| 3 | 30.03383 | 285.19214 | 143.09971 | 95.73556 | 72.05349 | G | 2637.30188 | 1319.15458 | 879.77214 | 660.08093 | 23 |
| 4 | 136.07568 | 448.25546 | 224.63137 | 150.09000 | 112.81932 | Y | 2580.28041 | 1290.64384 | 860.76499 | 645.82556 | 22 |
| 5 | 87.05529 | 562.29839 | 281.65283 | 188.10431 | 141.33005 | N | 2417.21709 | 1209.11218 | 806.41055 | 605.05973 | 21 |
| 6 | 70.06513 | 659.35116 | 330.17922 | 220.45524 | 165.59325 | P | 2303.17416 | 1152.09072 | 768.39624 | 576.54900 | 20 |
| 7 | 88.03931 | 774.37811 | 387.69269 | 258.79755 | 194.34998 | D | 2206.12139 | 1103.56433 | 736.04531 | 552.28581 | 19 |
| 8 | 101.10733 | 944.52003 | 472.76365 | 315.51153 | 236.88546 | K-Trimethyl | 2091.09444 | 1046.05086 | 697.70300 | 523.52907 | 18 |
| 9 | 86.09643 | 1057.60410 | 529.30569 | 353.20622 | 265.15648 | I | 1920.95252 | 960.97990 | 640.98902 | 480.99359 | 17 |
| 10 | 70.06513 | 1154.65687 | 577.83207 | 385.55714 | 289.41967 | P | 1807.86845 | 904.43786 | 603.29433 | 452.72257 | 16 |
| 11 | 120.08078 | 1301.72529 | 651.36628 | 434.57995 | 326.18678 | F | 1710.81568 | 855.91148 | 570.94341 | 428.45938 | 15 |
| 12 | 72.08078 | 1400.79371 | 700.90049 | 467.60275 | 350.95388 | V | 1563.74726 | 782.37727 | 521.92060 | 391.69227 | 14 |
| 13 | 70.06513 | 1497.84648 | 749.42688 | 499.95368 | 375.21708 | P | 1464.67884 | 732.84306 | 488.89780 | 366.92517 | 13 |
| 14 | 86.09643 | 1610.93055 | 805.96891 | 537.64837 | 403.48809 | I | 1367.62607 | 684.31667 | 456.54687 | 342.66198 | 12 |
| 15 | 60.04439 | 1697.96258 | 849.48493 | 566.65904 | 425.24610 | S | 1254.54200 | 627.77464 | 418.85218 | 314.39096 | 11 |
| 16 | 30.03383 | 1754.98405 | 877.99566 | 585.66620 | 439.50147 | G | 1167.50997 | 584.25862 | 389.84151 | 292.63295 | 10 |
| 17 | 120.08078 | 1902.05247 | 951.52987 | 634.68901 | 476.26857 | F | 1110.48850 | 555.74789 | 370.83435 | 278.37758 | 9 |
| 18 | 102.05496 | 2031.09507 | 1016.05117 | 677.70321 | 508.52922 | E | 963.42008 | 482.21368 | 321.81154 | 241.61048 | 8 |
| 19 | 30.03383 | 2088.11654 | 1044.56191 | 696.71036 | 522.78459 | G | 834.37748 | 417.69238 | 278.79734 | 209.34983 | 7 |
| 20 | 88.03931 | 2203.14349 | 1102.07538 | 735.05268 | 551.54133 | D | 777.35601 | 389.18164 | 259.79019 | 195.09446 | 6 |
| 21 | 87.05529 | 2317.18642 | 1159.09685 | 773.06699 | 580.05206 | N | 662.32906 | 331.66817 | 221.44787 | 166.33772 | 5 |
| 22 | 104.05286 | 2448.22692 | 1224.61710 | 816.74716 | 612.81219 | M | 548.28613 | 274.64670 | 183.43356 | 137.82699 | 4 |
| 23 | 86.09643 | 2561.31099 | 1281.15913 | 854.44185 | 641.08320 | I | 417.24563 | 209.12645 | 139.75339 | 105.06687 | 3 |
| 24 | 102.05496 | 2690.35359 | 1345.68043 | 897.45605 | 673.34385 | E | 304.16156 | 152.58442 | 102.05870 | 76.79585 | 2 |
| 25 | | | | | | R | 175.11896 | 88.06312 | 59.04450 | 44.53520 | 1 |

MS/MS fragmentation of GPTLLEALDQINEPK found in AT1G07920 (GTP binding Elongation factor Tu family protein)

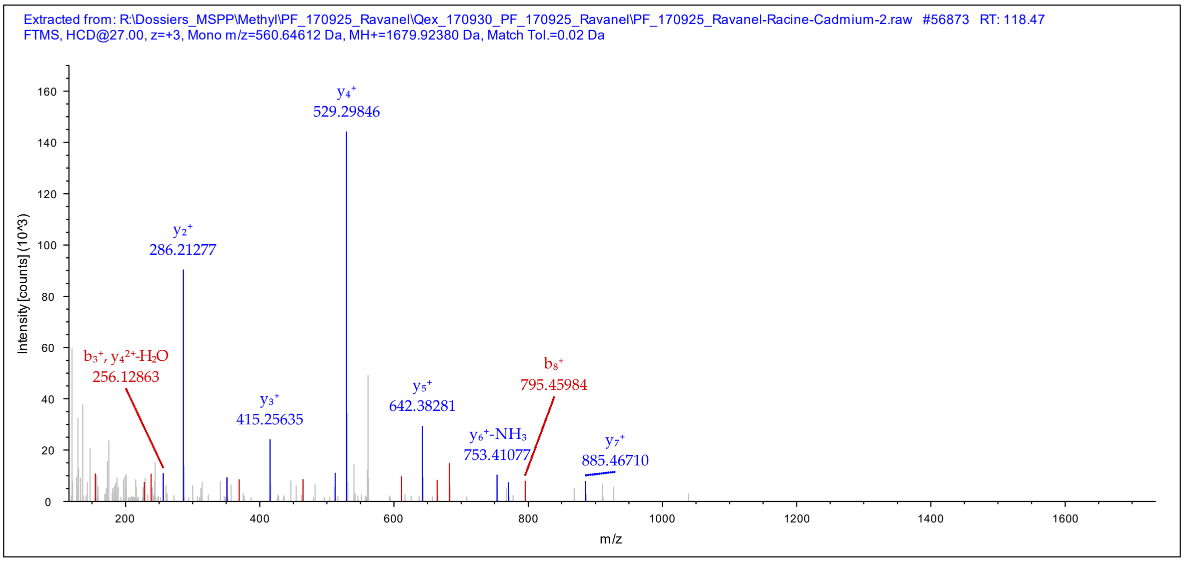


**Peptide Summary**

Sequence: GPTLLEALDQINEPK, K15-Trimethyl (42.04695 Da)
Charge: +3, Monoisotopic m/z: 560.64612 Da (+0.76 mmu/+1.36 ppm), MH+: 1679.92380 Da, RT: 118.47 min,
Identified with: Mascot (v1.30); IonScore:45, Exp Value:2.7E-003, Ions matched by search engine: 14/132
Fragment match tolerance used for search: 20 mmu

**Fragment Matches**

Value Type: Theo. Mass [Da]

Ion Series | Neutral Losses | Precursor Ions

| #1 | Immonium | $b^+$ | $b^{2+}$ | $b^{3+}$ | Seq. | $y^+$ | $y^{2+}$ | $y^{3+}$ | #2 |
|---|---|---|---|---|---|---|---|---|---|
| 1 | 30.03383 | 58.02875 | 29.51801 | 20.01443 | G | | | | 15 |
| 2 | 70.06513 | 155.08152 | 78.04440 | 52.36536 | P | 1622.90004 | 811.95366 | 541.63820 | 14 |
| 3 | 74.06004 | 256.12920 | 128.56824 | 86.04792 | T | 1525.84727 | 763.42727 | 509.28727 | 13 |
| 4 | 86.09643 | 369.21327 | 185.11027 | 123.74261 | L | 1424.79959 | 712.90343 | 475.60471 | 12 |
| 5 | 86.09643 | 482.29734 | 241.65231 | 161.43730 | L | 1311.71552 | 656.36140 | 437.91002 | 11 |
| 6 | 102.05496 | 611.33994 | 306.17361 | 204.45150 | E | 1198.63145 | 599.81936 | 400.21533 | 10 |
| 7 | 44.04948 | 682.37706 | 341.69217 | 228.13054 | A | 1069.58885 | 535.29806 | 357.20113 | 9 |
| 8 | 86.09643 | 795.46113 | 398.23420 | 265.82523 | L | 998.55173 | 499.77950 | 333.52209 | 8 |
| 9 | 88.03931 | 910.48808 | 455.74768 | 304.16754 | D | 885.46766 | 443.23747 | 295.82740 | 7 |
| 10 | 101.07094 | 1038.54666 | 519.77697 | 346.85374 | Q | 770.44071 | 385.72399 | 257.48509 | 6 |
| 11 | 86.09643 | 1151.63073 | 576.31900 | 384.54843 | I | 642.38213 | 321.69470 | 214.79889 | 5 |
| 12 | 87.05529 | 1265.67366 | 633.34047 | 422.56274 | N | 529.29806 | 265.15267 | 177.10420 | 4 |
| 13 | 102.05496 | 1394.71626 | 697.86177 | 465.57694 | E | 415.25513 | 208.13120 | 139.08989 | 3 |
| 14 | 70.06513 | 1491.76903 | 746.38815 | 497.92786 | P | 286.21253 | 143.60990 | 96.07569 | 2 |
| 15 | | | | | K-Trimethyl | 189.15976 | 95.08352 | 63.72477 | 1 |

MS/MS fragmentation of STNLDWYKGPTLLEALDQINEPK found in AL1G18230 (GTP binding Elongation factor Tu family protein)

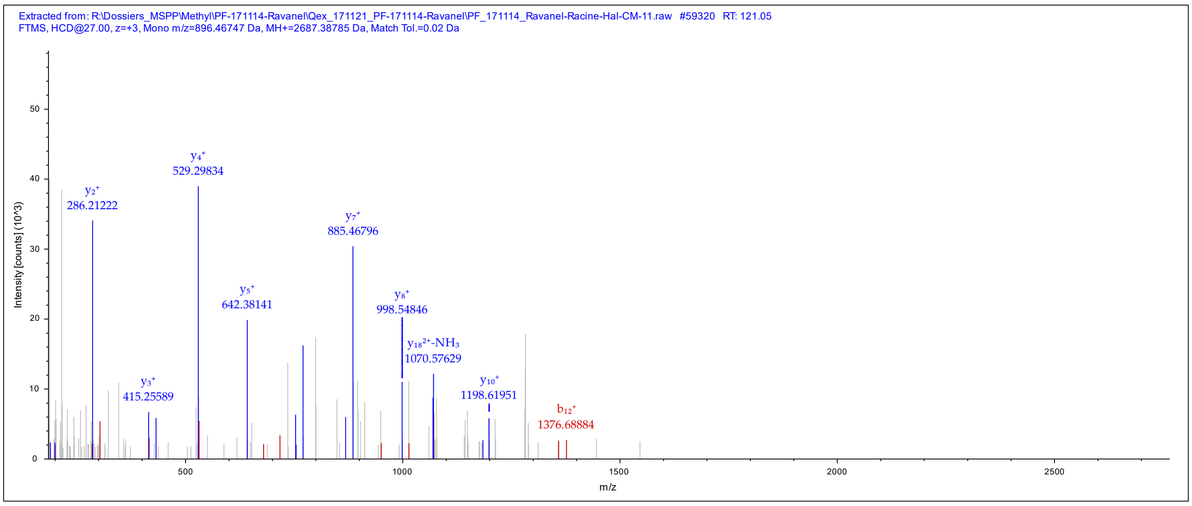


**Peptide Summary**

Sequence: STNLDWYKGPTLLEALDQINEPK, K23-Trimethyl (42.04695 Da)
Charge: +3, Monoisotopic m/z: 896.46747 Da (-1.64 mmu/-1.83 ppm), MH+: 2687.38785 Da, RT: 121.05 min,
Identified with: Mascot (v1.30); IonScore:33, Exp Value:7.1E-002, Ions matched by search engine: 14/256
Fragment match tolerance used for search: 20 mmu

**Fragment Matches**

Value Type: Theo. Mass [Da]

Ion Series | Neutral Losses | Precursor Ions

| #1 | Immonium | b+ | b2+ | b3+ | Seq. | y+ | y2+ | y3+ | #2 |
|---|---|---|---|---|---|---|---|---|---|
| 1 | 60.04439 | 88.03931 | 44.52329 | 30.01795 | S | | | | 23 |
| 2 | 74.06004 | 189.08699 | 95.04713 | 63.70051 | T | 2600.36075 | 1300.68401 | 867.45843 | 22 |
| 3 | 87.05529 | 303.12992 | 152.06860 | 101.71482 | N | 2499.31307 | 1250.16017 | 833.77587 | 21 |
| 4 | 86.09643 | 416.21399 | 208.61063 | 139.40951 | L | 2385.27014 | 1193.13871 | 795.76156 | 20 |
| 5 | 88.03931 | 531.24094 | 266.12411 | 177.75183 | D | 2272.18607 | 1136.59667 | 758.06687 | 19 |
| 6 | 159.09168 | 717.32026 | 359.16377 | 239.77827 | W | 2157.15912 | 1079.08320 | 719.72456 | 18 |
| 7 | 136.07568 | 880.38358 | 440.69543 | 294.13271 | Y | 1971.07980 | 986.04354 | 657.69812 | 17 |
| 8 | 101.10733 | 1008.47855 | 504.74291 | 336.83103 | K | 1808.01648 | 904.51188 | 603.34368 | 16 |
| 9 | 30.03383 | 1065.50002 | 533.25365 | 355.83819 | G | 1679.92151 | 840.46439 | 560.64535 | 15 |
| 10 | 70.06513 | 1162.55279 | 581.78003 | 388.18911 | P | 1622.90004 | 811.95366 | 541.63820 | 14 |
| 11 | 74.06004 | 1263.60047 | 632.30387 | 421.87167 | T | 1525.84727 | 763.42727 | 509.28727 | 13 |
| 12 | 86.09643 | 1376.68454 | 688.84591 | 459.56636 | L | 1424.79959 | 712.90343 | 475.60471 | 12 |
| 13 | 86.09643 | 1489.76861 | 745.38794 | 497.26105 | L | 1311.71552 | 656.36140 | 437.91002 | 11 |
| 14 | 102.05496 | 1618.81121 | 809.90924 | 540.27525 | E | 1198.63145 | 599.81936 | 400.21533 | 10 |
| 15 | 44.04948 | 1689.84833 | 845.42780 | 563.95429 | A | 1069.58885 | 535.29806 | 357.20113 | 9 |
| 16 | 86.09643 | 1802.93240 | 901.96984 | 601.64898 | L | 998.55173 | 499.77950 | 333.52209 | 8 |
| 17 | 88.03931 | 1917.95935 | 959.48331 | 639.99130 | D | 885.46766 | 443.23747 | 295.82740 | 7 |
| 18 | 101.07094 | 2046.01793 | 1023.51260 | 682.67749 | Q | 770.44071 | 385.72399 | 257.48509 | 6 |
| 19 | 86.09643 | 2159.10200 | 1080.05464 | 720.37218 | I | 642.38213 | 321.69470 | 214.79889 | 5 |
| 20 | 87.05529 | 2273.14493 | 1137.07610 | 758.38649 | N | 529.29806 | 265.15267 | 177.10420 | 4 |
| 21 | 102.05496 | 2402.18753 | 1201.59740 | 801.40069 | E | 415.25513 | 208.13120 | 139.08989 | 3 |
| 22 | 70.06513 | 2499.24030 | 1250.12379 | 833.75162 | P | 286.21253 | 143.60990 | 96.07569 | 2 |
| 23 | | | | | K-Trimethyl | 189.15976 | 95.08352 | 63.72477 | 1 |

MS/MS fragmentation of NYDPQKDKR found in AT1G08360 (Ribosomal protein L1p/L10e family)

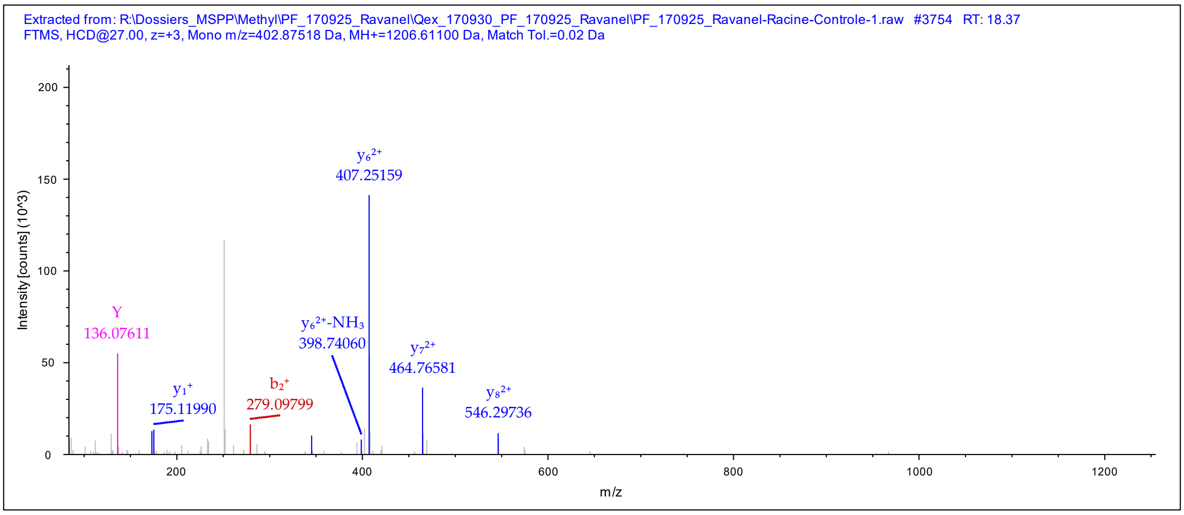


#### Peptide Summary

Sequence: NYDPQKDKR, N1-Deamidated (0.98402 Da), K8-Trimethyl (42.04695 Da)
Charge: +3, Monoisotopic m/z: 402.87518 Da (-0.12 mmu/-0.31 ppm), MH+: 1206.61100 Da, RT: 18.37 min,
Identified with: Mascot (v1.30); IonScore:31, Exp Value:2.6E-002, Ions matched by search engine: 5/96
Fragment match tolerance used for search: 20 mmu

#### Fragment Matches

Value Type: Theo. Mass [Da]

Ion Series | Neutral Losses | Precursor Ions

| #1 | Immonium | $b^+$ | $b^{2+}$ | $b^{3+}$ | Seq. | $y^+$ | $y^{2+}$ | $y^{3+}$ | #2 |
|---|---|---|---|---|---|---|---|---|---|
| 1 | 87.05529 | 116.03422 | 58.52075 | 39.34959 | N-Deamid... | | | | 9 |
| 2 | 136.07568 | 279.09754 | 140.05241 | 93.70403 | Y | 1091.58442 | 546.29585 | 364.53299 | 8 |
| 3 | 88.03931 | 394.12449 | 197.56588 | 132.04635 | D | 928.52110 | 464.76419 | 310.17855 | 7 |
| 4 | 70.06513 | 491.17726 | 246.09227 | 164.39727 | P | 813.49415 | 407.25071 | 271.83623 | 6 |
| 5 | 101.07094 | 619.23584 | 310.12156 | 207.08347 | Q | 716.44138 | 358.72433 | 239.48531 | 5 |
| 6 | 101.10733 | 747.33081 | 374.16904 | 249.78179 | K | 588.38280 | 294.69504 | 196.79912 | 4 |
| 7 | 88.03931 | 862.35776 | 431.68252 | 288.12411 | D | 460.28783 | 230.64755 | 154.10079 | 3 |
| 8 | 101.10733 | 1032.49968 | 516.75348 | 344.83808 | K-Trimethyl | 345.26088 | 173.13408 | 115.75848 | 2 |
| 9 | | | | | R | 175.11896 | 88.06312 | 59.04450 | 1 |

MS/MS fragmentation of MGLENMDVESLKK found in AT1G08360 (Ribosomal protein L1p/L10e family)

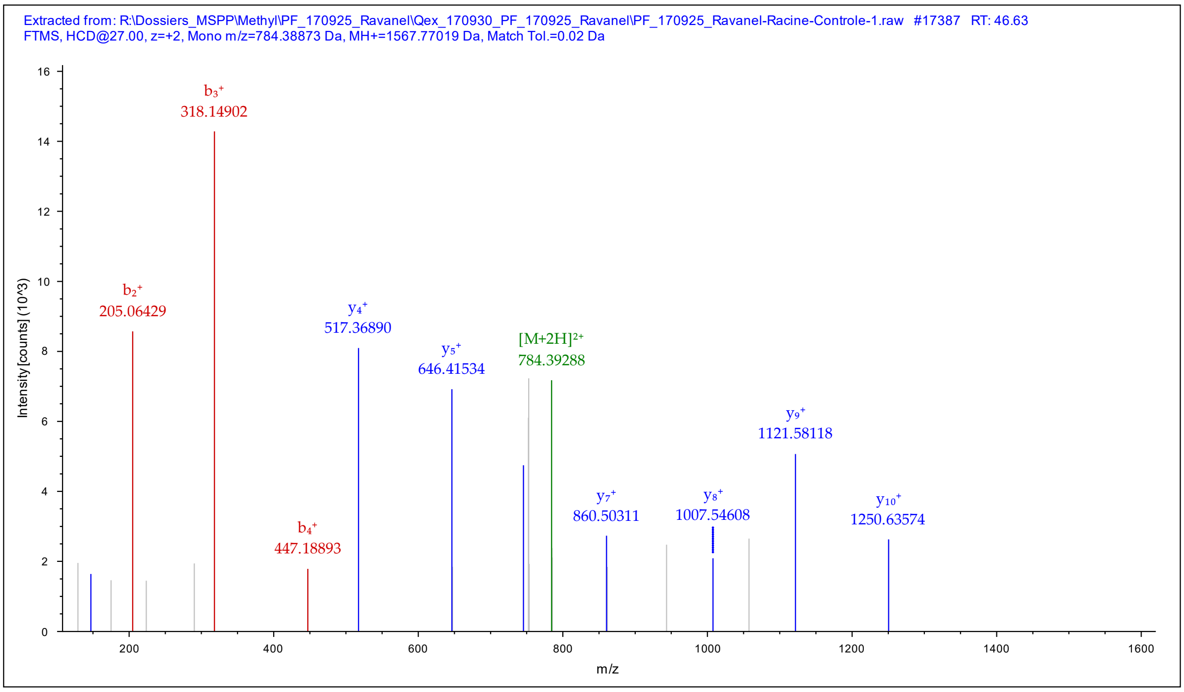


#### Peptide Summary

Sequence: MGLENMDVESLKK, M1-Oxidation (15.99492 Da), M6-Oxidation (15.99492 Da), K12-Trimethyl (42.04695 Da)
Charge: +2, Monoisotopic m/z: 784.38873 Da (-0.26 mmu/-0.33 ppm), MH+: 1567.77019 Da, RT: 46.63 min,
Identified with: Mascot (v1.30); IonScore:62, Exp Value:4.3E-005, Ions matched by search engine: 8/128
Fragment match tolerance used for search: 20 mmu

#### Fragment Matches

Value Type: Theo. Mass [Da]

Ion Series | Neutral Losses | Precursor Ions

| #1 | Immonium | b+ | b2+ | Seq. | y+ | y2+ | #2 |
|---|---|---|---|---|---|---|---|
| 1 | 104.05286 | 148.04269 | 74.52498 | M-Oxidation | | | 13 |
| 2 | 30.03383 | 205.06416 | 103.03572 | G | 1420.73529 | 710.87128 | 12 |
| 3 | 86.09643 | 318.14823 | 159.57775 | L | 1363.71382 | 682.36055 | 11 |
| 4 | 102.05496 | 447.19083 | 224.09905 | E | 1250.62975 | 625.81851 | 10 |
| 5 | 87.05529 | 561.23376 | 281.12052 | N | 1121.58715 | 561.29721 | 9 |
| 6 | 104.05286 | 708.26918 | 354.63823 | M-Oxidation | 1007.54422 | 504.27575 | 8 |
| 7 | 88.03931 | 823.29613 | 412.15170 | D | 860.50880 | 430.75804 | 7 |
| 8 | 72.08078 | 922.36455 | 461.68591 | V | 745.48185 | 373.24456 | 6 |
| 9 | 102.05496 | 1051.40715 | 526.20721 | E | 646.41343 | 323.71035 | 5 |
| 10 | 60.04439 | 1138.43918 | 569.72323 | S | 517.37083 | 259.18905 | 4 |
| 11 | 86.09643 | 1251.52325 | 626.26526 | L | 430.33880 | 215.67304 | 3 |
| 12 | 101.10733 | 1421.66517 | 711.33622 | K-Trimethyl | 317.25473 | 159.13100 | 2 |
| 13 | | | | K | 147.11281 | 74.06004 | 1 |

MS/MS fragmentation of MGLSNMDVEALKK found in AL4G22860 (Ribosomal protein L1p/L10e family)

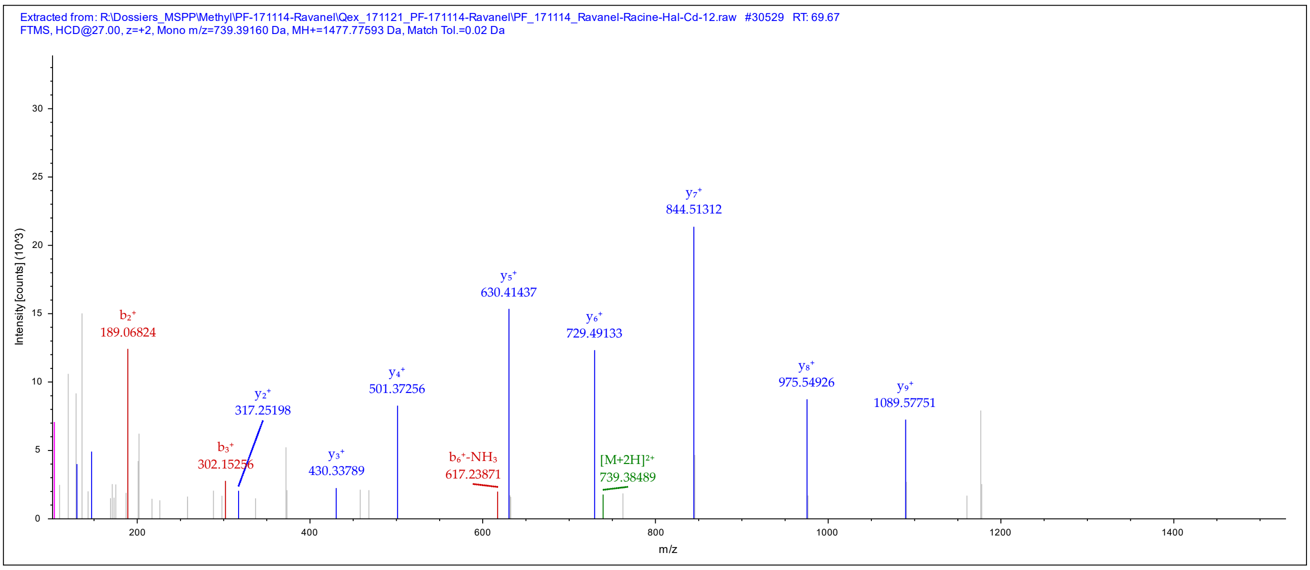


Peptide Summary

Sequence: MGLSNMDVEALKK, K12-Trimethyl (42.04695 Da)
Charge: +2, Monoisotopic m/z: 739.39160 Da (+0.27 mmu/+0.36 ppm), MH+: 1477.77593 Da, RT: 69.67 min,
Identified with: Mascot (v1.30); IonScore:48, Exp Value:1.3E-003, Ions matched by search engine: 9/128
Fragment match tolerance used for search: 20 mmu

Fragment Matches

Value Type: Theo. Mass [Da]

Ion Series | Neutral Losses | Precursor Ions

| #1 | Immonium | $b^+$ | $b^{2+}$ | Seq. | $y^+$ | $y^{2+}$ | #2 |
|---|---|---|---|---|---|---|---|
| 1 | 104.05286 | 132.04778 | 66.52753 | M | | | 13 |
| 2 | 30.03383 | 189.06925 | 95.03826 | G | 1346.73489 | 673.87108 | 12 |
| 3 | 86.09643 | 302.15332 | 151.58030 | L | 1289.71342 | 645.36035 | 11 |
| 4 | 60.04439 | 389.18535 | 195.09631 | S | 1176.62935 | 588.81831 | 10 |
| 5 | 87.05529 | 503.22828 | 252.11778 | N | 1089.59732 | 545.30230 | 9 |
| 6 | 104.05286 | 634.26878 | 317.63803 | M | 975.55439 | 488.28083 | 8 |
| 7 | 88.03931 | 749.29573 | 375.15150 | D | 844.51389 | 422.76058 | 7 |
| 8 | 72.08078 | 848.36415 | 424.68571 | V | 729.48694 | 365.24711 | 6 |
| 9 | 102.05496 | 977.40675 | 489.20701 | E | 630.41852 | 315.71290 | 5 |
| 10 | 44.04948 | 1048.44387 | 524.72557 | A | 501.37592 | 251.19160 | 4 |
| 11 | 86.09643 | 1161.52794 | 581.26761 | L | 430.33880 | 215.67304 | 3 |
| 12 | 101.10733 | 1331.66986 | 666.33857 | K-Trimethyl | 317.25473 | 159.13100 | 2 |
| 13 | | | | K | 147.11281 | 74.06004 | 1 |

MS/MS fragmentation of AGKGSATLSMAYAGALFADACLK found in AL1G61640 (Lactate/malate dehydrogenase family protein)

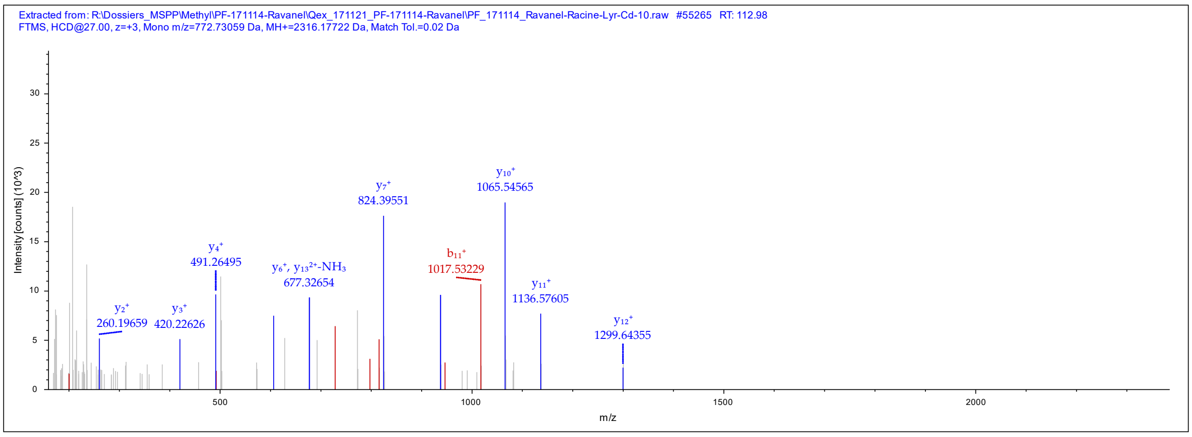


**Peptide Summary**

Sequence: AGKGSATLSMAYAGALFADACLK, K3-Trimethyl (42.04695 Da), C21-Carbamidomethyl (57.02146 Da)
Charge: +3, Monoisotopic m/z: 772.73059 Da (+1.48 mmu/+1.92 ppm), MH+: 2316.17722 Da, RT: 112.98 min,
Identified with: Mascot (v1.30); IonScore:50, Exp Value:1.3E-003, Ions matched by search engine: 9/256
Fragment match tolerance used for search: 20 mmu

**Fragment Matches**

Value Type: Theo. Mass [Da]

Ion Series | Neutral Losses | Precursor Ions

| #1 | Immonium | b+ | b2+ | b3+ | Seq. | y+ | y2+ | y3+ | #2 |
|---|---|---|---|---|---|---|---|---|---|
| 1 | 44.04948 | 72.04440 | 36.52584 | 24.68632 | A | | | | 23 |
| 2 | 30.03383 | 129.06587 | 65.03657 | 43.69347 | G | 2245.13566 | 1123.07147 | 749.05007 | 22 |
| 3 | 101.10733 | 299.20779 | 150.10753 | 100.40745 | K-Trimethyl | 2188.11419 | 1094.56073 | 730.04291 | 21 |
| 4 | 30.03383 | 356.22926 | 178.61827 | 119.41460 | G | 2017.97227 | 1009.48977 | 673.32894 | 20 |
| 5 | 60.04439 | 443.26129 | 222.13428 | 148.42528 | S | 1960.95080 | 980.97904 | 654.32178 | 19 |
| 6 | 44.04948 | 514.29841 | 257.65284 | 172.10432 | A | 1873.91877 | 937.46302 | 625.31111 | 18 |
| 7 | 74.06004 | 615.34609 | 308.17668 | 205.78688 | T | 1802.88165 | 901.94446 | 601.63207 | 17 |
| 8 | 86.09643 | 728.43016 | 364.71872 | 243.48157 | L | 1701.83397 | 851.42062 | 567.94951 | 16 |
| 9 | 60.04439 | 815.46219 | 408.23473 | 272.49225 | S | 1588.74990 | 794.87859 | 530.25482 | 15 |
| 10 | 104.05286 | 946.50269 | 473.75498 | 316.17241 | M | 1501.71787 | 751.36257 | 501.24414 | 14 |
| 11 | 44.04948 | 1017.53981 | 509.27354 | 339.85145 | A | 1370.67737 | 685.84232 | 457.56397 | 13 |
| 12 | 136.07568 | 1180.60313 | 590.80520 | 394.20589 | Y | 1299.64025 | 650.32376 | 433.88493 | 12 |
| 13 | 44.04948 | 1251.64025 | 626.32376 | 417.88493 | A | 1136.57693 | 568.79210 | 379.53049 | 11 |
| 14 | 30.03383 | 1308.66172 | 654.83450 | 436.89209 | G | 1065.53981 | 533.27354 | 355.85145 | 10 |
| 15 | 44.04948 | 1379.69884 | 690.35306 | 460.57113 | A | 1008.51834 | 504.76281 | 336.84430 | 9 |
| 16 | 86.09643 | 1492.78291 | 746.89509 | 498.26582 | L | 937.48122 | 469.24425 | 313.16526 | 8 |
| 17 | 120.08078 | 1639.85133 | 820.42930 | 547.28863 | F | 824.39715 | 412.70221 | 275.47057 | 7 |
| 18 | 44.04948 | 1710.88845 | 855.94786 | 570.96767 | A | 677.32873 | 339.16800 | 226.44776 | 6 |
| 19 | 88.03931 | 1825.91540 | 913.46134 | 609.30998 | D | 606.29161 | 303.64944 | 202.76872 | 5 |
| 20 | 44.04948 | 1896.95252 | 948.97990 | 632.98902 | A | 491.26466 | 246.13597 | 164.42640 | 4 |
| 21 | 76.02155 | 2056.98317 | 1028.99522 | 686.33257 | C-Carbami... | 420.22754 | 210.61741 | 140.74736 | 3 |
| 22 | 86.09643 | 2170.06724 | 1085.53726 | 724.02726 | L | 260.19688 | 130.60208 | 87.40381 | 2 |
| 23 | | | | | K | 147.11281 | 74.06004 | 49.70912 | 1 |

MS/MS fragmentation of TTQFCSGGK found in AT1G78820 (D-mannose binding lectin protein)

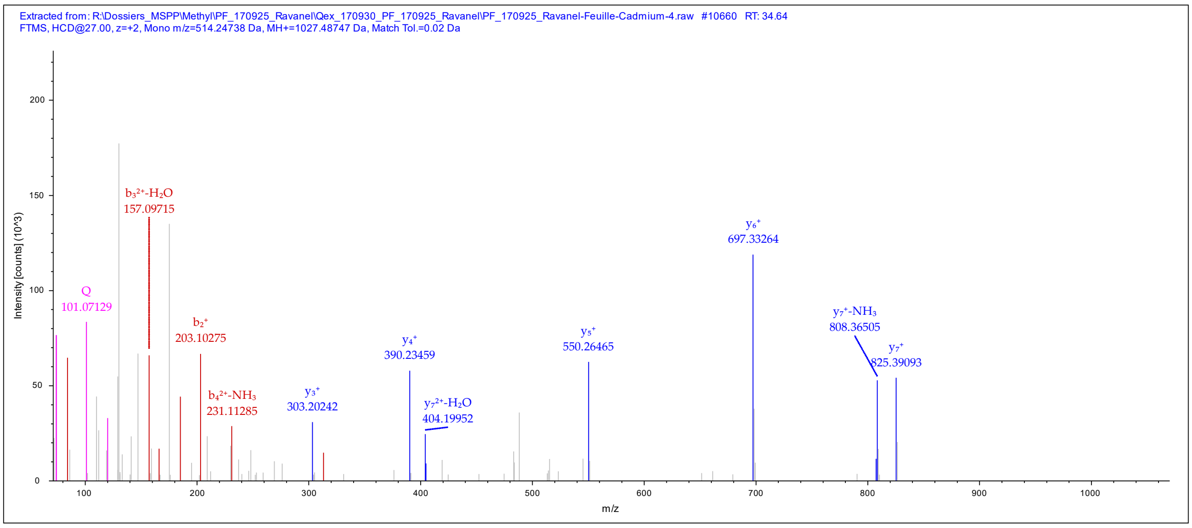


**Peptide Summary**

Sequence: TTQFCSGGK, C5-Carbamidomethyl (57.02146 Da), K9-Trimethyl (42.04695 Da)
Charge: +2, Monoisotopic m/z: 514.24738 Da (-0.14 mmu/-0.26 ppm), MH+: 1027.48747 Da, RT: 34.64 min,
Identified with: Mascot (v1.30); IonScore:44, Exp Value:6.1E-004, Ions matched by search engine: 5/88
Fragment match tolerance used for search: 20 mmu

**Fragment Matches**

Value Type: Theo. Mass [Da]

Ion Series | Neutral Losses | Precursor Ions

| #1 | Immonium | b⁺ | b²⁺ | Seq. | y⁺ | y²⁺ | #2 |
|---|---|---|---|---|---|---|---|
| 1 | 74.06004 | 102.05496 | 51.53112 | T | | | 9 |
| 2 | 74.06004 | 203.10264 | 102.05496 | T | 926.44007 | 463.72367 | 8 |
| 3 | 101.07094 | 331.16122 | 166.08425 | Q | 825.39239 | 413.19983 | 7 |
| 4 | 120.08078 | 478.22964 | 239.61846 | F | 697.33381 | 349.17054 | 6 |
| 5 | 76.02155 | 638.26029 | 319.63378 | C-Carbami... | 550.26539 | 275.63633 | 5 |
| 6 | 60.04439 | 725.29232 | 363.14980 | S | 390.23473 | 195.62100 | 4 |
| 7 | 30.03383 | 782.31379 | 391.66053 | G | 303.20270 | 152.10499 | 3 |
| 8 | 30.03383 | 839.33526 | 420.17127 | G | 246.18123 | 123.59425 | 2 |
| 9 | | | | K-Trimethyl | 189.15976 | 95.08352 | 1 |

MS/MS fragmentation of YTGEGESEEAKEGMFVK found in AT2G21330 (Fructose-bisphosphate aldolase 1)

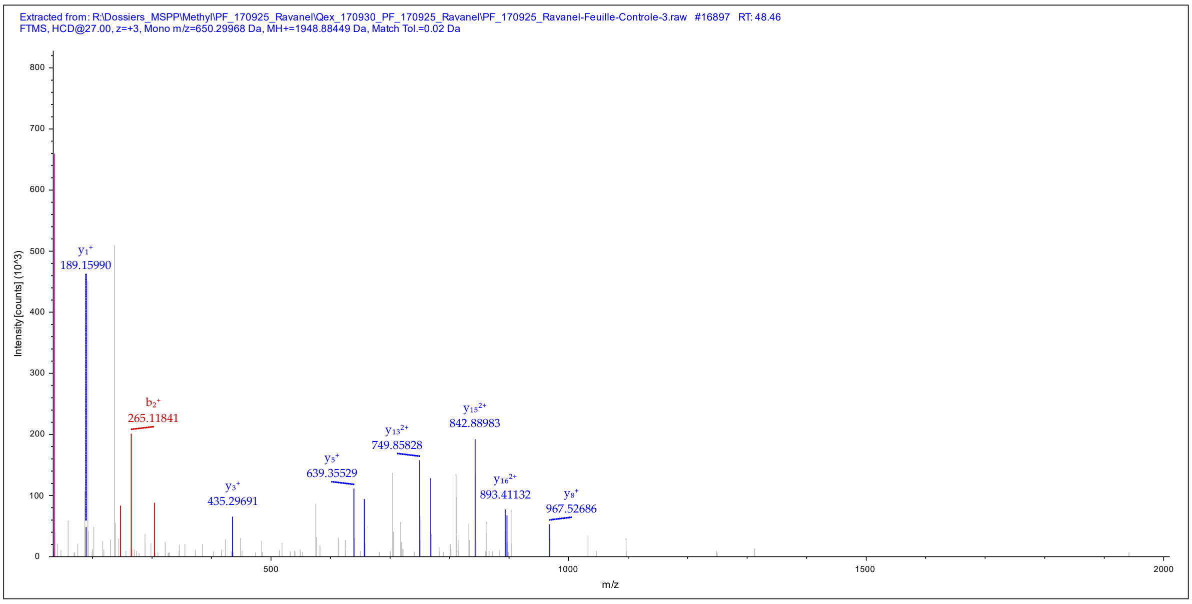


Peptide Summary

Sequence: YTGEGESEEAKEGMFVK, M14-Oxidation (15.99492 Da), K17-Trimethyl (42.04695 Da)
Charge: +3, Monoisotopic m/z: 650.29968 Da (-0.02 mmu/-0.03 ppm), MH+: 1948.88449 Da, RT: 48.46 min,
Identified with: Mascot (v1.30); IonScore:44, Exp Value:1.1E-003, Ions matched by search engine: 17/152
Fragment match tolerance used for search: 20 mmu

Fragment Matches

Value Type: Theo. Mass [Da]

Ion Series | Neutral Losses | Precursor Ions

| #1 | Immonium | $b^+$ | $b^{2+}$ | $b^{3+}$ | Seq. | $y^+$ | $y^{2+}$ | $y^{3+}$ | #2 |
|---|---|---|---|---|---|---|---|---|---|
| 1 | 136.07568 | 164.07060 | 82.53894 | 55.36172 | Y | | | | 17 |
| 2 | 74.06004 | 265.11828 | 133.06278 | 89.04428 | T | 1785.82123 | 893.41425 | 595.94526 | 16 |
| 3 | 30.03383 | 322.13975 | 161.57351 | 108.05143 | G | 1684.77355 | 842.89041 | 562.26270 | 15 |
| 4 | 102.05496 | 451.18235 | 226.09481 | 151.06563 | E | 1627.75208 | 814.37968 | 543.25554 | 14 |
| 5 | 30.03383 | 508.20382 | 254.60555 | 170.07279 | G | 1498.70948 | 749.85838 | 500.24134 | 13 |
| 6 | 102.05496 | 637.24642 | 319.12685 | 213.08699 | E | 1441.68801 | 721.34764 | 481.23419 | 12 |
| 7 | 60.04439 | 724.27845 | 362.64286 | 242.09767 | S | 1312.64541 | 656.82634 | 438.21999 | 11 |
| 8 | 102.05496 | 853.32105 | 427.16416 | 285.11187 | E | 1225.61338 | 613.31033 | 409.20931 | 10 |
| 9 | 102.05496 | 982.36365 | 491.68546 | 328.12607 | E | 1096.57078 | 548.78903 | 366.19511 | 9 |
| 10 | 44.04948 | 1053.40077 | 527.20402 | 351.80511 | A | 967.52818 | 484.26773 | 323.18091 | 8 |
| 11 | 101.10733 | 1181.49574 | 591.25151 | 394.50343 | K | 896.49106 | 448.74917 | 299.50187 | 7 |
| 12 | 102.05496 | 1310.53834 | 655.77281 | 437.51763 | E | 768.39609 | 384.70168 | 256.80355 | 6 |
| 13 | 30.03383 | 1367.55981 | 684.28354 | 456.52479 | G | 639.35349 | 320.18038 | 213.78935 | 5 |
| 14 | 104.05286 | 1514.59522 | 757.80125 | 505.53659 | M-Oxidation | 582.33202 | 291.66965 | 194.78219 | 4 |
| 15 | 120.08078 | 1661.66364 | 831.33546 | 554.55940 | F | 435.29660 | 218.15194 | 145.77038 | 3 |
| 16 | 72.08078 | 1760.73206 | 880.86967 | 587.58220 | V | 288.22818 | 144.61773 | 96.74758 | 2 |
| 17 | | | | | K-Trimethyl | 189.15976 | 95.08352 | 63.72477 | 1 |

MS/MS fragmentation of NFEGLDLGKMDEANDSGLASYVAGQIDR found in AL3G26800/AL3G26790 ((S)-2-hydroxy-acid oxidase)

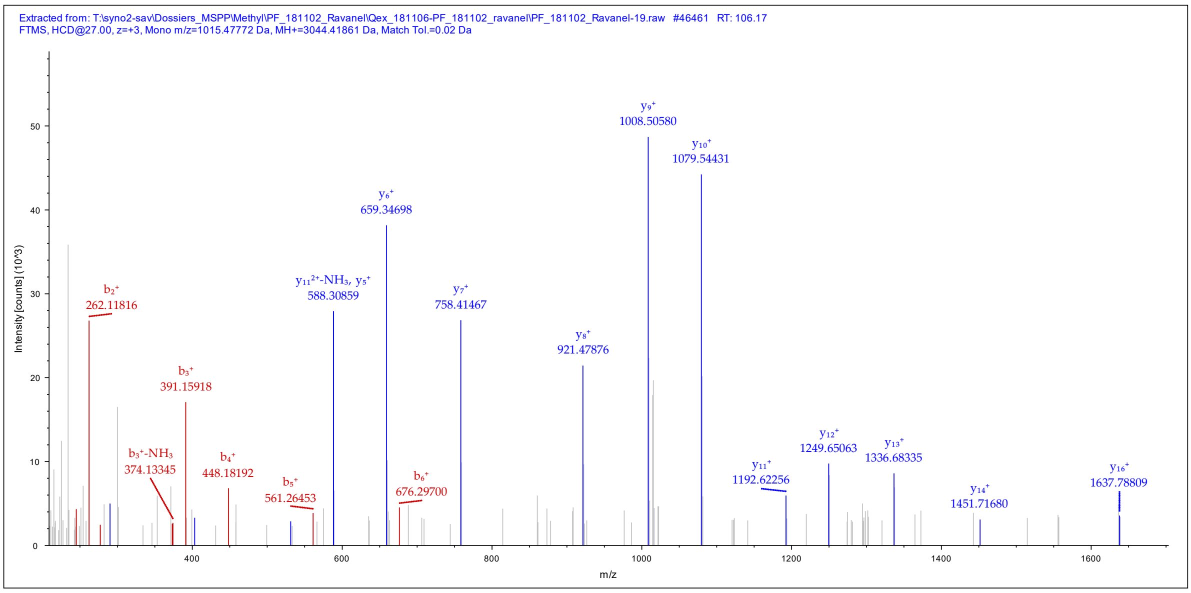


Sequence: NFEGLDLGKMDEANDSGLASYVAGQIDR, K9-Trimethyl (42.04695 Da), M10-Oxidation (15.99492 Da), N14-Deamidated (0.98402 Da)
Charge: +3, Monoisotopic m/z: 1015.47772 Da (+1.04 mmu/+1.03 ppm), MH+: 3044.41861 Da, RT: 106.17 min,
Identified with: Mascot (v1.30); IonScore:60, Exp Value:4.4E-004, Ions matched by search engine: 11/324
Fragment match tolerance used for search: 20 mmu

Fragment Matches

Value Type: Theo. Mass [Da]

Ion Series | Neutral Losses | Precursor Ions

| #1 | Immonium | b+ | b2+ | b3+ | Seq. | y+ | y2+ | y3+ | #2 |
|---|---|---|---|---|---|---|---|---|---|
| 1 | 87.05529 | 115.05021 | 58.02874 | 39.02159 | N | | | | 28 |
| 2 | 120.08078 | 262.11863 | 131.56295 | 88.04439 | F | 2930.37256 | 1465.68992 | 977.46237 | 27 |
| 3 | 102.05496 | 391.16123 | 196.08425 | 131.05859 | E | 2783.30414 | 1392.15571 | 928.43957 | 26 |
| 4 | 30.03383 | 448.18270 | 224.59499 | 150.06575 | G | 2654.26154 | 1327.63441 | 885.42537 | 25 |
| 5 | 86.09643 | 561.26677 | 281.13702 | 187.76044 | L | 2597.24007 | 1299.12367 | 866.41821 | 24 |
| 6 | 88.03931 | 676.29372 | 338.65050 | 226.10276 | D | 2484.15600 | 1242.58164 | 828.72352 | 23 |
| 7 | 86.09643 | 789.37779 | 395.19253 | 263.79745 | L | 2369.12905 | 1185.06816 | 790.38120 | 22 |
| 8 | 30.03383 | 846.39926 | 423.70327 | 282.80460 | G | 2256.04498 | 1128.52613 | 752.68651 | 21 |
| 9 | 101.10733 | 1016.54118 | 508.77423 | 339.51858 | K-Trimethyl | 2199.02351 | 1100.01539 | 733.67936 | 20 |
| 10 | 104.05286 | 1163.57659 | 582.29193 | 388.53038 | M-Oxidation | 2028.88159 | 1014.94443 | 676.96538 | 19 |
| 11 | 88.03931 | 1278.60354 | 639.80541 | 426.87270 | D | 1881.84618 | 941.42673 | 627.95358 | 18 |
| 12 | 102.05496 | 1407.64614 | 704.32671 | 469.88690 | E | 1766.81923 | 883.91325 | 589.61126 | 17 |
| 13 | 44.04948 | 1478.68326 | 739.84527 | 493.56594 | A | 1637.77663 | 819.39195 | 546.59706 | 16 |
| 14 | 87.05529 | 1593.71021 | 797.35874 | 531.90825 | N-Deamid... | 1566.73951 | 783.87339 | 522.91802 | 15 |
| 15 | 88.03931 | 1708.73716 | 854.87222 | 570.25057 | D | 1451.71256 | 726.35992 | 484.57570 | 14 |
| 16 | 60.04439 | 1795.76919 | 898.38823 | 599.26125 | S | 1336.68561 | 668.84644 | 446.23339 | 13 |
| 17 | 30.03383 | 1852.79066 | 926.89897 | 618.26840 | G | 1249.65358 | 625.33043 | 417.22271 | 12 |
| 18 | 86.09643 | 1965.87473 | 983.44100 | 655.96309 | L | 1192.63211 | 596.81969 | 398.21555 | 11 |
| 19 | 44.04948 | 2036.91185 | 1018.95956 | 679.64213 | A | 1079.54804 | 540.27766 | 360.52086 | 10 |
| 20 | 60.04439 | 2123.94388 | 1062.47558 | 708.65281 | S | 1008.51092 | 504.75910 | 336.84182 | 9 |
| 21 | 136.07568 | 2287.00720 | 1144.00724 | 763.00725 | Y | 921.47889 | 461.24308 | 307.83115 | 8 |
| 22 | 72.08078 | 2386.07562 | 1193.54145 | 796.03006 | V | 758.41557 | 379.71142 | 253.47671 | 7 |
| 23 | 44.04948 | 2457.11274 | 1229.06001 | 819.70910 | A | 659.34715 | 330.17721 | 220.45390 | 6 |
| 24 | 30.03383 | 2514.13421 | 1257.57074 | 838.71625 | G | 588.31003 | 294.65865 | 196.77486 | 5 |
| 25 | 101.07094 | 2642.19279 | 1321.60003 | 881.40245 | Q | 531.28856 | 266.14792 | 177.76770 | 4 |
| 26 | 86.09643 | 2755.27686 | 1378.14207 | 919.09714 | I | 403.22998 | 202.11863 | 135.08151 | 3 |
| 27 | 88.03931 | 2870.30381 | 1435.65554 | 957.43945 | D | 290.14591 | 145.57659 | 97.38682 | 2 |
| 28 | | | | | R | 175.11896 | 88.06312 | 59.04450 | 1 |

MS/MS fragmentation of ATSAGMKEQEAVNFLEK found in AL3G43370/AL7G44410 (20S proteasome alpha subunit)

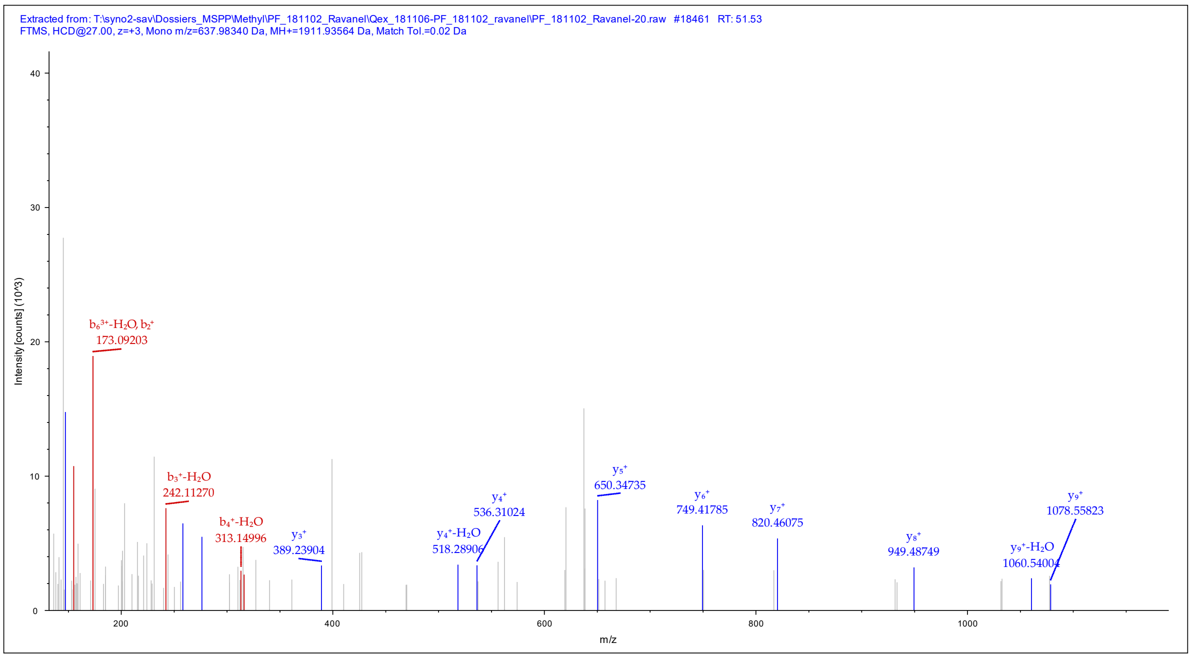


Sequence: ATSAGMKEQEAVNFLEK, M6-Oxidation (15.99492 Da), K7-Trimethyl (42.04695 Da), Q9-Deamidated (0.98402 Da)
Charge: +3, Monoisotopic m/z: 637.98340 Da (-0.43 mmu/-0.67 ppm), MH+: 1911.93564 Da, RT: 51.53 min,
Identified with: Mascot (v1.30); IonScore:37, Exp Value:6.4E-002, Ions matched by search engine: 9/168
Fragment match tolerance used for search: 20 mmu

Fragment Matches

Value Type: Theo. Mass [Da]

Ion Series | Neutral Losses | Precursor Ions

| #1 | Immonium | $b^+$ | $b^{2+}$ | $b^{3+}$ | Seq. | $y^+$ | $y^{2+}$ | $y^{3+}$ | #2 |
|---|---|---|---|---|---|---|---|---|---|
| 1 | 44.04948 | 72.04440 | 36.52584 | 24.68632 | A | | | | 17 |
| 2 | 74.06004 | 173.09208 | 87.04968 | 58.36888 | T | 1840.89980 | 920.95354 | 614.30479 | 16 |
| 3 | 60.04439 | 260.12411 | 130.56569 | 87.37955 | S | 1739.85212 | 870.42970 | 580.62223 | 15 |
| 4 | 44.04948 | 331.16123 | 166.08425 | 111.05859 | A | 1652.82009 | 826.91368 | 551.61155 | 14 |
| 5 | 30.03383 | 388.18270 | 194.59499 | 130.06575 | G | 1581.78297 | 791.39512 | 527.93251 | 13 |
| 6 | 104.05286 | 535.21811 | 268.11269 | 179.07755 | M-Oxidation | 1524.76150 | 762.88439 | 508.92535 | 12 |
| 7 | 101.10733 | 705.36003 | 353.18365 | 235.79153 | K-Trimethyl | 1377.72609 | 689.36668 | 459.91355 | 11 |
| 8 | 102.05496 | 834.40263 | 417.70495 | 278.80573 | E | 1207.58417 | 604.29572 | 403.19957 | 10 |
| 9 | 101.07094 | 963.44523 | 482.22625 | 321.81993 | Q-Deamid... | 1078.54157 | 539.77442 | 360.18537 | 9 |
| 10 | 102.05496 | 1092.48783 | 546.74755 | 364.83413 | E | 949.49897 | 475.25312 | 317.17117 | 8 |
| 11 | 44.04948 | 1163.52495 | 582.26611 | 388.51317 | A | 820.45637 | 410.73182 | 274.15697 | 7 |
| 12 | 72.08078 | 1262.59337 | 631.80032 | 421.53597 | V | 749.41925 | 375.21326 | 250.47793 | 6 |
| 13 | 87.05529 | 1376.63630 | 688.82179 | 459.55028 | N | 650.35083 | 325.67905 | 217.45513 | 5 |
| 14 | 120.08078 | 1523.70472 | 762.35600 | 508.57309 | F | 536.30790 | 268.65759 | 179.44082 | 4 |
| 15 | 86.09643 | 1636.78879 | 818.89803 | 546.26778 | L | 389.23948 | 195.12338 | 130.41801 | 3 |
| 16 | 102.05496 | 1765.83139 | 883.41933 | 589.28198 | E | 276.15541 | 138.58134 | 92.72332 | 2 |
| 17 | | | | | K | 147.11281 | 74.06004 | 49.70912 | 1 |

MS/MS fragmentation of KTPNSYMLQQFDNPANPK found in AL4G42620 (*O*-acetylserine(thiol)lyase)

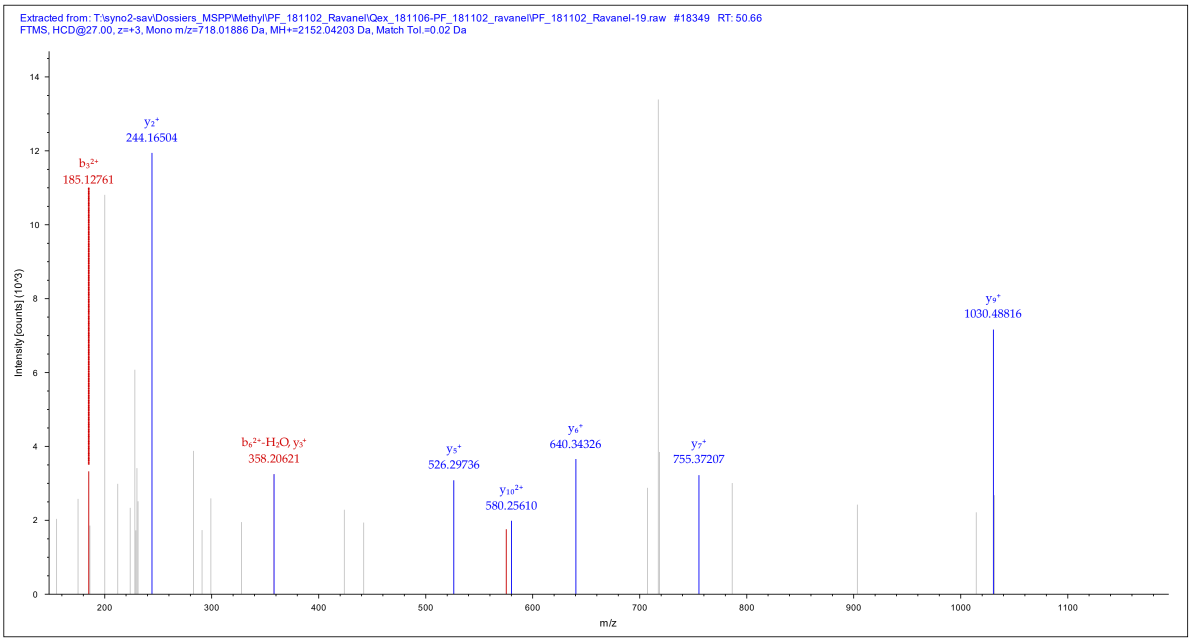


Sequence: KTPNSYMLQQFDNPANPK, K1-Trimethyl (42.04695 Da), M7-Oxidation (15.99492 Da), Q9-Deamidated (0.98402 Da)
Charge: +3, Monoisotopic m/z: 718.01886 Da (+1.34 mmu/+1.87 ppm), MH+: 2152.04203 Da, RT: 50.66 min,
Identified with: Mascot (v1.30); IonScore:34, Exp Value:1.6E-001, Ions matched by search engine: 6/204
Fragment match tolerance used for search: 20 mmu

Fragment Matches

Value Type: Theo. Mass [Da]

Ion Series | Neutral Losses | Precursor Ions

| #1 | Immonium | $b^+$ | $b^{2+}$ | $b^{3+}$ | Seq. | $y^+$ | $y^{2+}$ | $y^{3+}$ | #2 |
|---|---|---|---|---|---|---|---|---|---|
| 1 | 101.10733 | 171.14920 | 86.07824 | 57.72125 | K-Trimethyl | | | | 18 |
| 2 | 74.06004 | 272.19688 | 136.60208 | 91.40381 | T | 1981.89609 | 991.45168 | 661.30355 | 17 |
| 3 | 70.06513 | 369.24965 | 185.12846 | 123.75473 | P | 1880.84841 | 940.92784 | 627.62099 | 16 |
| 4 | 87.05529 | 483.29258 | 242.14993 | 161.76904 | N | 1783.79564 | 892.40146 | 595.27007 | 15 |
| 5 | 60.04439 | 570.32461 | 285.66594 | 190.77972 | S | 1669.75271 | 835.37999 | 557.25576 | 14 |
| 6 | 136.07568 | 733.38793 | 367.19760 | 245.13416 | Y | 1582.72068 | 791.86398 | 528.24508 | 13 |
| 7 | 104.05286 | 880.42334 | 440.71531 | 294.14596 | M-Oxidation | 1419.65736 | 710.33232 | 473.89064 | 12 |
| 8 | 86.09643 | 993.50741 | 497.25734 | 331.84065 | L | 1272.62195 | 636.81461 | 424.87883 | 11 |
| 9 | 101.07094 | 1122.55001 | 561.77864 | 374.85485 | Q-Deamid... | 1159.53788 | 580.27258 | 387.18414 | 10 |
| 10 | 101.07094 | 1250.60859 | 625.80793 | 417.54105 | Q | 1030.49528 | 515.75128 | 344.16994 | 9 |
| 11 | 120.08078 | 1397.67701 | 699.34214 | 466.56385 | F | 902.43670 | 451.72199 | 301.48375 | 8 |
| 12 | 88.03931 | 1512.70396 | 756.85562 | 504.90617 | D | 755.36828 | 378.18778 | 252.46094 | 7 |
| 13 | 87.05529 | 1626.74689 | 813.87708 | 542.92048 | N | 640.34133 | 320.67430 | 214.11863 | 6 |
| 14 | 70.06513 | 1723.79966 | 862.40347 | 575.27140 | P | 526.29840 | 263.65284 | 176.10432 | 5 |
| 15 | 44.04948 | 1794.83678 | 897.92203 | 598.95044 | A | 429.24563 | 215.12645 | 143.75339 | 4 |
| 16 | 87.05529 | 1908.87971 | 954.94349 | 636.96475 | N | 358.20851 | 179.60789 | 120.07435 | 3 |
| 17 | 70.06513 | 2005.93248 | 1003.46988 | 669.31568 | P | 244.16558 | 122.58643 | 82.06004 | 2 |
| 18 | | | | | K | 147.11281 | 74.06004 | 49.70912 | 1 |

MS/MS fragmentation of VENIVVIGHSACGGIKGLMSFPLDGNNSTDFIEDWVK found in AL6G25520/AL3G10670/AL3G10670.t3 (carbonic anhydrase)

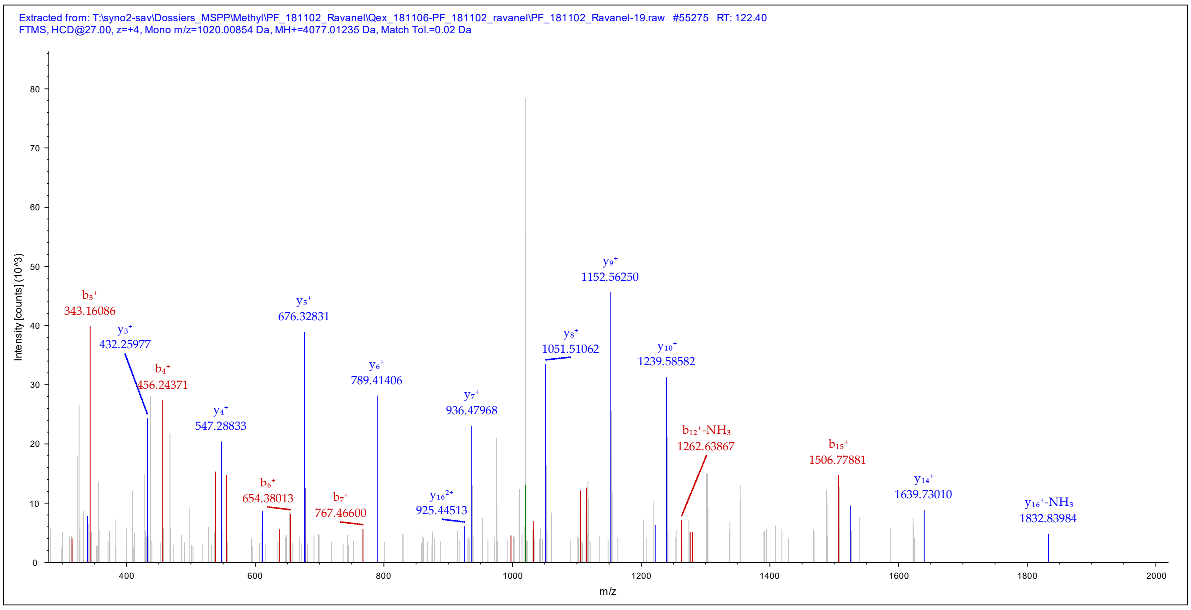


Sequence: VENIVVIGHSACGGIKGLMSFPLDGNNSTDFIEDWVK, C12-Carbamidomethyl (57.02146 Da), K16-Trimethyl (42.04695 Da), M19-Oxidation (15.99492 Da)
Charge: +4, Monoisotopic m/z: 1020.00854 Da (+0.63 mmu/+0.62 ppm), MH+: 4077.01235 Da, RT: 122.40 min,
Identified with: Mascot (v1.30); IonScore:36, Exp Value:2.3E-001, Ions matched by search engine: 7/424
Fragment match tolerance used for search: 20 mmu

Fragment Matches

Value Type: Theo. Mass [Da]

Ion Series | Neutral Losses | Precursor Ions

| #1 | Immonium | $b^+$ | $b^{2+}$ | $b^{3+}$ | $b^{4+}$ | Seq. | $y^+$ | $y^{2+}$ | $y^{3+}$ | $y^{4+}$ | #2 |
|---|---|---|---|---|---|---|---|---|---|---|---|
| 1 | 72.08078 | 100.07570 | 50.54149 | 34.03008 | 25.77438 | V | | | | | 37 |
| 2 | 102.05496 | 229.11830 | 115.06279 | 77.04428 | 58.03503 | E | 3977.94140 | 1989.47434 | 1326.65198 | 995.24081 | 36 |
| 3 | 87.05529 | 343.16123 | 172.08425 | 115.05859 | 86.54576 | N | 3848.89880 | 1924.95304 | 1283.63778 | 962.98016 | 35 |
| 4 | 86.09643 | 456.24530 | 228.62629 | 152.75328 | 114.81678 | I | 3734.85587 | 1867.93157 | 1245.62347 | 934.46943 | 34 |
| 5 | 72.08078 | 555.31372 | 278.16050 | 185.77609 | 139.58389 | V | 3621.77180 | 1811.38954 | 1207.92878 | 906.19841 | 33 |
| 6 | 72.08078 | 654.38214 | 327.69471 | 218.79890 | 164.35099 | V | 3522.70338 | 1761.85533 | 1174.90598 | 881.43130 | 32 |
| 7 | 86.09643 | 767.46621 | 384.23674 | 256.49359 | 192.62201 | I | 3423.63496 | 1712.32112 | 1141.88317 | 856.66420 | 31 |
| 8 | 30.03383 | 824.48768 | 412.74748 | 275.50074 | 206.87738 | G | 3310.55089 | 1655.77908 | 1104.18848 | 828.39318 | 30 |
| 9 | 110.07127 | 961.54659 | 481.27693 | 321.18705 | 241.14210 | H | 3253.52942 | 1627.26835 | 1085.18132 | 814.13781 | 29 |
| 10 | 60.04439 | 1048.57862 | 524.79295 | 350.19772 | 262.90011 | S | 3116.47051 | 1558.73889 | 1039.49502 | 779.87309 | 28 |
| 11 | 44.04948 | 1119.61574 | 560.31151 | 373.87676 | 280.65939 | A | 3029.43848 | 1515.22288 | 1010.48434 | 758.11508 | 27 |
| 12 | 76.02155 | 1279.64639 | 640.32683 | 427.22031 | 320.66706 | C-Carbami... | 2958.40136 | 1479.70432 | 986.80530 | 740.35580 | 26 |
| 13 | 30.03383 | 1336.66786 | 668.83757 | 446.22747 | 334.92242 | G | 2798.37071 | 1399.68899 | 933.46175 | 700.34813 | 25 |
| 14 | 30.03383 | 1393.68933 | 697.34830 | 465.23463 | 349.17779 | G | 2741.34924 | 1371.17826 | 914.45460 | 686.09277 | 24 |
| 15 | 86.09643 | 1506.77340 | 753.89034 | 502.92932 | 377.44881 | I | 2684.32777 | 1342.66752 | 895.44744 | 671.83740 | 23 |
| 16 | 101.10733 | 1676.91532 | 838.96130 | 559.64329 | 419.98429 | K-Trimethyl | 2571.24370 | 1286.12549 | 857.75275 | 643.56638 | 22 |
| 17 | 30.03383 | 1733.93679 | 867.47203 | 578.65045 | 434.23966 | G | 2401.10178 | 1201.05453 | 801.03878 | 601.03090 | 21 |
| 18 | 86.09643 | 1847.02086 | 924.01407 | 616.34514 | 462.51067 | L | 2344.08031 | 1172.54379 | 782.03162 | 586.77553 | 20 |
| 19 | 104.05286 | 1994.05628 | 997.53178 | 665.35694 | 499.26953 | M-Oxidation | 2230.99624 | 1116.00176 | 744.33693 | 558.50452 | 19 |
| 20 | 60.04439 | 2081.08831 | 1041.04779 | 694.36762 | 521.02753 | S | 2083.96082 | 1042.48405 | 695.32512 | 521.74566 | 18 |
| 21 | 120.08078 | 2228.15673 | 1114.58200 | 743.39043 | 557.79464 | F | 1996.92879 | 998.96803 | 666.31445 | 499.98766 | 17 |
| 22 | 70.06513 | 2325.20950 | 1163.10839 | 775.74135 | 582.05783 | P | 1849.86037 | 925.43382 | 617.29164 | 463.22055 | 16 |
| 23 | 86.09643 | 2438.29357 | 1219.65042 | 813.43604 | 610.32885 | L | 1752.80760 | 876.90744 | 584.94072 | 438.95736 | 15 |
| 24 | 88.03931 | 2553.32052 | 1277.16390 | 851.77836 | 639.08559 | D | 1639.72353 | 820.36540 | 547.24603 | 410.68634 | 14 |
| 25 | 30.03383 | 2610.34199 | 1305.67463 | 870.78551 | 653.34095 | G | 1524.69658 | 762.85193 | 508.90371 | 381.92960 | 13 |
| 26 | 87.05529 | 2724.38492 | 1362.69610 | 908.79982 | 681.85169 | N | 1467.67511 | 734.34119 | 489.89655 | 367.67424 | 12 |
| 27 | 87.05529 | 2838.42785 | 1419.71756 | 946.81413 | 710.36242 | N | 1353.63218 | 677.31973 | 451.88224 | 339.16350 | 11 |
| 28 | 60.04439 | 2925.45988 | 1463.23358 | 975.82481 | 732.12043 | S | 1239.58925 | 620.29826 | 413.86793 | 310.65277 | 10 |
| 29 | 74.06004 | 3026.50756 | 1513.75742 | 1009.50737 | 757.38235 | T | 1152.55722 | 576.78225 | 384.85726 | 288.89476 | 9 |
| 30 | 88.03931 | 3141.53451 | 1571.27089 | 1047.84969 | 786.13908 | D | 1051.50954 | 526.25841 | 351.17470 | 263.63284 | 8 |
| 31 | 120.08078 | 3288.60293 | 1644.80510 | 1096.87249 | 822.90619 | F | 936.48259 | 468.74493 | 312.83238 | 234.87611 | 7 |
| 32 | 86.09643 | 3401.68700 | 1701.34714 | 1134.56718 | 851.17721 | I | 789.41417 | 395.21072 | 263.80957 | 198.10900 | 6 |
| 33 | 102.05496 | 3530.72960 | 1765.86844 | 1177.58138 | 883.43786 | E | 676.33010 | 338.66869 | 226.11488 | 169.83798 | 5 |
| 34 | 88.03931 | 3645.75655 | 1823.38191 | 1215.92370 | 912.19459 | D | 547.28750 | 274.14739 | 183.10068 | 137.57733 | 4 |
| 35 | 159.09168 | 3831.83587 | 1916.42157 | 1277.95014 | 958.71442 | W | 432.26055 | 216.63391 | 144.75837 | 108.82060 | 3 |
| 36 | 72.08078 | 3930.90429 | 1965.95578 | 1310.97295 | 983.48153 | V | 246.18123 | 123.59425 | 82.73193 | 62.30077 | 2 |
| 37 | | | | | | K | 147.11281 | 74.06004 | 49.70912 | 37.53366 | 1 |

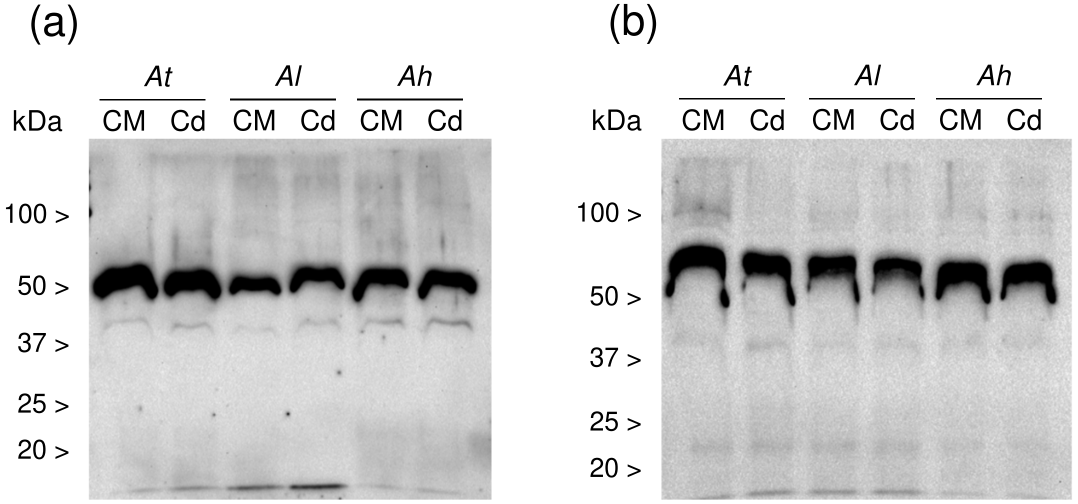


**Supplemental Figure S4:** Immunodetection of EEF1A in roots and leaves from Arabidopsis plants challenged with Cd. Plants grown in hydropony were maintained in control medium (CM) or challenged with 5 µM CdSO4 for 9 days. Soluble proteins (5 µg per lane) from root and leaf tissues were analyzed by western blot using antibodies to EEF1A (Agrisera, AS10 934). (a) - Leaf soluble proteins. (b) - Root soluble proteins. *At, A. thaliana*; *Al, A. lyrata*; *Ah, A. halleri* (AU population).

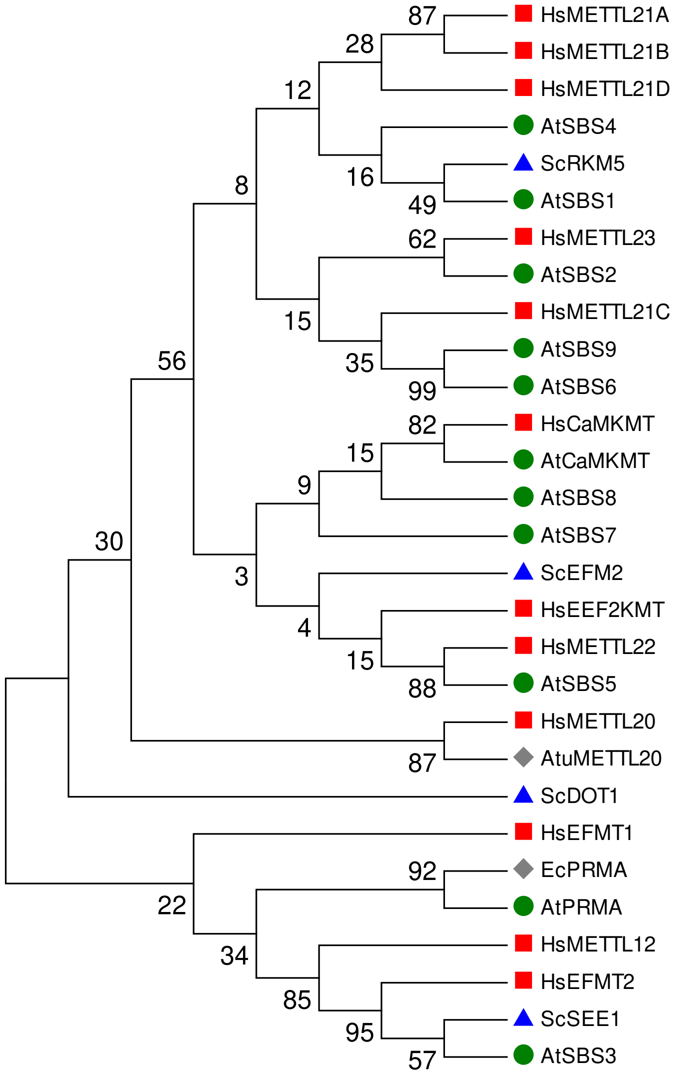


**Supplemental Figure S5:** Phylogenetic analysis of KMTs from the seven-beta strand (SBS) superfamily. The analysis included 29 amino acid sequences from *Homo sapiens* (Hs), *Saccharomyces cerevisiae* (Sc), *Escherichia coli* (Ec), *Agrobacterium tumefaciens* (Atu), and *A. thaliana* (At). The *A. thaliana* sequences were identified by BlastP search using SBS sequences from other species as queries. Sequences were aligned using ClustalW and the phylogenic tree was inferred by using the Maximum Likelihood method based on the Le-Gascuel model (Le & Gascuel, 2008). A discrete Gamma distribution was used to model evolutionary rate differences among sites (5 categories (+G, parameter = 2.7507). The rate variation model allowed for some sites to be evolutionarily invariable ([+I], 1.0515% sites). The consensus bootstrap tree is shown and branch support values (in % for 1,000 replicates) are indicated. Branches corresponding to partitions reproduced in less than 50% bootstrap replicates were collapsed. Analyses were done with the Mega 6.06 software (Tamura *et al*., 2013). UniProt accession numbers of query sequences: AtuMETTL20, A9CHJ5; EcPRMA, P0A8T1 ; HsCaMKMT, Q7Z624; HsEEF2KMT, Q96G04; HsEFMT1, Q8WVE0; HsEFMT2, Q5JPI9; HsMETTL12, A8MUP2; HsMETTL20, Q8IXQ9; HsMETTL21A, Q8WXB1;HsMETTL21B, Q96AZ1; HsMETTL21C, Q5VZV1; HsMETTL21D, Q9H867; HsMETTL22, Q9BUU2; HsMETTL23, Q86XA0; ScDOT1, Q04089; ScEFM2, P38347; ScRKM5, Q12367; ScSEE1, P40516. Accession numbers of the *A. thaliana* sequences are available in Supplemental Table S3.

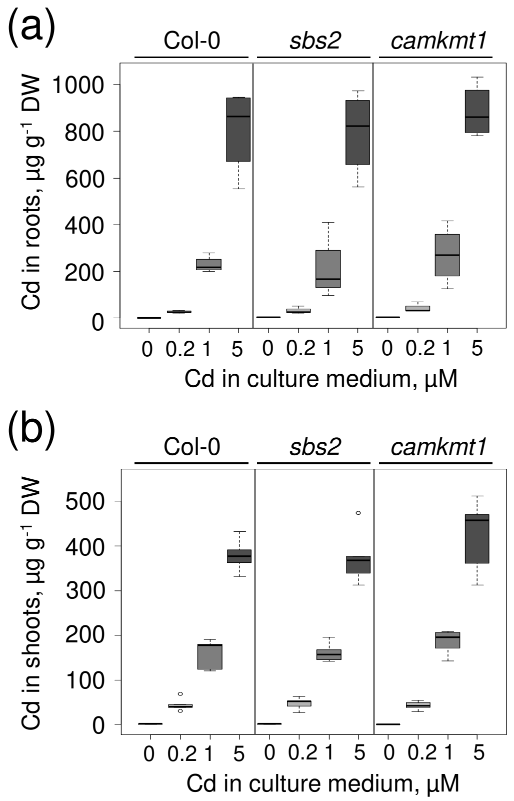


**Supplemental Figure S6**: Absorption and translocation of Cd in the *camkmt1* and *sbs2* mutants. Plants grown in hydropony were challenged with various concentrations of Cd (0, 0.2, 1 and 5 µM $CdSO_4$) for 7 days. Roots (a) and shoots (b) were dehydrated, mineralized in nitric acid, and $^{111}Cd$ was measured by ICP-MS. Each distribution represents n=5 seedlings from two biological replicates. No significant difference could be found between *camkmt1* or *sbs2* and Col-0 at any Cd concentration in the culture medium.

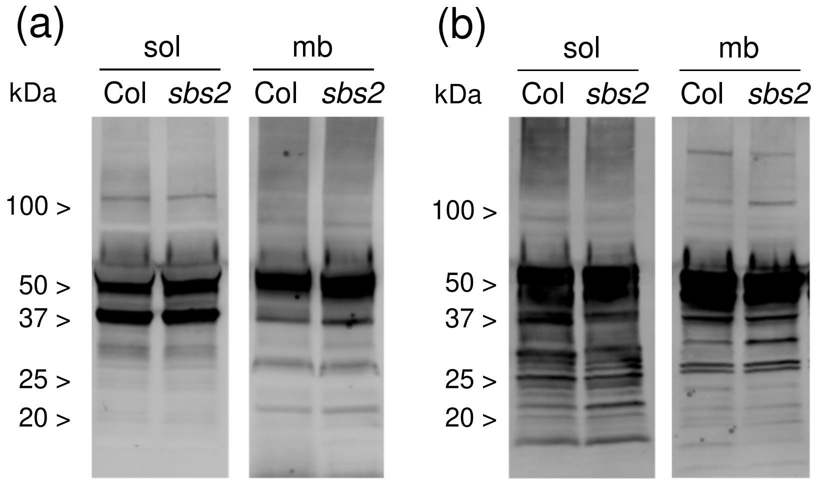


**Supplemental Figure S7:** Immunodetection of Lys-trimethylated proteins in roots and leaves from *sbs2* and Col-0 plants. Plants were grown in hydropony in control conditions for four weeks. Soluble and membrane proteins (30 µg per lane) from root and leaf tissues were analyzed by western blot using antibodies specific to trimethyl-Lys. (a) - Leaf soluble (sol) and membrane (mb) proteins. (b) - Root soluble and membrane proteins.
